# Autophagy regulates neuronal excitability by controlling cAMP/Protein Kinase A signaling

**DOI:** 10.1101/2022.02.11.480034

**Authors:** M. Overhoff, F. Tellkamp, S. Hess, J. Tutas, M. Tolve, M. Faerfers, L. Ickert, M. Mohammadi, E. De Bruyckere, E. Kallergi, A. Dell Vedove, V. Nikoletopoulou, B. Wirth, J. Isensee, T. Hucho, D. Puchkov, D. Isbrandt, M. Krüger, P. Kloppenburg, N.L. Kononenko

## Abstract

Autophagy provides nutrients during starvation and eliminates detrimental cellular components. However, accumulating evidence indicates that autophagy is not merely a housekeeping process. Here, we show that the protein AuTophaGy 5 (ATG5) functions in neurons to regulate the cAMP-dependent protein kinase A (PKA)-mediated phosphorylation of a synapse-confined proteome. This function of ATG5 is independent of bulk turnover of synaptic proteins and requires the targeting of PKA inhibitory R1 subunits to autophagosomes. Neuronal loss of ATG5 causes synaptic accumulation of PKA R1, which sequesters the PKA catalytic subunit and diminishes the cAMP/PKA-dependent phosphorylation of postsynaptic cytoskeletal proteins mediating AMPAR trafficking. Glutamatergic neurons-confined ATG5 deletion augments AMPAR-dependent excitatory neurotransmission and causes the appearance of spontaneous recurrent seizures in mice. Our findings identify a novel role of autophagy in regulating PKA signaling at glutamatergic synapses and suggest the PKA as a target for restoration of synaptic function in neurodegenerative conditions with autophagy dysfunction.

## Introduction

Autophagy is an evolutionary conserved catabolic process that serves to provide nutrients during starvation and to eliminate defective proteins and organelles via lysosomal degradation (*1*). The most prevalent form of autophagy is macroautophagy (hereafter autophagy), and during this process, portions of the cytoplasm are sequestered within double-membraned vesicles termed autophagosomes. These undergo subsequent maturation steps before being delivered to the lysosomes for degradation. Nutrient starvation is the most common signal for autophagy induction in various model organisms, including mice (*2*). Interestingly, although starvation has a debated role in the regulation of autophagic activity in cultured mammalian neurons (*3*, *4*), nutrient deprivation and/or mechanistic target of rapamycin complex 1 (mTORC1) inhibition have been shown to induce neuronal autophagy *in-vivo* (*5*, *6*), although this effect was more pronounced in the hypothalamus compared to other brain regions (*7*). In spite of this, the function of starvation-induced autophagy in the brain is not well investigated. For example, while starvation-induced autophagy promotes the survival of cancer cells, it is the selective role of autophagy in the clearance of toxic proteins and dysfunctional organelles that is suggested to play a major protective role in neurons (*8*).

Both selective and starvation-induced autophagy use the same core machinery to generate autophagosomes (*1*), which includes a component of ubiquitin (Ub)-like conjugation machinery ATG5. During the initial steps of autophagy, the Ub-like protein ATG12 conjugates with ATG5 (ATG12~ATG5), which further establishes a complex with ATG16L1. The ATG5 complex is then recruited to the autophagy initiation site, where it catalyzes the conjugation of LC3 to phosphatidylethanolamine, an event required for autophagosome elongation and closure. In contrast to starvation-induced autophagy that is thought to sequester random cytoplasmic cargo under nutrient-poor conditions, selective autophagy operates both under normal vegetative conditions and in response to various external stimuli and relies on autophagic receptors (e.g. p62, NBR1) that bridge the cargo and autophagosomal membrane (*9*). Degradative cargo of selective autophagy in neurons includes, among others, several pre- and postsynaptic proteins (*10*–*12*), axonal ER (*13*), and mitochondria (*14*). Such housekeeping function of neuronal autophagy is crucial for turnover of damaged and/or dysfunctional synaptic proteins and organelles and is required to maintain the pool of functional synaptic vesicles (SV) (*6*, *10*) and/or presynaptic Ca^2+^ homeostasis (*13*). Alterations in the function of autophagy proteins have been implicated in several neurodegenerative disorders characterized by protein inclusion formation, whereas knock-out (KO) of numerous ATG genes in neuronal progenitors and/or in cerebellar Purkinje neurons causes severe neurodegeneration, accompanied by dramatic accumulation of ubiquitinated protein aggregates (reviewed in (*15*)). However, accumulating evidence indicates that neuronal autophagy is not merely a housekeeping process. Neurons utilize components of the autophagy machinery to regulate neurotrophic signaling (*16*, *17*) and microtubule dynamics (*18*). Autophagy is also crucial for synapse assembly and pruning (*19*–*22*), and is required for synaptic plasticity and memory formation (*11*, *23*–*26*). How precisely autophagy contributes to synaptic function is currently under debate.

Here, we describe that autophagy can regulate synaptic function independently of its role in the turnover of damaged proteins and organelles. By using neuronal-confined mouse models of ATG5 deficiency in either excitatory or inhibitory neurons and a combination of SILAC and label-free quantitative proteomics, high-content microscopy, and live-imaging approaches, we show that autophagy functions in neurons to mediate the starvation-induced turnover of synapse-localized regulatory subunits R1α and R1β of protein kinase A (PKA). ATG5 loss in neurons is accompanied by sequestration of PKA catalytic subunit by its inhibitory R1α/β subunits, which diminishes PKA signaling and causes cAMP-dependent remodeling of cytoskeletal phosphoproteome confined to the postsynaptic density of excitatory synapses. Autophagy-deficient synapses are characterized by increased thickness of the postsynaptic density (PSD), impaired trafficking of AMPA receptors, and augmented excitatory neurotransmission, a PKA-dependent phenotype that likely results in the appearance of seizures in mice with glutamatergic forebrain-confined ATG5 deletion. Our findings identify a previously unknown role of starvation-induced autophagy in the regulation of PKA-dependent signaling in the brain, where it functions as a modulator of synaptic activity in cortical excitatory neurons.

## Results

### ATG5 deficiency in excitatory and inhibitory neurons results in differential excitability phenotype

To understand how precisely autophagy contributes to neuronal function, we capitalized on two mouse lines lacking crucial autophagy component ATG5 either in forebrain excitatory (*Atg5*flox:flox/*CamKII*α-Cre^tg^ KO mice, further defined as *Atg5*flox:*CamKII*α-Cre KO, Fig. 1A, fig. S1A-C) or inhibitory (*Atg5*flox:flox/*Slc32α1*-Cre^tg^ KO mice, further defined as *Atg5*flox:*Slc32a1-Cre* KO, Fig. 1B, fig. S1D) neurons. Similar to previously published *Atg5*flox:*CamKII*α-Cre KO mice (*18*), *Atg5*flox:*Slc32a1*-Cre KO mice were viable but revealed a cessation of weight gain starting at about one month of age when compared to their WT littermates (fig. S1E). Autophagy inhibition in *Atg5*flox:*Slc32a1-Cre* KO brains was reflected by significantly increased protein levels of autophagy receptor p62 (fig. S1F) and downregulated levels of LC3 at autophagosomal membranes (monitored via LC3II levels, fig. S1G). In line with the absence of neurodegeneration in mice lacking ATG5 in *CamKIIα*-dependent manner (*18*), we did not detect a loss of GABAergic neurons in the hippocampus and the entorhinal cortex of 3-month-old *Atg5*flox:*Slc32a1-Cre* KO mice (fig. S1H-K), a phenotype, which was consistent up to 10 months of age (fig. S1L).

**Figure 1.**
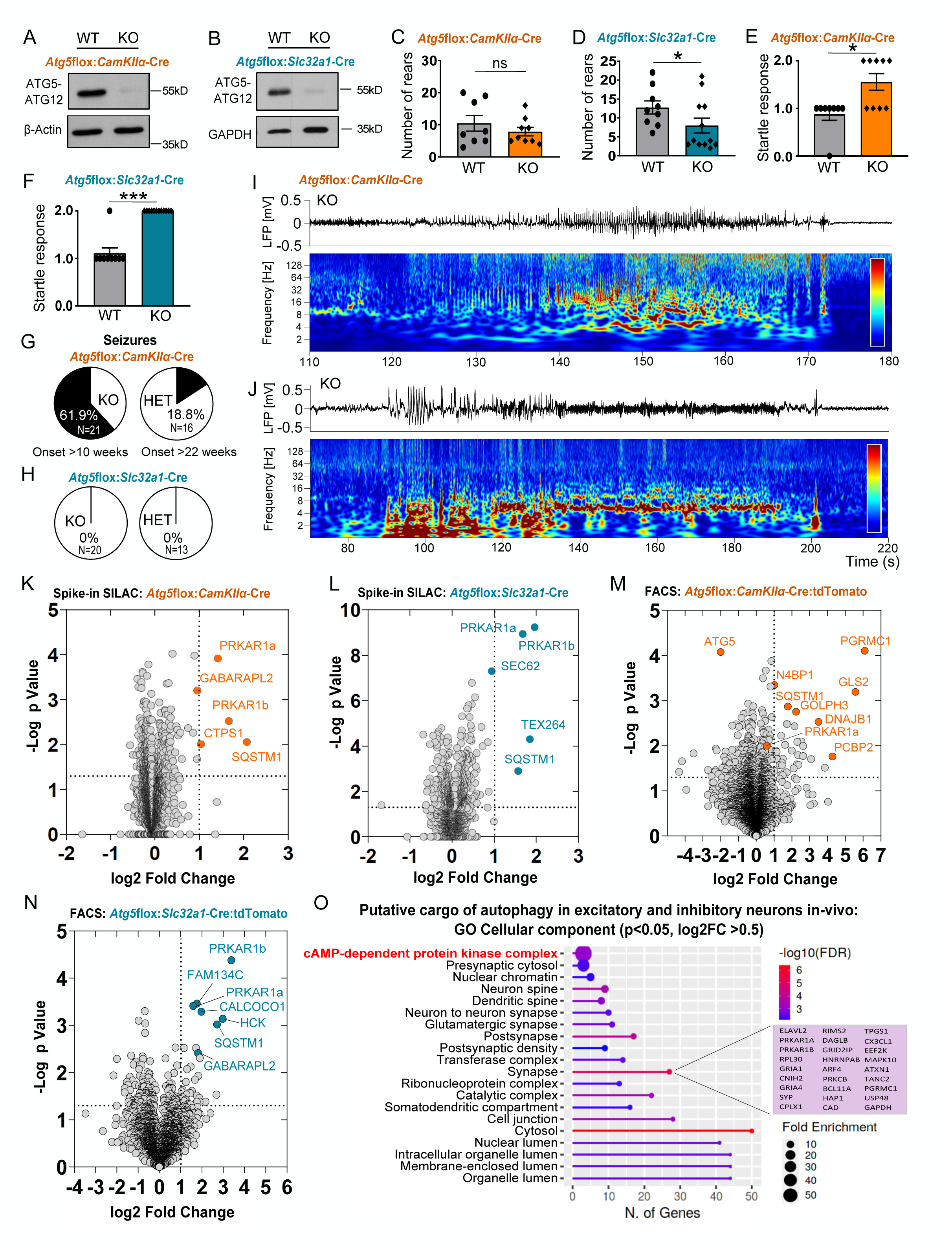
ATG5 deficiency in excitatory and inhibitory neurons results in differential neuronal excitability, despite similarities in proteome alterations. **(A, B)** Western Blot analysis of ATG5 protein levels in cortical brain lysates of 13-week-old *Atg5*flox:*CamKII*α-Cre WT/KO mice (A) or in striatal brain lysates of 13-week-old *Atg5*flox:*Slc32a1*-Cre WT/KO mice (B) (see also quantification in figs. S1A and S1D). **(C-D)** Rearing behavior analysis (number of standings on hind limbs) of *Atg5*flox:*CamKIIα*-Cre WT/KO mice ((C), WT: 10.50± 2.464, KO: 7.889± 1.338, n_WT_=8, n_KO_=9) and *Atg5*flox:*Slc32a1*-Cre WT/KO mice ((D), WT: 12.78± 1.730, KO: 6.818± 1.715, p=0.0262, n_WT_=9, n_KO_=11, two-tailed unpaired t-test). **(E-F)** Startle response analysis of *Atg5*flox:*CamKIIα*-Cre WT/KO mice ((E), WT: 0.8750±0.1250, KO: 1.556±0.1757; p= 0.0204, n_WT_=8, n_KO_= 9), as well as *Atg5*flox:*Slc32a1*-Cre WT/KO mice ((F) WT: 1.111±0.1111, KO: 2.000±0.000; p<0.0001, n_WT_=9, n_KO_=12, two-tailed unpaired Mann-Whitney test). **(G,H)** Behavioral seizure analysis in *Atg5*flox:*CamKII*α-Cre KO and heterozygous (HET) mice (G), as well as in *Atg5*flox:*Slc32a1*-Cre KO/HET mice (H). Mice were observed twice a week during 4 months after birth (the number animals and the age of seizure onset are indicated in the graph). No seizures were detected in WT mice set to 100%. **(I,J**) Examples of electrocorticogram (ECoG) traces (top) and corresponding wavelet spectrograms (bottom, warm colors indicate high power) from *Atg5*flox:*CamKII*α-Cre KO mice during a spontaneous generalized tonic-clonic seizure (I), or during a focal seizure associated with repetitive grooming and head-nodding (J). Representative example from n=4 recordings for each genotype. **(K-L)** Volcano plot of differentially expressed proteins in *Atg5*flox:*CamKIIα*-Cre KO cortical brain lysates (K) and *Atg5*flox:*Slc32a1*-Cre KO striatal brain lysates (L), analyzed using a SILAC-based proteomic approach (n_*Atg5*flox:*CamKII*α-Cre_=3, n_*Atg5*flox:*Slc32a1*-Cre_=5 mice per genotype) (see also fig. S1P). Orange- and blue-color-coded circles indicate all protein deregulated at p<0.05 and log2 fold change of >1. **(M-N)** Volcano plot of differentially expressed proteins in FACS-sorted *Atg5*flox:*CamKII*α-Cre:tdTomato (M) and *Atg5*flox:*Slc32a1*-Cre:tdTomato (N) KO neurons, analyzed using label-free proteomic analysis (n_*Atg5*flox:*CamKII*α-Cre:tdTomato_=4, n_*Atg5*flox:*Slc32a1*-Cre:tdTomato_= 5 mice per genotype). Orange- and blue-color-coded circles highlight highly deregulated proteins at p<0.05 and log2 fold change of >1. **(O)** ShinyGO v0.741-based gene ontology (GO) analysis of “cellular component” -enriched terms in the proteome (cut-off p<0.05, log2FC>0.5) of FACS sorted *Atg5*flox:*CamKII*α-Cre:tdTomato KO and *Atg5*flox:*Slc32a1*-Cre:tdTomato KO neurons (see also fig. S1R-U).

Even though ATG5 was equally dispensable for the survival of excitatory and inhibitory neurons, its loss resulted in differential effects on exploratory behavior in two lines conditionally lacking ATG5. Whereas *Atg5*flox:*CamKII*α-Cre KO mice were unaffected in their ability to explore the environment (Fig. 1C), *Atg5*flox:*Slc32a1-Cre* KO animals were less likely to exhibit a spontaneous rearing behavior in the open field when compared to their control littermates (i.e., either *Atg5*wt:wt/*Slc32a1*-Cre^tg^, *Atg5*flox:wt/*Slc32a1*-Cre^wt^ or *Atg5*flox:flox/*Slc32a1-Cre*^wt^, further defined as *Atg5*flox:*Slc32a1*-Cre WT, and/or *Atg5*flox:wt/*CamKII*α-Cre^wt^ and *Atg5*flox:flox/*CamKII*α-Cre^wt^, further defined as *Atg5*flox:*CamKIIα*-Cre WT, see also Methods section for the definition of WT and KO genotypes) (Fig. 1D). Overall locomotor activity was unaltered in both mouse lines (fig S1M). Since decreased rearing in mice is associated with increased anxiety, which can potentiate the startle reflex, we subjected both mouse lines to loud noise presentation and measured the startle response. Interestingly, although *Atg5*flox:*CamKII*α-Cre and *Atg5*flox:*Slc32a1-Cre* KO animals were both responding with potentiated startle reflex to stimulus presentation (Fig. 1E,F), only a significant percentage of *Atg5*flox:*CamKII*α-Cre KO mice reacted to the stimulus with the appearance of behavioral epileptic seizures (Fig. 1G-J, Suppl. Video 1). The seizures were present in 62% of homozygous 10-week-old *Atg5*flox/flox:*CamKII*α-Cre mice and in 19% of heterozygous 22-week-old *Atg5*flox/wt:*CamKIIα*-Cre mice. The presence of pathological electrocorticogram (ECoG) patterns, such as interictal spikes, spike trains, and spontaneous unprovoked electrographic generalized tonic-clonic seizures was documented in chronic radiotelemetry recordings from the somatomotor cortical region of 12-week-old *Atg5*flox/flox:*CamKII*α-Cre mice indicative of their increased neuronal network hyperexcitability (Fig. 1I,J, see also Data S1 for REM, SWS, and active awake periods in WT and KO animals). Interestingly, the behavioral seizures were also evident in a minor percentage of animals (<4%) with decreased levels of ATG5 in GABAergic neurons, but their onset was significantly delayed (>10-month-old) (fig. S1N). Epileptic seizures reflect hypersynchronous neuronal activity and can be either a result of intrinsic neuronal hyperexcitability and/or be a cause of neuronal network reorganization, e.g. due to neurodegeneration. The differential effect on the appearance of behavioral seizures in mice with autophagy-deficiency either in excitatory or inhibitory neurons taken together with the absence of neurodegeneration in these animals (*18*) suggests that ATG5 functions in the two classes of neurons to regulate their intrinsic excitability.

### PKA R1α/β levels are highly upregulated in autophagy-deficient excitatory and inhibitory neurons

What is the mechanism by which ATG5 regulates neuronal excitability? A recent study using mice lacking ATG5 in cortical (pallial) progenitors suggested that ATG5 regulates neurotransmission by selectively degrading the components of presynaptic tubular ER (*13*), whereas another study proposed that autophagy could regulate neuronal excitability by governing the degradation of damaged SV proteins (*10*). To reveal whether the changes in axonal ER components and/or bulk synaptic proteome are reflected in brains conditionally lacking ATG5 in either excitatory or inhibitory neurons, we conducted quantitative proteomics studies. First, we performed global proteomic profiling of conditional ATG5 KO mice using a spike-in SILAC mouse approach, where heavy SILAC reference brain tissue (*27*) was spiked into brain lysates obtained from either 12-13-week-old *Atg5*flox:*CamKII*α-Cre and *Atg5*flox:*Slc32a1*-Cre KO mice or their control littermates (fig. S1O). In agreement with Kuijpers et al (*13*) we found no major alterations in bulk levels of SV proteins in brains lacking ATG5 in either excitatory or inhibitory neurons (see also fig. S2S,T). In fact, except for known autophagy receptors such as GABARAPL2 and p62 (SQSTM1), only a few proteins were highly upregulated at the significance levels of p<0.05 and log2 fold change of >1 (Fig. 1K), and these changes were only slightly intensified at log2 fold change of >0.5 (fig. S1P). Among highly upregulated proteins in *Atg5*flox:*Slc32a1*-Cre KO brains we detected recently identified reticulophagy receptors TEX264 and SEC62, implicating reticulophagy in maintaining the physiology of forebrain inhibitory neurons (Fig. 1L). Intriguingly, in both types of autophagy-deficient neurons, regulatory subunits of PKA holoenzyme R1α (PRKAR1α) and R1β (PRKAR1b) were the most significantly upregulated proteins. These findings provided the first line of evidence that autophagy may regulate PKA signaling in excitatory and inhibitory neurons.

To understand whether changes in levels of PKA R1α and R1β represent a cell-autonomous role of autophagy in neurons, we developed a fluorescence-activated cell sorting (FACS)–based strategy to perform label-free proteomic analysis of WT and ATG5 KO neurons isolated from brains of 3-4-week-old *Atg5*flox:*CamKII*α-Cre and *Atg5*flox:*Slc32a1*-Cre KO mice (the age when the contribution of debris to cell sorting is relatively low) and their control littermates, carrying the tdTomato allele (fig. S1Q). Proteomic analysis of FACS-sorted neurons identified significantly more dysregulated proteins in both classes of autophagy-deficient neurons compared to the SILAC-based approach. We observed a significant decrease in ATG5 levels in FACS-sorted *Atg5*flox:*CamKII*α-Cre KO neurons and its complete loss in neurons isolated from *Atg5*flox:*Slc32a1*-Cre KO mice (Fig. 1M,N, Suppl. Table 1), reflecting the reliability of our FACS-sorting protocol. Protein classification analysis revealed that most of the proteins enriched in the KO neurons *in-vivo* belonged to metabolite interconversion enzymes, nucleic acid metabolism proteins and/or protein-modifying enzymes (fig. S1R). In agreement with the data from SILAC-based proteomics, PKA R1α and R1β subunits were among the significantly upregulated proteins in FACS-sorted autophagy-deficient neurons. This upregulation was less robust in autophagy-deficient excitatory neurons, likely reflecting their confinement to peripheral synaptic processes (see also Fig. 2), which cannot be detected in the FACS-based proteomics approach. We next subjected proteins identified in FACS-based proteomics in both mouse lines to a Gene Ontology (GO) term enrichment analysis aiming at identifying cellular components and molecular functions, which are cell-autonomously regulated by autophagy in both classes of neurons. GO term analysis of proteome from FACS-sorted autophagy-deficient excitatory and inhibitory neurons revealed an enrichment of several pre- and postsynaptic proteins (e.g., RIMS2, CPLX1, SYP, CNIH2, TANC2, DAGLB) (Fig. 1O, fig. S1R, see also Suppl. Table 1). These data agree with several recent studies and highlight the cell-autonomous role for autophagy in maintaining the homeostasis of synaptic proteins in the soma of excitatory and inhibitory neurons *in-vivo* (*7*, *11*, *25*, *28*). In line with recent work (*28*), we were not able to detect enrichment and/or upregulation of mitophagy regulators such as PINK1 or Parkin. This may either reflect the absence of mitochondrial clearance by autophagy in the cell soma of neurons or suggest the existence of ATG5-independent pathways for neuronal mitophagy.

**Figure 2.**
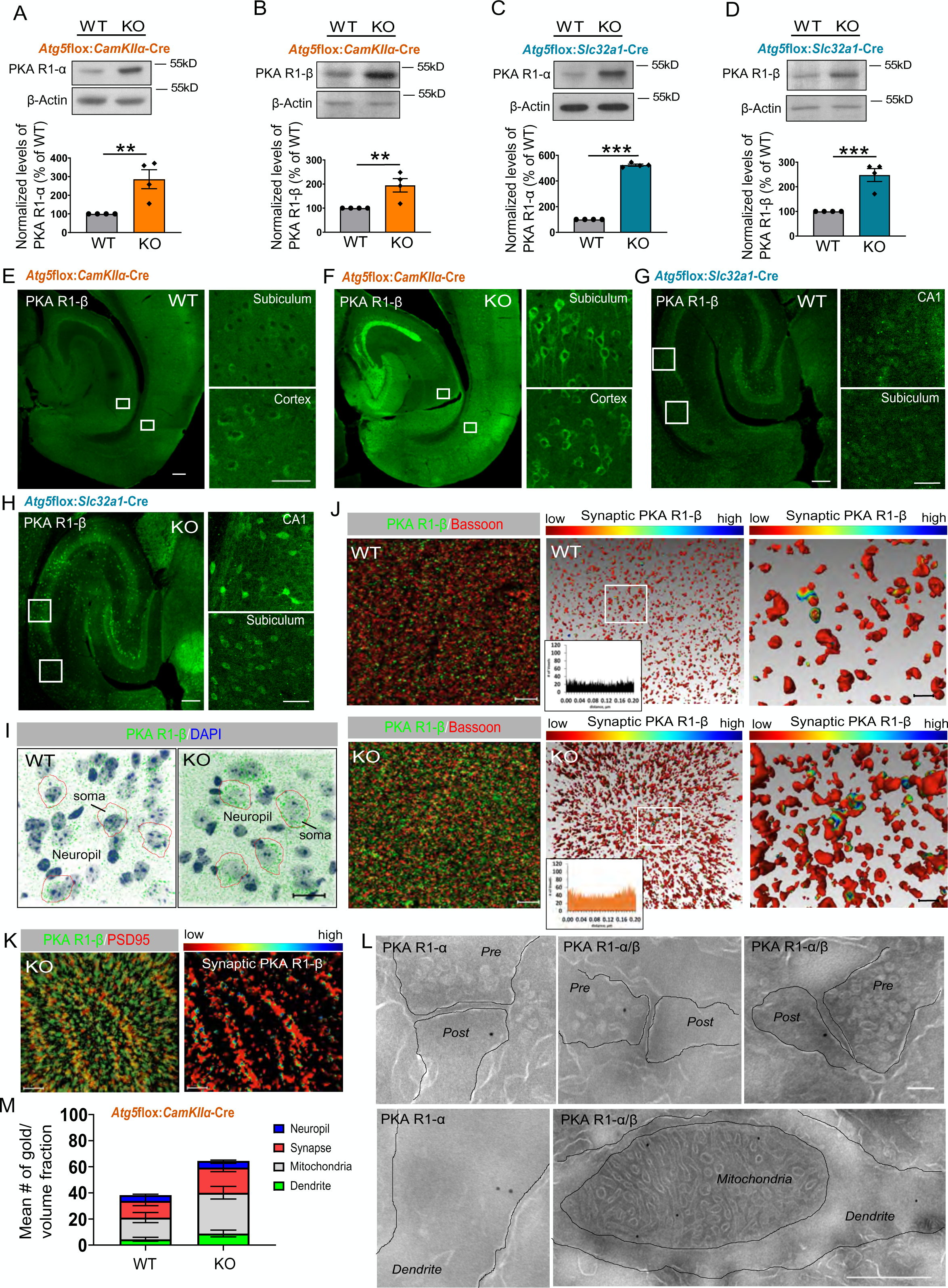
PKA regulatory subunits 1α and 1β are robustly upregulated in autophagy-deficient brains. **(A, B)** PKA R1-α (A) and PKA R1-β (B) protein levels are significantly increased in cortical brain lysates from 13-week-old *Atg5*flox:*CamKII*α-Cre KO mice compared to the WT set to 100% (KO_PKA R1-α_: 286.1± 50.53%, p=0.005, KO_PKA R1-β_: 194.7± 27.82%, p=0.007, one-sample t-test). n=4 for each genotype. **(C, D)** PKA R1-α (C) and PKA R1-β (D) protein levels are significantly increased in striatal lysates of *Atg5*flox:*Slc32a1* KO mice compared to the WT set to 100% (KO_PKA R1-α_: 524.1± 8.35%, p<0.0001, KO_PKA R1-β_: 247.9± 26.26%, p=0.0007, one-sample t-test). n=4 for each genotype. **(E-H)** Immunohistochemical analysis of PKA R1-β distribution in horizontal brain sections from *Atg5*flox:*CamKIIα*-Cre WT/KO (E, F) and *Atg5*flox:*Slc32a1*-Cre WT/KO (G,H) mice. Scale bars: 200 μM in large panels, 50 μM in small panels. **(I)** AMIRA-based 3Dvisualization of PKA R1-β distribution in *Atg5*flox:*CamKIIα*-Cre WT/KO cortex. Scale bar 20 μm. **(J)** Immunohistochemical profile (most left) and the 3D surface rendering of Amira 3D reconstructions (right) of PKA R1-β and bassoon colocalization in *Atg5*flox:*CamKIIα*-Cre WT and KO cortex. In the 3D analysis, the PKA R1-β neuropil staining is represented by the 3D reconstruction of its surface in red and PKA R1-β/Bassoon contacts are color coded, with the cold to warm colors spreading from 0- to 250-nm distance between the surface of either Bassoon-positive synapses and/or PKA R1-β-positive neuropil (see color-coded horizontal bar for the distance definition). Insert shows the histogram of the number of Bassoon-labeled voxels (3D pixels) found within 250 nm of the PKA R1-β neuropil staining. Scale bar 5 μm in dual-channel, 1 μm in 3D-reconstruction zoom. **(K)** Immunohistochemical profile (most left) and the 3D surface rendering of Amira 3D reconstruction (right) of PKA R1-α colocalization with the PSD95 in *Atg5*flox:*CamKIIα*-Cre KO CA1 area of the hippocampus. Scale bar 2 μm. **(L,M)** Electron microscopy-based analysis of immunogold-labeled PKA R1-α/β on Tokuyasu cryosections of the CA1 neuropil area of the hippocampus of Atg5flox:*CamKIIα*-Cre KO mice. PKA R1-α/β is enriched at synapses (WT: 12.762± 3.625, KO: 19.323± 3.172), dendritic compartment (WT: 4.462± 1.539, KO: 8.869± 2.559) and mitochondria (WT: 16.618± 3.865, KO: 31.291± 4.829) of KO mice. Values in (M) are the mean number of gold particles per volume fraction in n=21 WT and n=22 (KO) images analyzed with a grid size of 500k and obtained from two mice for each genotype. No immunogold labelling was detected in samples where the PKA R1-α/β antibody were omitted (negative control). Scale bar: upper row 50 nm, 500 nm lower picture.

In agreement with robust upregulation of PKA R1α and R1β subunits in ATG5-deficient neurons, GO term enrichment analysis revealed “cAMP-dependent protein kinase complex” (Fig. 1O) and PKA-associated pathways (fig. S1S-U) as top-ranked GO terms. PKA is a key kinase involved in the regulation of multiple intracellular signal transduction pathways. PKA holoenzyme is a heterotetramer consisting of a dimer of two regulatory (R) subunits, with each binding a catalytic subunit. PKA signaling is activated when two molecules of cyclic adenosine monophosphate (cAMP) bind each regulatory subunit of the PKA heterotetramer, leading to its dissociation and releasing of the catalytic subunits (*29*). Catalytic subunits then become active to phosphorylate protein substrates at serine or threonine residues, resulting in changes in cell function. Importantly, in the brain, the PKA function is essential in the regulation of neuronal and network activities, where PKA takes a major role in controlling AMPA and NMDA glutamate receptor and sodium channels activity at synapses (*30*). This established role of PKA in the regulation of synaptic function together with our findings indicating a strong upregulation in the levels of inhibitory R1α and R1β subunits of PKA complex in autophagy-deficient neurons suggests autophagy to control neuronal excitation by regulating the PKA signaling.

### Neuronal autophagy stabilizes synaptic levels of PKA type 1 inhibitory subunits

The data described above suggest that autophagy regulates the protein level of inhibitory subunits of the PKA holoenzyme R1α and R1β in excitatory and inhibitory neurons. To further test this hypothesis, we first validated the results from the proteomics studies by Western Blotting. As illustrated in Fig. 2A-D the levels of R1α and R1β were robustly upregulated in the cortex and hippocampus of 12-13-week-old *Atg5*flox:*CamKII*α-Cre KO mice (Fig. 2A,B, fig. S2A,B), as well as in the striatum, cortex, and cerebellum of 12-13-week-old *Atg5*flox:*Slc32a1*-Cre KO mice (Fig. 2C,D, fig. S2C,D), indicating that autophagy regulates levels of inhibitory subunits of PKA holoenzyme across different brain regions. In mammals, four inhibitory R subunit (R1α, R1β, R2α, and R2β) have been identified, comprising the type 1 or the type 2 PKA isozymes, respectively (*29*). Since all regulatory subunits are expressed in the brain, we investigated the levels of R2α and R2β in brain lysates of autophagy-deficient mice. The levels of R2α, and R2β were neither altered in brains of *Atg5*flox:*CamKII*α-Cre KO mice (fig. S2E,F), nor in brains of *Atg5*flox:*Slc32a1*-Cre KO mice (Fig. S2G). Furthermore, ATG5 was also dispensable for the regulation of bulk levels of catalytic PKA-Cα subunit in neurons *in-vivo* (fig S2H-L). Additionally, the levels of R1α and R1β, but not PKA-Cα, were also dysregulated in the *in-vitro* cortico-hippocampal culture system, where the deletion of ATG5 (fig. S2M-O) was driven by a tamoxifen-dependent activation of the *CAG*-Cre promoter (Cre^Tmx^) (*18*). This effect was not specific to ATG5, since the deletion of ATG16L1, another component of the ATG12~ATG5-ATG16L1 conjugate, crucial for LC3 lipidation (*18*), was sufficient to stabilize the levels of R1α and R1β in tamoxifen-treated primary neurons isolated from *Atg16L1* flox:*CAG*-Cre^Tmx^ mice (fig. S2P), as well as in cortical brain lysates of 12-13-week-old *Atg16L1*flox: *Slc32a1*-Cre KO mice (fig. S2Q). Taken together, our data indicate that neuronal autophagy functions *in-vivo* and *in-vitro* to selectively regulate the levels of type 1 PKA inhibitory subunits.

Next, we determined the exact cellular and sub-cellular localization of R1α and R1β in neurons. To this aim, we first performed immunohistochemical imaging of R1α and R1β in brains of *Atg5*flox:*CamKII*α-Cre and *Atg5*flox:*Slc32a1*-Cre WT and KO mice, using R1β antibodies, which recognize R1α in addition, when used for immunohistochemistry (*31*). As shown in Fig. 2E-H, R1β was prominently expressed in the hippocampus and the cortex of control mice, and this expression pattern was intensified by the ATG5 deletion. To understand the localization of R1β at the cellular level, we employed the 3D-based reconstruction approach, which revealed strong R1β accumulation in neuronal soma (Fig. 2I), and its confinement to Bassoon-positive cortical synapses (Fig. 2J) in *Atg5*flox:*CamKIIα*-Cre KO brains. In agreement with dendritic localization of R1β in wildtype mice (*31*), we found that in autophagy-deficient excitatory neurons, R1β was enriched in dendrites and co-localized with postsynaptic density marker PSD95 (Fig. 2K, fig. S2R). Although murine R1α and R1β share 82% sequence identity, they are not functionally redundant, which can be partially explained by their differential subcellular localization, where R1α is cytosolic, whereas R1β is selectively enriched in the mitochondria (*31*). To understand the subcellular localization of R1α and R1β in autophagy-deficient neurons, we performed immunogold-based labeling on Tokuyasu cryosections of the CA1 neuropil area of the hippocampus in *Atg5*flox:*CamKIIα*-Cre WT and KO mice. In agreement with (*31*), our immunogold analysis revealed a relatively sparse cytoplasmic labeling for R1α in wildtype neurons and its accumulation in the cytosol of dendrites and at the PSD of neurons, lacking ATG5 (Fig. 2L). However, due to the sparse content of R1α antibody labeling in the WT, we were not able to quantify these differences. In contrast, the antibody recognizing both R1α and R1β subunits was efficient in labeling R1α and R1β subcellular distribution in both WT and ATG5-deficient excitatory neurons (the specificity of immunogold labeling was confirmed by the complete absence of gold particles in the “negative control” samples). Intriguingly, we found that autophagy deficiency causes enrichment of type 1 inhibitory PKA subunits at synapses and mitochondria (Fig. 2M). These data, taken together with the fact that no alterations in bulk levels of SV proteins were identified upon neuronal autophagy deficiency in SILAC-based proteomics (fig. S2S,T), suggest that autophagy might contribute to neuronal function via regulating the PKA-dependent signaling at synapses and/mitochondria. Since recently, autophagy-mediated degradation of R1α has been shown to regulate the mitochondrial metabolism in response to glucose starvation in non-neuronal cells (*32*), we focused our further studies on the autophagy-dependent PKA function at synapses.

### PKA type 1 subunits are degraded by starvation-induced autophagy

The observed changes in the levels of PKA R1α and R1β could be attributed either to its accumulation as autophagy substrate or to related secondary effects due to enhanced or decreased activation of signaling pathways. To understand how precisely autophagy regulates levels of R1α and R1β in neuronal cells, we first tested the effect of starvation and/or mTORC1 inhibition, two conditions known to induce autophagy, on the levels of PKA R1α in NSC34 cells. We found that PKA R1α levels were significantly decreased in neuronal cells subjected to amino acid and serum starvation using media containing 5mM glucose (Fig. 3A,B, fig. S3A). In contrast to starvation, the decrease in PKA R1α was not evident upon treatment of NSC34 cells with an ATP-competitive inhibitor of mTORC1, Torin1 (Fig. 3A, fig. S3A), although the levels of pS6 Ser-235/236, used as a readout of mTOR activity, were equally decreased (fig. S3B,C), suggesting that starvation decreases R1α levels independently of mTORC1 activity. Starvation-dependent decrease in PKA R1α was rescued by acute suppression of autophagy using a specific V-ATPase inhibitor Bafilomycin A1 (Fig. 3C). PKA R1α levels were also stabilized in NSC34 cells cultured in amino acid- and serum-rich media and treated with Bafilomycin A1 (Fig. 3D,E), indicating that autophagy can directly be responsible for the degradation of the R1α. To prove that R1α and RIβ inhibitory subunits of PKA holoenzyme are autophagic cargo of neuronal autophagosome, we tested the levels of both subunits in autophagosomes isolated from the mouse brain. Both R1α and R1β were localized to purified brain autophagosomes (Fig. 3F). The role of starvation in the activation of neuronal PKA signaling was also demonstrated by increased phosphorylation of PKA substrates in primary WT neurons deprived of amino acids and serum, a phenotype that could be prevented by applcation of the ATP-site PKA inhibitor H89 (Fig. 3G,H). This may reflect the liberation of PKA catalytic subunits due to facilitated autophagy-dependent degradation of PKA R1 subunits in neurons.

**Figure 3.**
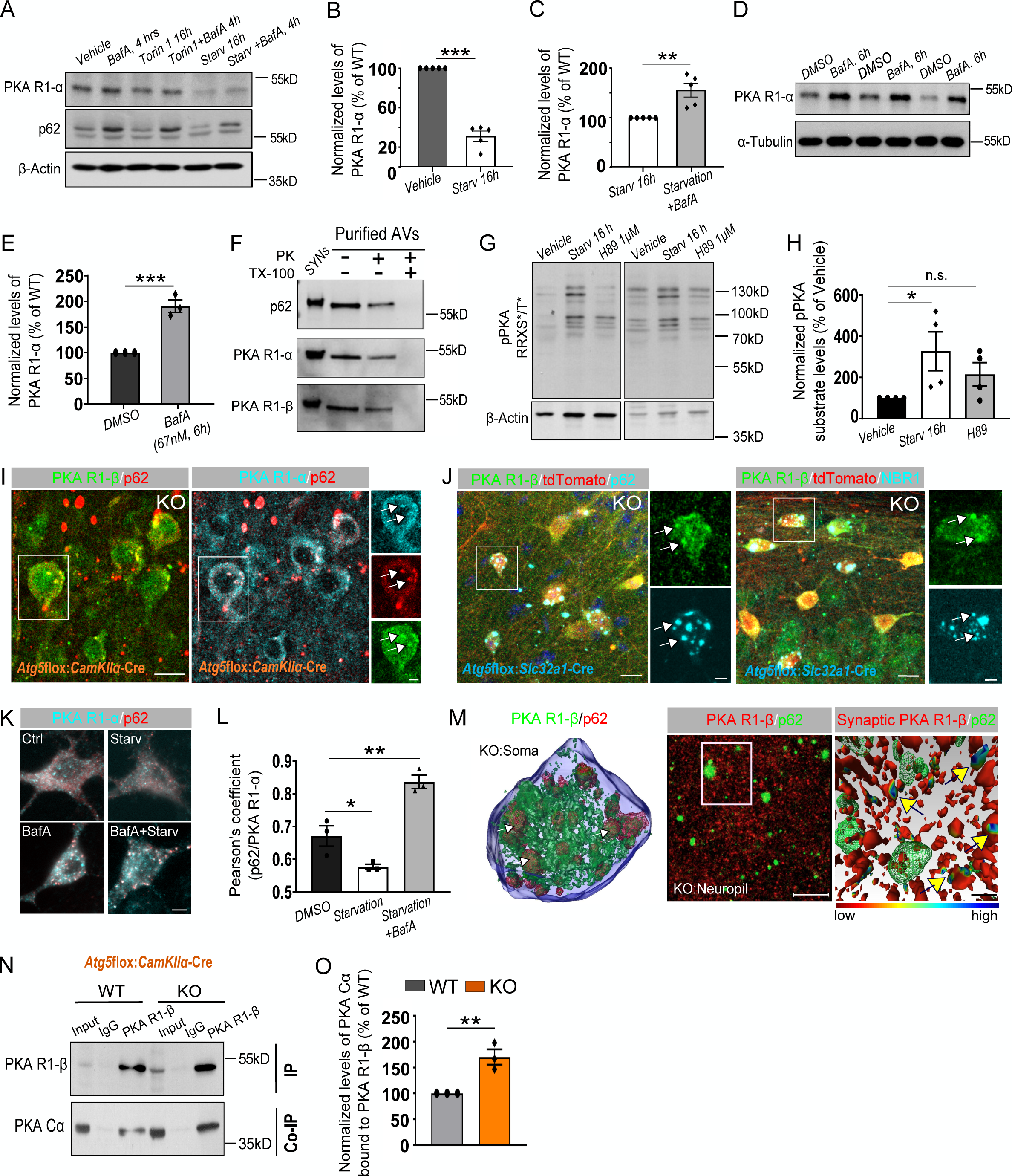
PKA R1-α and R1-β are degraded by starvation-induced autophagy in neurons. **(A)** Western Blot analysis of PKA R1-α protein levels in NSC34 cells, treated with 250 nM Torin1 or deprived of amino-acids and serum for 16 hours. Lysosomal degradation was blocked in the last 4 hours using 67 nM of Bafilomycin A (BafA). **(B)** Starvation significantly reduced PKA R1-α protein levels in NSC34 cells (vehicle set to 100%, starvation: 31.19± 5.085%, p=<0.0001, one-sample t-test). n=5. **(C)** Application of 67 nM of BafA for 4 hours before harvesting was sufficient to stabilize the PKA R1-α protein levels in starved NSC34 cells (starvation set to 100%, starvation+BafA: 48.52± 9.627%, p=0.002, one-sample t-test). n=5. **(D-E)** Treatment of NSC34 cells with BafA (67nM) for 6 h significantly increased PKA R1-α protein levels compared to the DMSO-treated group set to 100% (BafA: 191.0± 12.12%, p=0.0008, one-sample t-test). n=3. **(F)** Western blot analysis of purified autophagosomes (AVs) (50μg AVs/lane) with and without Proteinase K (PK) treatment. TX-100 (1% final) was used as a positive control for the activity of PK. Synaptosome lysates (Syn, 30μg/lane) were used as a positive control for the signal of antibodies. **(G, H**) Phosphorylation state of proteins containing PKA substrate RRXS/T motif was increased upon 16 hours of amino-acid and serum starvation in primary cortico-hippocampal neurons at DIV14 and was suppressed by 1 μM H89 supplementation in the media (Vehicle set to 100%, Starvation: 326.60±94.53%, H89: 214.80±56.87%, p=0.033, one-way ANOVA with Dunn’s multiple comparison test). n=4. The pPKA substrate levels were normalized to the total protein amount stained with Ponceau S. **(I-J)** Representative confocal images of the hippocampus CA1 area from *Atg5*flox:*CamKIIα*-Cre and *Atg5*flox:*Slc32a1*-Cre WT/KO mice, immunostained for PKA R1-α/β and co-immunostained for p62 and/or NBR1. White rectangular boxes indicate areas magnified to the right. White arrows indicate p62/NBR1 inclusion bodies positive for PKA R1. Scale bar: large panel 15 μm, small panel: 5 μm. **(K-L)** Representative fluorescent images and subsequent analysis of PKA R1-a/p62 colocalization (Pearson’s correlation coefficient) in primary cortico-hippocampal neurons, which have undergone 16 hours amino-acids and serum starvation and were additionally treated with BafA (67nM) to visualize the lysosomes (control: 0.671± 0.031, starvation: 0.577± 0.007, starvation+BafA: 0.836± 0.020, p_control vs starvation_ =0.040, p_control vs starvation+BafA_=0.0032, one-Way ANOVA with Dunnett’ multiple comparison test). n=3 (≥36 neurons in total). Scale bar: 5 μm. **(M)** 3D-reconstruction of *Atg5*flox:*CamKIIα*-Cre KO soma (most left), as well as the immunohistochemical profile and the 3D surface rendering of *Atg5*flox:*CamKIIα*-Cre KO neuropil revealing the PKA R1-β/p62 colocalization in the soma (white arrows), but not in processes (yellow arrows). Scale bar: 10 μM. See also fig. S4D,E for uncropped 3D reconstruction and voxel histogram. **(N)** Co-immunoprecipitation of endogenous PKA R1-β with PKA-Cα from *Atg5*flox:*CamKIIα*-Cre WT/KO mouse brain lysates (WT set to 100%, KO: 170.3± 14.88%, p=0.005, one-sample t-test, n=3). Input, 1.5% of the total lysate was added to the assay.

If R1α and R1β are a selective cargo of neuronal autophagosomes, then autophagy dysfunction should lead to their accumulation in p62 (SQSTM1)-positive inclusions. In agreement with this hypothesis, we observed a confinement of R1α and R1β to p62-containing puncta in cell soma of neurons of *Atg5*flox:*CamKIIα*-Cre KO mice (Fig. 3I). Loss of ATG5 in GABAergic neurons also caused the soma-confined accumulation of R1β to inclusion bodies containing p62, as well as another selective autophagy receptor NBR1 (Fig. 3J). In the soma of WT neurons the colocalization of R1α and p62 was regulated by starvation and could be facilitated by autophagy inhibition (Fig. 3K,L). Interestingly, in contrast to the cell soma (Fig. 3M, left), the enrichment of R1β at ATG5 KO deficient synapses was not accompanied by p62 accumulation (Fig. 3M, right, fig. S3D,E), suggesting that initial targeting of R1β to synaptic autophagosome membranes does not require the selective autophagy receptor p62 and might be mediated by its interaction with LC3. In agreement with this hypothesis, co-immunoprecipitation analysis revealed biochemical association of R1β with LC3 in the brain (fig. S3F). Finally, significantly more PKA R1α and R1β was found to be associated with PKA catalytic subunit in brain lysates of *Atg5*flox:*CamKII*α-Cre KO mice (Fig. 3N,O). Taken together, these results demonstrate that starvation-dependent autophagy regulates levels of inhibitory type 1 PKA subunits in neurons, and its loss stabilizes R1α and R1β levels, ultimately causing the sequestration of the PKA catalytic subunit.

### ATG5 regulates the cAMP/PKA/CREB1 signaling axis in the brain

A prediction from the data described above is that upregulated levels of type 1 PKA regulatory subunits in neurons with ATG5 deletion (without compensation of the PKA-Cα levels, see fig. S2) diminish PKA signaling. PKA activity is tightly controlled by cAMP and can be elevated upon supplementation of cells with the cAMP-elevating agent forskolin, a phenotype reflected in phosphorylation of CREB (pCREB) at Ser 133. As shown in Fig. 4A, CREB phosphorylation was increased in cultured WT cortico-hippocampal neurons treated with 10μM of forskolin for 5min, whereas total CREB levels were unaltered (Fig. 4A,B). In contrast, ATG5 deletion resulted in the absence of pCREB response to forskolin stimulation, indicating that cAMP-dependent PKA signaling is inhibited upon autophagy deficiency in neurons. Intriguingly, we observed a decreased forskolin-induced pCREB response also in mouse embryonic fibroblasts (MEF) isolated from mice where ATG5 deletion was driven by the addition of tamoxifen (*18*) (Fig. 4C,D), suggesting that the function of autophagy in the regulation of PKA signaling is not neuron-specific. To investigate whether neuronal ATG5 regulates PKA/pCREB activation *in-vivo*, we analyzed forskolin-dependent pCREB responses in cortical neurons isolated from 8-week-old *Atg5*flox:*CamKII*α-Cre:tdTomato WT or KO mice using quantitative high content screening microscopy. Our analysis revealed that forskolin-mediated activation of pCREB was abolished in cortical excitatory neurons in adult mice (Fig. 4E,F). These changes were independent of cAMP levels per se, which were unaltered both in brain lysates (Fig. S4A) and isolated neurons of *Atg5*flox:*CamKII*α-Cre KO mice (Fig. S4B,C). Activation of pCREB using a brief pulse of depolarizing KCl solution (40mM) was also impaired (Fig. 4E), suggesting that ATG5 could regulate neuronal activity-dependent pCREB activation.

**Figure 4.**
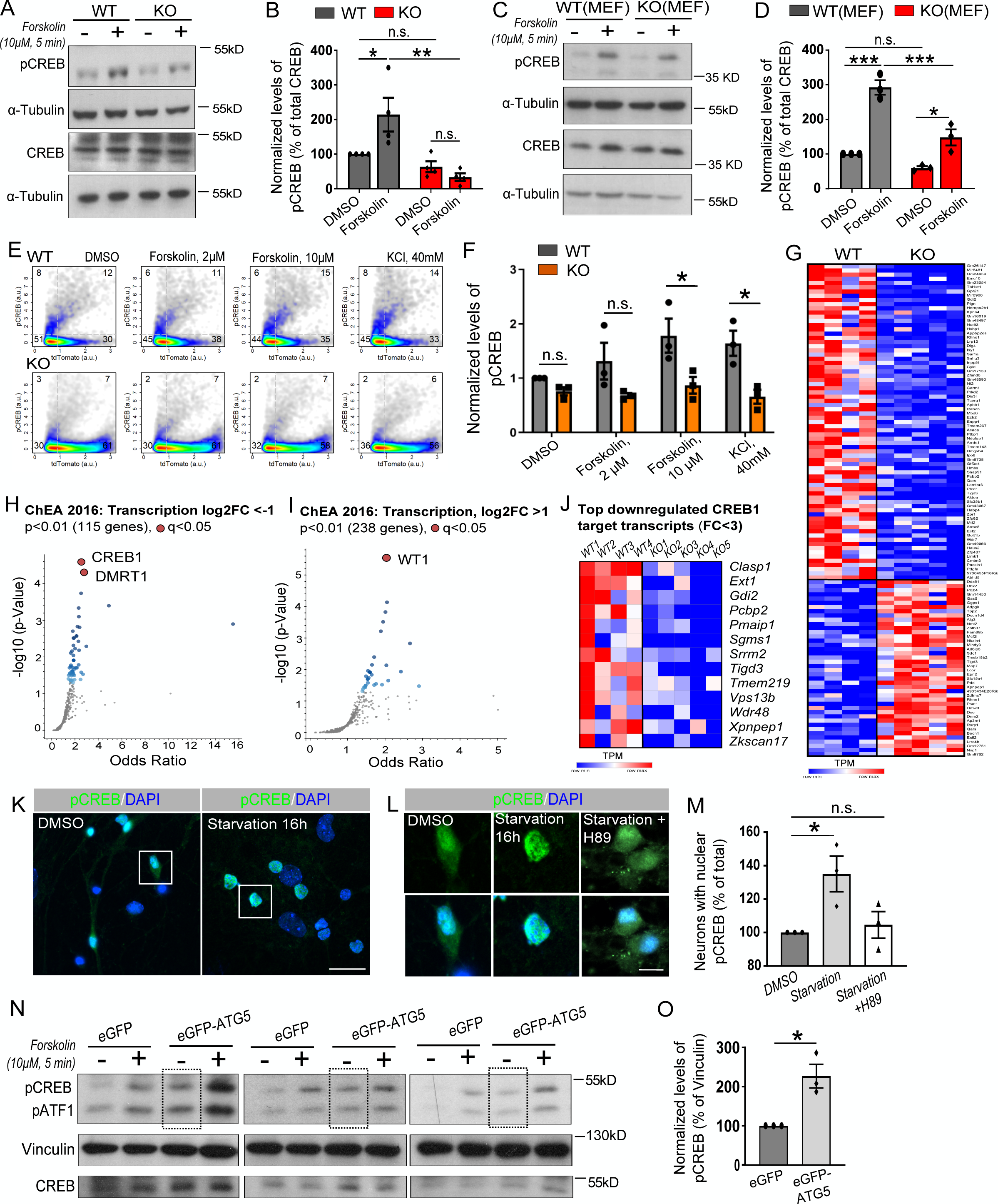
ATG5 regulates neuronal cAMP/PKA/pCREB signaling. **(A-B)** Western Blot analysis of pCREB protein levels in *Atg5*flox:*CAG*-Cre^Tmx^ WT/ KO cultured neurons at DIV14, treated with forskolin (10μM, 5 min) (WT_DMSO_ set to 100%, WT_Forskolin_: 214.17± 48.87%, KO_DMSO_: 63.26± 15.85%, KO_Forskolin_: 33.50± 10.84%, p_WT DMSO/WT Forskolin_=0.042, p_WT Forskolin/KO Forskolin_=0.002, two-way ANOVA with Tukey’s multiple comparisons). n=4. **(C-D)** Western Blot analysis of pCREB protein levels in forskolin-treated WT and ATG5 KO MEF cells (WT_DMSO_ set to 100%, WT_Forskolin_: 292.36± 20.76%, KO_DMSO_: 59.96± 4.85%, KO_Forskolin_: 148.10± 23.20%, p_WT DMSO/WT Forskolin_=0.0001, p_WT Forskolin/KO Forskolin_=0.0009, p_KO DMSO/KO Forskolin_=0.018, twoway ANOVA with Tukey’s multiple comparisons). n=3. **(E-F)** High-content screening microscopy analysis of pCREB fluorescence intensity in neurons isolated from 8-12-week-old *Atg5*flox:*CamKII*α-Cre:tdTomato WT/KO mice, treated either with DMSO or with 2μM, 10μM Forskolin or 40mM KCl (WT_DMSO_ set to 1, KO_DMSO_: 0.76±0.07, WT_Forskolin 2μM_: 1.31±0.34, KO_Forskolin 2μM_: 0.68±0.04, WT_Forskolin 10μM_: 1.78±0.31, KO_Forskolin 10μM_: 0.87±0.15, WT_KCl 40mM_: 1.64±0.23, KO_KCl 40mM_: 0.66±0.13, p_WT Forskolin 10μM/KO Forskolin 10μM_= 0.047, p_WT KCl 40mM/KO KCl 40mM_= 0.034, two-way ANOVA with Tukey’ s multiple comparisons test). n=3. **(G)** Next-generation RNA sequencing-based Kallisto transcriptome output of FACS-sorted neurons isolated from 3-week-old *Atg5*flox:*Slc32a1*-Cre:tdTomato WT/KO mice (cut-off p<0.02, FC<-2/>2) (n_WT_=4, n_KO_=5). See also fig. S4D for differential gene expression analysis. **(H,I)** EnrichR-based analysis of transcription factor-binding site enrichment using ChEA2016 databank. Analysis was applied to differentially downregulated (H) and upregulated (I) gene sets (cut off p<0.01, log2FC<-1/>1), obtained using Sleuth-based algorithm for gene expression analysis of RNA transcriptome shown in (G). **(J)** Top downregulated CREB1 target genes in autophagy-deficient FACS-sorted *Atg5*flox:*Slc32a1*-Cre:tdTomato KO neurons. **(K-M)** Representative fluorescent images and subsequent analysis of primary cortico-hippocampal neurons containing nuclear pCREB either under DMSO-treated condition or under condition when amino acids and serum starvation was induced for 16 hours in the absence or presence of 1μM of PKA inhibitor H89 (DMSO set 100%, starvation: 135.00± 10.64%, starvation + H89: 104.60±7.94%, p=0.041, one-way ANOVA with Dunnett’s multiple comparisons). n=3 (≥ 30 images). Scale bar: large panels: 15 μm, small panels: 5 μm. **(N,O)** The eGFP-ATG5 overexpression in MEF cells significantly increases the basal level of pCREB compared to eGFP-expressing MEF cells (eGFP: set to 100%, eGFP-ATG5: 226.70± 30.13%, p=0.017, one-sample t-test). n=3.

CREB belongs to the CREB1/CREM transcription factors subgroups, which bind to cAMP-responsive elements in target genes. To understand whether alterations of PKA signaling are reflected in the expression of CREM1/CREB target genes, we analyzed the gene expression in FACS-sorted WT and KO *Atg5*flox:*Slc32a1*-Cre:tdTomato neurons (see Fig. 1N) using the nextgeneration sequencing approach (Fig. 4G). We identified that ATG5 deletion caused downregulation of 732 transcripts (Fig. 4G, Suppl. Table 2). Although only a few genes were differentially expressed based on their corrected p-value (fig S4D, Suppl. Table 2), the analysis of transcription factor binding signatures (p< 0.01, FC<-2, 115 genes in total) revealed that 29.6 % of downregulated genes contained CREB1 response elements in their promoter region (Fig. 4H, Suppl. Table 2). On the other side, 66 out of 238 significantly upregulated genes (p<0.01, FC>2) were regulated by the WT1 transcription activity (Fig. 4I), which has been recently described as a crucial regulator of synaptic function and neuronal excitability (*33*). Intriguingly, among downregulated genes with CREB1 response elements, we identified genes encoding for proteins with a function in protein and vesicle trafficking (*Clasp1, Gdi2, Sgms1, Vps13b*) (Fig. 4J), suggesting that autophagy might indirectly regulate intracellular membrane trafficking by controlling the PKA/CREB1 signaling.

If neuronal levels of PKA type 1 inhibitory subunits are negatively regulated by starvation-induced autophagy (see Fig. 3), starvation should facilitate CREB translocation to the nucleus due to its increased phosphorylation by liberated PKA catalytic subunit. In agreement with this hypothesis, a proportion of neurons with nuclear pCREB was significantly increased upon amino acid and serum deprivation (Fig. 4K). This phenotype was PKA-dependent since blocking the catalytic activity of PKA by the ATP-site inhibitor H89 diminished pCREB translocation to the nucleus (Fig. 4L,M). Finally, to understand whether ATG5 is directly responsible for the regulation of the PKA/pCREB signaling axis, we analyzed the levels of pCREB in forskolin-stimulated MEF cells overexpressing either ATG5 tagged with eGFP or eGFP as a control. Overexpression of ATG5 augmented CREB phosphorylation responses upon forskolin treatment, indicating that ATG5 directly regulates PKA activity, likely by facilitating PKA type 1 inhibitory subunit degradation (Fig. 4N). Strikingly, pCREB levels were also increased in ATG5-overexpressing MEFs under forskolin-free conditions (Fig. 4O), indicating that ATG5, and thus autophagy could potentially fine-tune the sensitivity of PKA to cAMP levels in cells.

### Neuronal ATG5 regulates PKA-mediated phosphorylation of proteins enriched at the PSD of excitatory synapses

PKA-mediated regulation of CREB-dependent gene expression is known to be crucial for longterm memory (*34*). In addition, PKA-mediated phosphorylation can regulate neurotransmission by acting directly on presynaptic proteins regulating SV exo-/endocytosis and PSD-localized neurotransmitter receptors (*35*, *36*). The latter function of PKA in synaptic potentiation is an essential regulator of neuronal excitation. Thus, our findings above raise the question of whether autophagy regulates neuronal excitability by controlling PKA-dependent phosphorylation of the synaptic proteome. When PKA is activated, it phosphorylates substrates preferably on the recognition motif Arg-Arg-X-Ser/Thr-Y, and phosphorylated phospho-Ser/Thr residues can be monitored by using a specific antibody (RRXS/T). In WT neurons, application of forskolin induced a robust increase in phosphorylated phospho-Ser/Thr residues (Fig. 5A), whereas forskolin-dependent phosphorylation of PKA substrates was abolished in neurons lacking ATG5 *in-vitro* (Fig. 5B) and *in-vivo* (fig. S5A,B), suggesting a global reduction in PKA-dependent phosphoproteome upon autophagy deficiency.

**Figure 5.**
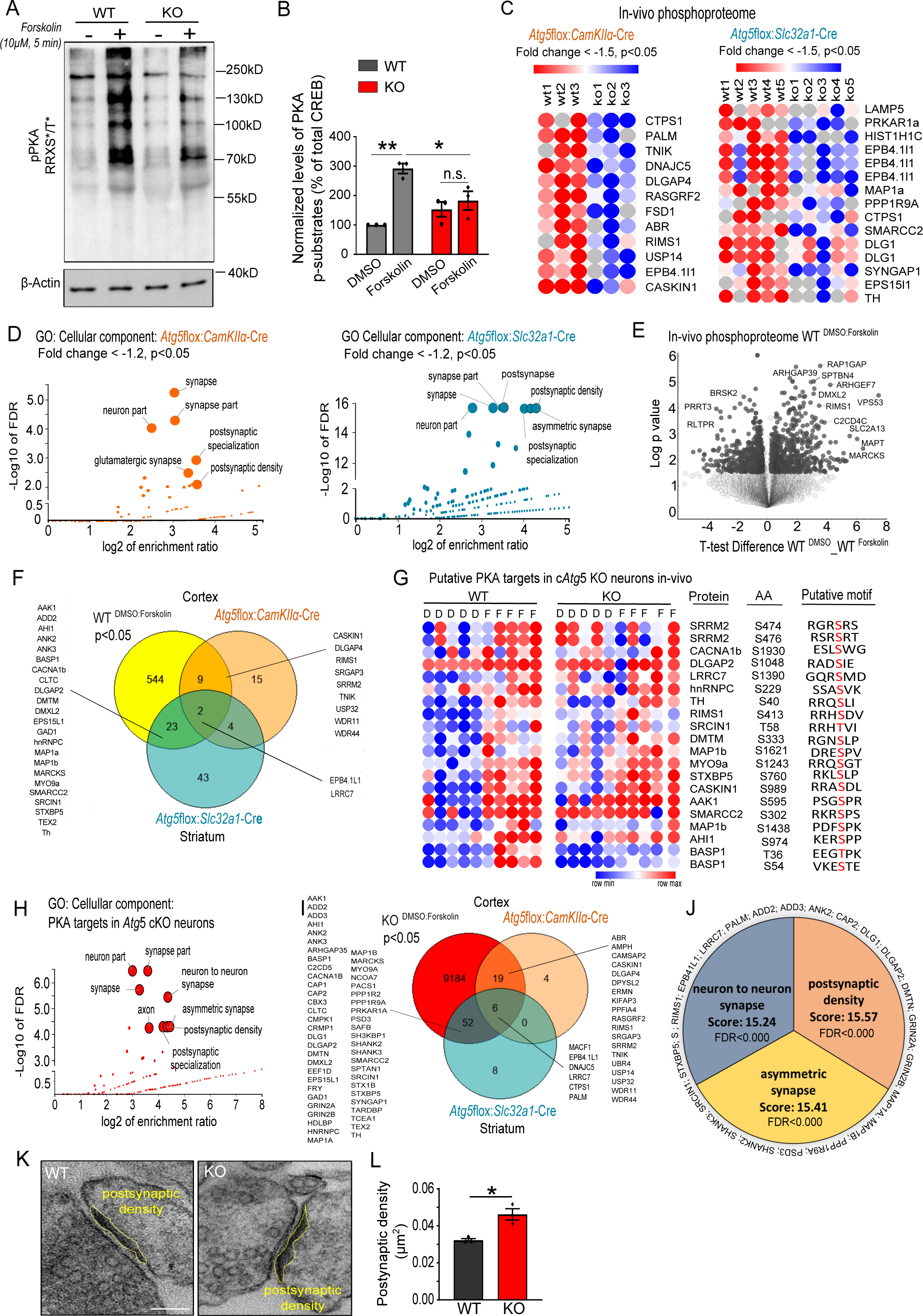
Neuronal ATG5 deficiency diminishes PKA-mediated phosphorylation of proteins enriched at the postsynaptic density of excitatory synapses. **(A-B)** Western Blot analysis of PKA substrates carrying RRXS/T motif in primary cortico-hippocampal *Atg5*flox:*CAG*-Cre^Tmx^ WT/KO neurons at DIV14, treated with 10 μM Forskolin for 5 min (WT_DMSO_ set to 100%, WT_Forskolin_: 291.85± 17.02%, KO_DMSO_: 152.97± 25.26%, KO_Forskolin_: 182.56± 31.89%, p_WTDMSO/WTForskolin_: 0.001, p_WTForskolin/KOForskolin_: 0.033, two-way ANOVA with Tukey’s multiple comparisons). n=3. **(C)** The top downregulated phosphopeptides (cut-off p<0.05, Fold change<-1.5) in either cortical lysates of *Atg5*flox:*CamKIIα*-Cre KO (n=3) or striatal lysates of *Atg5*flox:*Slc32a1*-Cre KO mice (n=5), identified by global phosphoproteome analysis (fig. S1O). The levels of phosphopeptides were normalized to their total protein levels. **(D)** WebGestalt-based GO analysis of “cellular component”-enriched terms among significantly downregulated phosphopeptides (cut-off p<0.05, Fold change <-1.2) in *Atg5*flox:*CamKIIα*-Cre KO and *Atg5*flox:*Slc32a1*-Cre KO brains. **(E)** Volcano plot of phosphopeptides identified using label-free phosphoproteome analysis of acute cortico-hippocampal slices from *Atg5*flox:*CamKIIα*-Cre WT mice treated either with DMSO or 50 μM Forskolin for 15 min (n=5). Phosphopeptides significantly upregulated (p<0.05) upon Forskolin treatment are highlighted in dark gray. **(F)** Venn diagram showing the number of phosphopeptides significantly downregulated in *Atg5*flox:*CamKII*α-Cre and *Atg5*flox:*Slc32a1*-Cre KO brains (cut-off p<0.05, Fold change <-1.2) and significantly upregulated in WT brain slices upon Forskolin treatment (cut-off p<0.05, Fold change >1.2). **(G)** Heat map representation of LFQ intensities of common phosphopeptides, identified in (F) and containing putative PKA phosphorylation motif (predicted by Perseus software) in *Atg5*flox:*CamKII*α-Cre WT and KO acute slices, treated either with DMSO or 50 μM Forskolin for 15 min (n=5). **(H)** WebGestalt-based GO analysis of “cellular component”-enriched terms among phosphopeptides represented in (G). **(I)** Venn diagram showing the number of phosphopeptides significantly downregulated in *Atg5*flox:*CamKII*α-Cre KO and *Atg5*flox:*Slc32a1*-Cre KO brains (cut-off p<0.05, Fold-change<-1.2) and the phosphophopeptides, which were “irresponsive” to Forskolin treatment in *Atg5*flox:*CamKIIα*-Cre KO acute slices (300μm) (p>0.05). **(J)** Top three enriched “cellular component” terms obtained by GO analysis of common phosphopeptides identified in (I). **(K-L)** EM analysis of the thickness of the postsynaptic density of *Atg5*flox:*CAG*-Cre^Tmx^ WT and KO cortico-hippocampal neurons (WT: 0.032± 0.001, KO: 0.046± 0.003). P=0.011, n=3 (≥21 synapses per condition). Scale bar: 120 nm.

To further define specific phosphorylation targets affected by neuronal ATG5 deletion, we decided to investigate phosphoproteome changes of *Atg5*flox:*CamKIIα*-Cre and *Atg5*flox:*Slc32a1-Cre* WT and KO brains using a spike-in SILAC mouse approach described above. This analysis yielded the identification of in total 104 significantly up- or downregulated (p<0.05, FC>1.2/<-1.2) phosphosites in *Atg5*flox:*CamKIIα*-Cre and 217 phosphosites in *Atg5*flox:*Slc32a1*-Cre lines whose abundance, when normalized to total protein levels, changed upon ATG5 KO condition. Of them, 31 and 82 phosphosites were decreased in *Atg5*flox:*CamKIIα*-Cre KO mice and *Atg5*flox:*Slc32a1*-Cre KO mice, respectively (FC<-1.2, p<0.05, Suppl. Table 3). Interestingly, in both mouse lines, gene products in which neuronal ATG5 deletion decreased phosphorylation included several scaffolding postsynaptic and a few presynaptic proteins (PALM, TNIK, DNAJC5, DLGAP4, RIMS1, EPB4.1L1, DLG1, SYNGAP1, and CASKIN1) (Fig. 5C) and were clustered in GO cellular component terms related to postsynaptic specialization of glutamatergic synapses in both mouse lines (Fig. 5D). On the general level, these data argue that autophagy deficiency decreases global phosphorylation of PSD proteins at excitatory glutamatergic synapses in both excitatory and inhibitory neurons.

Phosphorylation of PKA substrates in neurons is maintained at a relatively high state under basal conditions (*37*). Thus, global phosphorylation changes in brains lacking ATG5 could at least partially result from diminished PKA signaling. To directly test this hypothesis, we first performed an analysis of PKA phosphorylation substrates in forskolin-stimulated cortical slices of *Atg5*flox:*CamKII*α-Cre WT mice (Fig. 5E, Suppl. Table 4). Forskolin caused an increase in phosphorylation of several known PKA substrates, such as MAPT, RIMS1, STMN1, IP3R1, RASGRP2, SNAP-25, TH, most of which were enriched at the PSD of excitatory synapses (fig. S5C). By comparing PKA-dependent substrates in WT (FC<-1.2, p>0.05) and phosphorylation targets identified in total phosphoproteome of autophagy-deficient brains (Fig. 5C), we found that 11 substrates from *Atg5*flox:*CamKII*α-Cre KO brains and 25 substrates from *Atg5*flox:*Slc32a1*-Cre KO mice could be potential PKA targets (Fig. 5F). Subsequent analysis of these 36 substrates revealed that 20 of them contained a potential PKA phosphorylation motif and were less sensitive to forskolin treatment in *Atg5*flox:*CamKII*α-Cre KO cortical slices (Fig. 5G, fig. S5D,E). In agreement with the data above, cellular component GO enrichment analysis indicated that most of the putative PKA phospho-targets affected by the ATG5 deletion were localized to the PSD of glutamatergic synapses in both mouse lines (Fig. 5H). PKA substrates, identified by co-immunoprecipitation in cortical lysates of WT mice with PKA phosphorylation state-specific antibody (RRXS/T) were also enriched at the postsynaptic density as such (fig. S5F,G), whereas comparison of these targets with proteins identified in global phosphoproteome analysis of conditional ATG5 KO mice again yielded candidates confined to the PSD of excitatory synapses (fig. S5H,I). Finally, we found that the vast majority of substrates significantly downregulated in global KO phosphoproteome (see Fig. 5C) were not reacting to forskolin treatment in cortical slices prepared from *Atg5*flox:*CamKII*α-Cre KO mice (Fig. 5I) and those forskolin-“irresponsive” targets in KO neurons were enriched at the PSD of glutamatergic synapses (Fig. 5J)

The posttranslational modification of PSD-confined proteins by serine/threonine phosphorylations has been shown to have a profound effect on the dynamics of their localization. For instance, the anchoring of DLG at synapses is optimal when DLG is in the dephosphorylated state (*38*). If ATG5-dependent cAMP/PKA-dependent signaling is a crucial regulator of phosphorylation of PSD-confined proteins, then decreased phosphorylation of these proteins in the absence of ATG5 might affect the morphological property of the PSD. In agreement with this hypothesis, electron microscopic analysis reveals that the area of PSD was significantly increased in neurons lacking ATG5 (Fig. 5K,L). Collectively, these data indicate that autophagy functions in neurons to modulate the PKA-dependent phosphorylation of (post)synaptic scaffolding proteome independently of its role in synaptic protein homeostasis. Furthermore, our data suggest that ATG5-dependent protein phosphorylation can serve as a crucial regulator of postsynaptic protein localization and trafficking.

### Autophagy-dependent PKA signaling regulates AMPARs localization and function

What are the mechanistic implications of decreased phosphorylation of postsynaptic density-confined proteins for the physiology of ATG5 KO neurons? The postsynaptic density of synapses includes cytoskeletal scaffold proteins, receptors and ion channels, and the phosphorylation state of these components is known to be central to synaptic transmission (*39*). The size of postsynaptic density is a common characteristic of neuronal excitability *per se*, where the larger size of the PSD is positively correlated with higher numbers of AMPA-type glutamate receptors (AMPARs) (*40*, *41*). Furthermore, PKA directly regulates synaptic function via neuronal activity-dependent internalization of AMPAR subunit GLUR1 from the postsynaptic plasma membrane (*42*). This function of PKA requires the interaction of its type 2 regulatory subunit α with AKAP79/150 (*43*) and involves PKA-dependent phosphorylation of GLUR1 at Ser845 (*39*). Although our data indicate no alterations in levels of type 2 PKA regulatory subunits, it can still be plausible that autophagy-dependent PKA signaling regulates synaptic neurotransmission by directly controlling the trafficking of AMPARs. To test this hypothesis, we first analyzed the levels of GLUR1 phosphorylated at Ser845 in *Atg5*flox:*CamKIIα*-Cre and *Atg5*flox:*Slc32a1*-Cre WT and KO mice by Western Blotting. However, in agreement with data from global phosphoproteome profiling (Suppl. Table 3) and in line with unaltered levels of AKAP79/150 in total proteome data (Suppl. Table 1, fig. S6A), pGLUR1 levels normalized to the total GLUR1 levels were not changed in excitatory neurons lacking ATG5 (fig. S6B-E). Interestingly, GLUR1 phosphorylation at Ser845 was slightly reduced in striatal lysates from *Atg5*flox:*Slc32a1*-Cre KO mice, although this effect did not completely reach significance (p=0.050, fig. S6F-H). This phenotype was accompanied by a significant reduction in total levels of AKAP79/150 in inhibitory ATG5-deficient neurons (fig. S6I), suggesting that under basal conditions PKA-dependent phosphorylation might be differentially regulated in excitatory and inhibitory neurons.

Since the absence of alterations of direct PKA-dependent GLUR1 phosphorylation in excitatory neurons does not explain excitability changes in *Atg5*flox:*CamKIIα*-Cre KO mice (see Fig. 1), we took a closer look at the molecular function of proteins with decreased phosphorylation in mice lacking neuronal ATG5. In agreement with the fact that PKA phosphorylates multiple cytoskeletal substrates in non-neuronal cells (*44*), “cytoskeletal protein binding” (32% of proteins) and “actin-binding” (20% of proteins) were the most abundant GO molecular function terms among downregulated phospho-targets in autophagy-deficient neurons (Fig. 6A, fig. S6J,K, Suppl. Table 4, see also Fig. 5I). Strikingly, analysis performed using the human phenotype ontology database indicated that proteins identified as being decreased in their phosphorylation state in brains of autophagy-deficient mice are associated with EEG abnormality and seizure onset in humans (Fig. 6A). These data, taken together with the fact that about 37% of “cytoskeletal protein binding” proteins identified in Fig. 5I are directly involved in endocytosis (CLTC) and trafficking (ANK3, DLG1, DPYSL2, EPB41L1, PPP1R9A, SHANK3 and STXBP5) (*45*–*52*) of GLUR1 subunit of AMPARs (Fig. 6B), prompted us to speculate that altered GLUR1 trafficking, rather than its direct phosphorylation by PKA is a crucial determinant of neuronal excitability in ATG5 KO neurons. In line with this hypothesis and the data described above (see Fig. 5K,L), the colocalization of GLUR1 and postsynaptic density marker PSD95 was increased in neurons lacking ATG5 *in-vitro* (Fig. 6C,D) and *in-vivo* (fig. S6L). This phenotype directly resulted from the ATG5 loss-of-function since the overexpression of ATG5 tagged with eGFP normalized PSD-confined GLUR1 localization (Fig. 6E). On the other hand, the overexpression of type 1 PKA inhibitory subunits α and β in control neurons mimicked the phenotype of ATG5 KO neurons and caused the accumulation of GLUR1 at the PSD (Fig. 6F), indicating that increased postsynaptic levels of GLUR1 in autophagy-deficient neurons are a result of diminished PKA signaling. Of note, although total protein levels of GLUR1 were altered neither in global SILAC-based proteomic analysis (Suppl. Table 1), nor in the WB analysis of hippocampal lysates (Fig. S6B) (both reporting on soma, as well as neuropil levels of GLUR1), we detected a slight increase in levels of GLUR1 (log2 fold change=0.62) and GLUR4 (log2 fold change=0.84) subunits in FACS-sorted somata of cortical excitatory neurons. These changes could reflect compartment-specific role of autophagy in the degradation of soma-confined GLUR1 receptors. To directly probe whether the defects in GLUR1 distribution reflect diminished retrieval of GLUR1 from the plasma membrane, we employed pHluorin, fusion between the N-terminus of GLUR1 and a pH-sensitive variant of GFP that undergoes quenching within the acidic vesicle lumen and is dequenched upon exocytic fusion with the plasma membrane. In ATG5 KO neurons, superfused with buffers titrated either to pH 5.5 (to reveal the surface exposure of pHluorin) or to pH 8.5 (to reveal the total levels of pHluorin) (fig. S6M), we observed a significant elevation of the surface-stranded/total GLUR1-pHluorin ratio compared to WT neurons (Fig. 6G-I).

**Figure 6.**
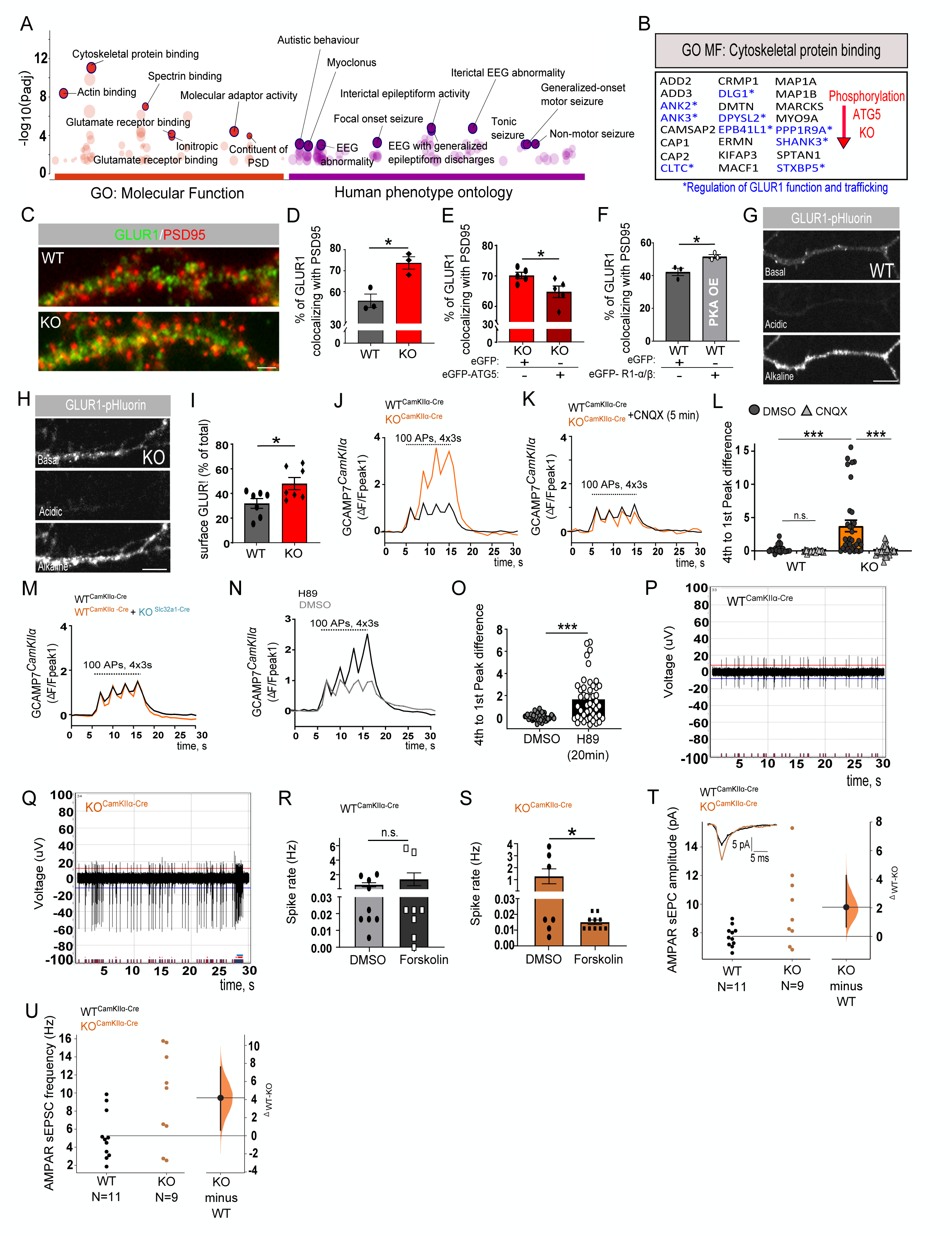
Autophagy-dependent PKA signaling controls postsynaptic GLUR1 localization and is responsible for the regulation of neuronal excitability. **(A)** g:Profiler-based analysis of GO molecular function and human phenotype enriched terms of “irresponsive” PKA target proteins in autophagy-deficient neurons identified in Fig. 5I (cut-off p<0.05, FC<-1.2). See also fig. S6J. **(B)** Representation of proteins from GO molecular function term “cytoskeletal protein binding” shown in (A). Highlighted in blue are proteins with a published function in GLUR1 function and trafficking. **(C-D)** Analysis of GLUR1 and PSD95 colocalization (% of GLUR1 puncta colocalizing with PSD95) in cultured *Atg5*flox:*CAG*-Cre^Tmx^ WT and KO neurons at DIV14 (WT: 55.67±3.18%, KO: 73.67±2.96%, p=0.014, n_WT_=23, n_KO_=26 from n=3, unpaired two-tailed t-test). **(E)** The overexpression of eGFP-ATG5 in *Atg5*flox:*CAG*-Cre^Tmx^ KO neurons diminishes the of GLUR1/PSD95 colocalization compared to the eGFP-overexpressing KO neurons (KO_eGFP_: 70.17±1.03%, KO_eGFP-ATG5_: 64.80±1.90%, n_KO+GFP_=33, n_KO+GFP-ATG5_=37 from n=5, p=0.038, unpaired two-tailed t-test). **(F)** The coexpression of eGFP-PKA R1-α and eGFP-PKA R1-β in cultured *Atg5*flox:*CAG*-Cre^Tmx^ WT neurons results in significantly increased colocalization between GLUR1 and PSD95 compared to the eGFP-expressing WT neurons (WT: 42.26±2.26%, KO: 51.67±1.20%, p=0.021, n_WT+GFP_=46, n_WTGFP+PKA R1 OE_=46 from n=3, unpaired two-tailed t-test). Analysis in (D-F) is performed using a custom-written Plugin for ImageJ (Data S2). **(G-I)** Live-cell imaging analysis of GLUR1 surface fraction in *Atg5*flox:*CAG*-Cre^Tmx^ WT/KO cultured neurons transfected with GLUR1-pHlourin. The total surface levels of GLUR1-pHlourin were determined from the difference in fluorescence intensity between acidic and alkaline buffer conditions (see also fig. S6M) (WT: 31.94±3.99%, KO: 48.01±4.93%, p=0.026, unpaired twotailed t-test). n_WT_=35, n_KO_=48 neurons from n=5. Scale bar in (H): 5 μm. **(J-L)** GCAMP7f responses to tetanic stimulation (four tetani, 100 APs at 100Hz, 3s interval) in *CamKII*α-Cre-eGFP-transduced primary cortico-hippocampal *Atg5*wt:wt/tdTomato (WT) and *Atg5*flox:flox/tdTomato (KO) neurons expressing m*CamKII*α-jGCaMP7f and treated either with DMSO (J) or with 10 μM CNQX for 5 min (K). Autophagy deficient cells showed significantly increased facilitation of GCAMP7f signal to electrical stimulation (ΔF/Fpeak1) compared to the WT, a phenotype which was blocked by the application of CNQX (WT_DMSO_: 0.28± 0.12, WT_CNQX_: −0.07± 0.03, KO_DMSO_: 3.74± 0.86, KO_CNQX_: −0.014± 0.13, p _WT DMSO/ KO DMSO_<0.0001, p_KO DMSO/KO CNQX_<0.0001, two-way ANOVA with Tukey’s multiple comparisons test). n_WT DMSO_= 25, n_WT CNQX_= 27, n_KO DMSO_= 33, n_KO CNQX_= 30 from n=4. See also fig. S6N for changes in WT neurons upon CNQX treatment. **(M)** GCAMP7f responses to tetanic stimulation (four tetani, 100 APs at 100Hz, 3s interval) in h*GAD67*-chl-iCre-transduced primary cortico-hippocampal *Atg5*wt:wt/tdTomato (WT) and *Atg5*flox:flox/tdTomato (KO) neurons expressing m*CamKII*α-jGCaMP7f (n_WT_= 69; n_KO_= 64 from n=4). Quantification in fig. S6O. **(N-O)** GCAMP7f responses to tetanic stimulation (four tetani, 100 APs at 100Hz, 3s interval) in *CamKII*α-Cre-eGFP-transduced primary cortico-hippocampal *Atg5*wt:wt/tdTomato neurons expressing *mCamKIIα*-jGCaMP7f and treated either with DMSO or with 10 μM of H89 for 20 min (DMSO: 0.07± 0.04, H89: 1.668± 0.28, p <0.0001, unpaired two-tailed t-test). n_DMSO_=48, nH89=47 neurons from n=3. **(P, Q)** Example traces of the network activity of primary cortico-hippocampal WT and *Atg5*flox:flox/*CamKII*-Cre KO neurons analyzed using the MEA system. (**R, S**) Increased spike rate in ATG5 KO excitatory neurons is rescued by long-term application of Forskolin (10μM for 72 hours) (WT_DMSO_: 0.59± 0.29, WT_Forskolin_: 1.35± 0.88, KO_DMSO_: 1.27± 0.62, KO_Forskolin_: 0.02± 0.001, p_KO DMSO/KO Forskolin_=0.027, unpaired two-tailed t-test). n_WT DMSO_=9, n_WT Forskolin_=8, n_KO DMSO_: 7, n_KO Forskolin_=10. **(T,U)** AMPAR-mediated sEPSCs in cortical pyramidal neurons of 12-week-old mice *Atg5*flox:flox mice, which were stereotactically injected either with *CamKII*α-eGFP AAV (WT) or with *CamKIIα*-Cre-eGFP AAV (KO) into the perirhinal cortex layer II/III. (T) Amplitude of AMPAR sEPSC is increased in pyramidal KO neurons compared to the WT (WT: 7.75± 0.72 pA, KO: 9.79± 2.75 pA; n_WT_=11, n_KO_=9; unpaired mean difference between WT and KO (Δ_WT-KO_): 2.04 [95%CI: 0.629, 4.22]; p_WT/KO_=0.0152 (two-sided permutation t-test)). Inset: representative traces of a WT sEPSC (petrol) and a KO sEPSC (ochre). (U) Frequency of AMPAR sEPSC is increased in pyramidal KO neurons compared to the WT (WT: 5.27± 2.66 Hz, KO: 9.47± 5.16 Hz; n_WT_=11, n_KO_=9; Δ_WT-KO_: 4.19 [95%CI: 0.57, 7.56]; p_WT/KO_=0.0326 (two-sided permutation t-test)). Each data point represents the mean of a 1 min recording interval of an individual neuron. The filled curve indicates the resampled Δ distribution (5000 bootstrap samples), given the observed data. Horizontally aligned with the mean of the test group, the Δ is indicated by the black circle. The 95% confidence interval of the mean difference is illustrated by the black vertical line.

Next, we wondered if increased density of GLUR1-containing AMPAR is the cause of neuronal hyperexcitability observed in *Atg5*flox:*CamKIIα*-Cre KO mice (see Fig. 1). We first employed an *in-vitro* GCAMP7f imaging assay to measure neuronal excitability. During action potential firing, voltage-dependent Ca^2+^ channels are activated, leading to an activity-dependent rise in intracellular Ca^2+^ levels, which can be detected by using the genetically encoded Ca^2+^-sensing probe GCAMP7f. To monitor calcium signals by live-cell imaging, primary cortical neurons isolated from *Atg5*wt:wt/tdTomato or *Atg5*flox:flox/tdTomato mice were infected with an adeno-associated (AAV) virus encoding a *CamKIIα*-GCaMP7f construct and, additionally, with *CamKIIα*-Cre AAV to induce a KO. As the GCaMP7f reporter is controlled by a *CamKIIα* promoter, only neuronal calcium signals from excitatory neurons were captured. We compared the calcium responses evoked by tetanic stimulation (four tetani, 100 APs at 100Hz, 3s interval) per cell and found that KO excitatory neurons had significantly increased facilitation of evoked calcium signals compared to controls (Fig. 6J). This facilitation was completely abolished by preincubation of KO neurons with the broad-spectrum AMPA/kainate receptor antagonist CNQX, confirming that neuronal excitability in autophagy-deficient neurons is driven by glutamate-bound AMPARs (Fig. 6JK,L). Of note, the application of CNQX for 5 minutes also reduced the facilitation in WT, albeit with less efficiency, indicated by a significant difference between two groups, but not in multiple comparison analysis (Fig. 6L, fig. S6N). This function of autophagy in the regulation of neuronal excitability was cell-autonomous for excitatory neurons (Fig. 6M, fig. S6O) and required the PKA activity since the application of PKA inhibitor H89 reproduced the facilitation of evoked calcium responses in control neurons (Fig. 6 N,O). Furthermore, long-term application of the cAMP booster forskolin (72h) was sufficient to diminish the frequency of action potential firing in excitatory ATG5 KO primary neurons, whereas the frequency of action potentials in WT neurons was not altered (Fig. 6P-S).

Finally, to assess if neuronal hyperexcitability, caused by defects in autophagy-dependent PKA signaling (see above), is reflected in the properties of AMPAR currents of cortical pyramidal neurons, we performed whole-cell patch-clamp recordings. We recorded spontaneous AMPAR-mediated excitatory postsynaptic currents (sEPSCs) in acute brain slices of 12-13-week-old *Atg5*flox:flox mice, which received a stereotactic injection of either control AAV (*CamKIIα*-eGFP) or *CamKIIα*-Cre-eGFP AAV into the layer II/III of the perirhinal cortex. We found that acute ATG5 deletion in these cortical excitatory neurons increased both the amplitude and the frequency of AMPA-mediated sEPSCs (Fig. 6 T,U). Taken together with the data described above, our findings strongly suggest that *Atg5* deficiency in postnatal excitatory neurons causes AMPAR-mediated hyperexcitability by increasing the number of synaptic AMPARs.

Collectively, our data reveal a previously unknown role for autophagy in the control of neuronal excitability via regulation of the phosphorylation status of postsynaptic density-confined proteins. Mechanistically, this function of autophagy involves starvation-induced activation of type 1 PKA signaling at synapses, which via cytoskeletal proteins phosphorylation controls the synaptic accumulation of AMPARs and regulates neuronal excitability.

## Discussion

In this study, we provide evidence that neuronal autophagy maintains the excitability of glutamatergic synapses by regulating the intracellular availability of inhibitory type 1 PKA subunits. To our surprise, we find that the crucial autophagy modifier ATG5 is not only dispensable for the survival of cortical excitatory neurons (*18*) but is also not required to maintain the viability of forebrain inhibitory neurons. Most importantly, the data presented here demonstrate that autophagy contributes to neuronal physiology by controlling the starvation-dependent PKA-mediated phosphorylation of postsynaptic density cytoskeletal proteins, a phenotype that regulates functional properties of AMPAR-mediated synaptic currents.

Autophagy is currently linked to neuronal function by mechanisms, which include its role in the clearance of damaged and/or defective intracellular components (*53*). This role of autophagy in neuronal homeostasis is especially crucial at synapses, which possess high metabolic demands and are vulnerable to accumulated damage and stress (reviewed in (*15*)). Autophagy has been described to regulate the degradation of several pre- and postsynaptic proteins (*12*, *54*–*56*), axonal ER (*13*), and synaptic mitochondria (*14*, *57*) and, is, thereby, crucial for the regulation of synaptic plasticity, including long-term depression (LTD) (*11*, *25*, *58*, *59*). Intriguingly, here we find that ATG5-mediated autophagy does not primarily contribute to the regulation of bulk levels of SV and/or mitochondrial proteins in excitatory or inhibitory neurons i*n-vivo*. Instead, our data suggest that autophagy indirectly regulates the synaptic function by controlling the PKA-dependent phosphorylation state of postsynaptic proteins, independently of their degradation. This function of autophagy requires the targeting of inhibitory PKA R1α and R1β subunits to neuronal autophagosomes and is induced by nutrient deprivation, including amino acid and serum starvation. Autophagy-deficient neurons reveal accumulation of synapse-confined PKA R1α and R1β, a phenotype that sequesters PKA catalytic subunit and impairs neuronal PKA signaling. Several lines of evidence indicate that starvation-induced autophagy is a crucial regulator of PKA signaling in neurons. First, we demonstrate that amino acid and serum deprivation induces phosphorylation of PKA substrates in neurons (Fig. 3). Second, the cAMP-mediated pCREB responses are diminished in autophagy-deficient excitatory and inhibitory neurons in *in-vivo* and *in-vitro*, whereas ATG5 overexpression is sufficient to activate the PKA signaling (Fig. 4). Finally, by analyzing the forskolin-induced phosphoproteome of acute brain slices, we demonstrate that ATG5 deletion decreases the phosphorylation state of several PKA putative neuronal target proteins (Fig. 5). PKA signaling in neurons is known to be a key player in the synaptic plasticity-induced protein synthesis, modulation of neurotransmitter receptor channel conductance, and activity-dependent AMPAR trafficking (*34*, *39*). In agreement with the essential role of PKA signaling in neurotransmission, we revealed that autophagy-dependent PKA signaling has an integral function in regulating the surface levels of the AMPAR subunit GLUR1 and is crucial in regulating the properties of synaptic AMPARs. Although the augmented frequency of sEPSCs described in the current study can also be a result of an increased number of synapses in autophagy-deficient neurons, our global proteome data analysis does not reveal changes in bulk levels of SV proteins. This data, taken together with our earlier findings, indicating no alteration in the number of spines in autophagy-deficient neurons (*18*), suggests the upregulation in synaptic GLUR1 levels as a primary mechanism behind increased AMPA-mediated spontaneous neurotransmission. In contrast to the majority of AMPARs, which contain GLUR2, GLUR1-containing AMPARs are Ca^2+^ permeable, and their permeability can be regulated by PKA (*60*). Whether autophagy regulates Ca^2+^ permeability of GLUR1-containing AMPAR is currently unclear and should be addressed in future studies, although our data indicating increased GCAMP7 signals in excitatory neurons lacking ATG5 (Fig. 6) would support this hypothesis. Equally unclear is the role of autophagy-dependent PKA signaling in regulating the trafficking of GLUR2 and other subunits of AMPARs. Recent work suggests that autophagy is required for GLUR2 internalization during long-term depression (*11*) and several other studies report that autophagy regulates GLUR1 and GLUR3 protein levels (*12*, *61*), suggesting a generalized effect of autophagy on AMPARs function, which can either occur via canonical mechanisms involving AMPAR degradation or be mediated via novel PKA-dependent phosphorylation pathway, presented here.

One of the most intriguing finding in our study is that the autophagy-mediated PKA signaling primarily regulates the phosphorylation state of cytoskeletal scaffolding proteins confined to the postsynaptic density of excitatory synapses, whereas its contribution to the regulation of presynaptic and/or mitochondrial proteins is minor. Intriguingly, the majority of candidates identified in our global phosphoproteomics analysis (including LRRC7, DLGAP4, DLGAP2, SHANK3, STXBP5, SYNGAP1, EPB4.1L1) have been recently described to be essential for PKA-dependent synaptic downscaling of glutamatergic neurons (*62*). Homeostatic synaptic scaling defines a negative regulation of postsynaptic strength due to increased or decreased neuronal activity and is crucial to prevent runaway excitation and maintain circuit stability in the neocortex. Homeostatic synaptic scaling is achieved by upregulation (upscaling) or downregulation (downscaling) of functionally available AMPARs at postsynaptic density by mechanisms, which include post-translational modifications of AMPARs itself, as well as postsynaptic scaffolding and accessory proteins that interact with AMPARs (*63*). Our data suggest that neuronal autophagy could function as a potential modifier of homeostatic synaptic plasticity, although future studies will be needed to support this hypothesis. Interestingly, homeostatic synaptic plasticity is disrupted in autism spectrum disorders (ASD) (*64*), a clinically heterogeneous group of neurodevelopmental diseases characterized, among others, by dysfunctional autophagy (*65*). Several ASD-linked proteins have been found to be deregulated in their phosphorylation state in autophagy-deficient excitatory and inhibitory neurons in the current study, including SHANK3, SYNGAP1, EPB4.1L1, DMXL2, DPYSL2, and RIMS1 (Fig. 5, Fig. 6a). Thus, it is possible, if not likely, that autophagy contributes to ASD etiology by regulating the PKA-dependent homeostatic synaptic plasticity, among other mechanisms. Indeed, the loss of autophagy has been associated with autistic-like synaptic pruning defects (*20*), whereas children with ASD are more likely to have epilepsy (*66*), akin to the appearance of seizures in mice conditionally lacking ATG5 reported here.

The work presented here identifies amino acids and serum starvation as an inducer of autophagy-mediated degradation of PKA type 1 inhibitory subunits in excitatory and inhibitory mouse forebrain neurons. Interestingly, this induction is independent of mTORC1 activity (Fig. 3) and is specific to PKA R1α and R1β, whereas the protein levels of PKA type 2 subunits are not affected by autophagy deficiency. These findings support the earlier study performed in non-neuronal cells, where R1α was reported to be localized to autophagosomes (*67*) and are in line with the recent data suggesting dynamic crosstalk between autophagy machinery and PKA (*68*, *69*). PKA R1α has also been observed in multivesicular bodies in mouse embryonic fibroblasts, suggesting the existence of alternative pathways for PKA type 1 trafficking and degradation in cells (*70*). What determines the specificity of degradation of PKA R1α and R1β subunits by autophagy? The PKA type 2 subunits are typically recruited to membranous compartments by AKAPs, whereas the PKA regulatory subunits type 1 are mainly cytosolic, although their association with mitochondria has also been reported (*31*). We believe that cytosolic confinement of PKA type 1 subunits predisposes them to starvation-induced autophagy by bulk sequestration of the cytoplasm. Degradation of PKA R1α by AKAP11-mediated selective autophagy was recently described to be required to maintain mitochondrial metabolism in cancer cells (*32*). Our study did not identify changes in AKAP11 in proteomic analysis of mice conditionally lacking ATG5, suggesting that autophagy-dependent degradation of PKA type 1 regulatory subunits in neurons could occur independently of AKAP11 and possibly be mediated by an interaction of PKA R1β with LC3 (see Fig. S3).

One final unresolved question is how mTORC1-independent induction of PKA degradation upon starvation is achieved. mTORC1-independent autophagy can occur via the ULK1-dependent pathway, mediated by the protein phosphatase 2A (*71*). Alternatively, growth factors and cytokines can also inhibit the VPS34 complex, possibly via the activation of the IP3/Ca^2+^/Calpain pathway (*72*, *73*). Further experiments are required to understand the relationship between the mTORC1-independent induction of autophagy-mediated PKA signaling in neurons. However, given the fact that the protein phosphatase 2A- and the Ca^2+^/Calpain-dependent pathways are both known mediators of synaptic function (*74*, *75*), we believe that the mechanism of mTORC1-independent autophagy might have been evolved in neurons to be specifically required at synapses, which are located far away from the soma, containing the components of mTORC1-sensing machinery.

Increased surface localization of AMPA receptors at the postsynaptic density of neurons is a known trigger of abnormal synchronous activity, a phenotype that can ultimately result in epileptic seizures (*76*). Together with our observation of spontaneous recurrent seizures in mice with ATG5-deficiency in forebrain excitatory (but not inhibitory) neurons (Fig. 1) and earlier study reporting epilepsy in *Atg7*flox:*CamKII*α-Cre KO mice (*77*), we propose a model according to which autophagy controls neuronal excitability by regulating the PKA-dependent efficiency of AMPA-mediated glutamatergic neurotransmission. Interestingly, nutritional interventions, such as a lipid-rich ketogenic diet, are a known modulator of neuronal excitability in patients suffering from epilepsy (*78*). Although relatively efficient, the ketogenic diet has several side effects, including leg cramps and digestive issues. The data presented here provide a therapeutic potential for the use of FDA-approved autophagy-inducing drugs (i.e. metformin and spermidine) for the treatment of epilepsy. Strikingly, PKA signaling has been recently shown to restore synaptic responses in a mouse model of amyotrophic lateral sclerosis (*79*), a disease that is characterized by defective autophagy, among others (*80*). We believe that the components of PKA signaling machinery should be explored as a potential target for therapeutic restoration of synaptic function in other neurodegenerative conditions characterized by autophagy dysfunction, including Parkinson’s disease and Alzheimer’s diseases.

## Methods

### Mouse models

All mice were housed in ventilated polycarbonate cages at standard 12/12 hour day/night cycles. Food and water were provided *ad libitum*. All animal experiments are approved by the LANUV Cologne NRW and ethics committee and are conducted to the committee’s guidelines (AZ 81-02.04.2020.A418, AZ 84-02.04.2016.A041, AZ 81-02.05.40.20.075, AZ 84-02.04.2016.A451). Conditional tamoxifen-inducible Atg5 and Atg16l1 KO mice (Atg5flox/flox: B6.Cg-Tg(CAG-Cre/Esr1*)5Amc/J) and *Atg5*flox/flox:*CamKIIα*-Cre mice previously described (*17*). Mice conditionally lacking ATG5 and/or ATG16L1 in inhibitory neurons were created by crossing *Atg5*flox/flox mice with *Vgat-ires*-Cre line (#016962, The Jackson Laboratory), kindly provided by Prof. Matthew Poy (Johns Hopkins University School of Medicine, St. Petersburg, USA). Conditional ATG5 KO reporter mice were created by crossing both lines to Ai9(RCL-tdT) (tdTomato) line (The Jackson Laboratory) that was received from Prof. Matteo Bergami (CECAD, Cologne, Germany). *Atg5*flox/flox:*Slc32a1*-Cre and *Atg5*flox/flox:*CamKIIα*-Cre mice were used as knockouts (KO). In order to reduce the number of animals used for experiments (3R principle), mice carrying *Atg5*wt/wt:*Slc32a1*-Cre^tg^, *Atg5*wt:wt/*Slc32a1*-Cre^wt^, *Atg5*flox/wt:*Slc32a1*-Cre^wt^, *Atg5*flox/wt: *CamKIIα*-Cre^wt^ or *Atg5*flox/flox: *CamKIIα*-Cre^wt^ genotypes were used as controls (wildtype (WT)) (no phenotypical abnormalities were detected between all mentioned “WT” genotypes). Similarly, *Atg16L1*flox/flox:*Slc32a1*-Cre^tg^ mice were used as knockouts (KO), whereas mice carrying *Atg16L1*wt/wt:*Slc32a1*-Cre^tg^ and *Atg16L1*wt:wt/*Slc32a1*-Cre^wt^, *Atg5*flox/wt:*Slc32a1*-Cre^wt^ genotypes were used as WT. Primers used for mouse genotyping are shown in Table S5.

### Purification of mouse autophagosomes (AVs)

The isolation of purified AVs was performed from ten cortices and hippocampi of adult (postnatal day 60, P60) C57BL/6J male mice as described previously (*7*). Briefly, the tissue was homogenized in 10% sucrose, 10 mM Hepes and 1 mM EDTA (pH 7.3) by 20 strokes using a Dounce glass homogenizer. The material was then diluted in half volume of homogenization buffer (HB) (250 mM sucrose, 10 mM Hepes and 1 mM EDTA, pH 7.3) containing 1.5 mM glycyl-L-phenylalanine 2-napthylamide (GPN). After incubation at 37°C for 7 min, the material was centrifuged at 2000 xg for 2 min at 4°C, and the nuclear pellet was discarded. The isolated post-nuclear supernatant was then loaded on discontinuous Nycodenz gradients that were centrifuged at 141.0000g for 1 hour at 4°C, to remove the cytosolic, mitochondrial and peroxisomal fraction. The isolated autophagosomal and endoplasmic reticulum material was diluted with an equal volume of HB buffer and overlaid on Nycodenz-Percoll gradients for centrifugation at 72000 xg for 30 min at 4°C. After centrifugation, to remove the Percoll silica particles, the AVs’ resulting interface was then diluted with 0.7 volumes of 60% Buffered Optiprep overlaid by 30% Buffered Optiprep and HB buffer. The Optiprep gradients were then centrifuged at 71000g for 30min at 4°C. The collected AVs were diluted in three volumes of HB buffer and centrifuged for 30 min at 16000g at 4°C. The concentration of the purified isolated AVs was then measured by BCA, following the manufacturer’s instructions. Purified AVs were treated with Proteinase K (PK) (20ng/μl) on ice for 20 min, in the presence or absence of 1% Triton X-100, and then 4 mM of PMSF was added for 10 min on ice for PK inactivation. The samples were then centrifuged at 16.900 xg for 10 min at 4°C, and the autophagosomal pellets were resuspended in Laemmli buffer, boiled for 5 min at 95°C and analyzed by Immunoblotting.

### Electrocorticogram recordings

Telemetric electrocorticogram analyses were performed using implantable radio transmitters (models TA11EA-F10 or TA11ETA-F10, DataSciences International). Adult (at least 3 months old) male mice received 0.05 mg/kg buprenorphine (subcutaneously) for analgesia and were anesthetized with 1.5–2% isoflurane in 100% oxygen. Midline skin incisions were made on the top of the skull and in the neck. The transmitter body was implanted subcutaneously. The tips of the leads were placed 2 mm posterior to bregma and 1.8 mm right to the midline for the recording electrode, and 1 mm posterior to lambda and 1–2 mm left to the midline for the reference electrode. The electrodes were fixed with dental cement. Radio transmitters allowed simultaneous monitoring of electrocorticogram and motor activity in undisturbed, freely moving mice housed in their home cages. Telemetry data and corresponding video data were recorded 48-72 hrs after surgery and were stored digitally using Ponemah software (Data Sciences International). Electrocorticogram recordings lasted over 50 h, sampled at 500 Hz, with continuous video recording.

### Recording of network activity of primary neurons using microelectrode arrays (MEA) system

Cortico-hippocampal neurons from postnatal *Atg5*flox:flox mice were isolated as described before (*17*) and plated in 24-well plates with gold electrodes on epoxy (24W700/100F-288, Multi Channel Systems). At DIV5, the KO was induced by transduction with *CamKIIα*-Cre (AAV9.CamKII.HI.eGFP-Cre.WPRESV40, Penn Vector Core). *CamKIIα*-eGFP (AAV9.CamKII0.4.eGFP.WPRE.rBG, Penn Vector Core) was used for the control group. MEA recordings were acquired at DIV15 using the Multiwell-Screen software of the Multiwell-MEA-System (Multi Channel Systems). Recordings were taken for 3 min with a sampling rate of 20000 Hz and automatic threshold estimation. The analysis was performed with the Multiwell-Analyser software (Multi Channel Systems) using default parameters if not otherwise stated. Per well, the best channel electrode was selected for analysis. A well was considered active when at least 3 active electrodes were detected in the well (out of 12).

### Immunohistochemical analysis of brain sections

Mice at 12-15-week were euthanized by an overdose of 1.2% ketamine, and 0.16% xylazine in PBS (i.p., 10 μl per 10 g body weight) and transcardial perfusion was performed as previously described (*17*). Brains were carefully dissected and postfixed in 4% PFA overnight before being processed for immunohistochemistry as previously described (*17*). Corresponding horizontal 40μm sections from WT and KO littermates were washed three times in PBS (3 × 5 min each). Sections were blocked with 10% normal goat serum (NGS) or normal donkey serum (NDS) in 0.5% PBS-T for 1 hour at RT. Primary antibodies (Table S8) were incubated on sections in 3% NGS/NDS and 0.3% PBS-T overnight at 4°C. Sections were washed three times 10 min in 0.3% PBS-T before incubation in fluorescence labelled secondary antibodies in 3% NGS/NDS and 0.3% PBS-T for 1 hour at RT protected from light (see Tables S8, S9). The sections were imaged at a Leica SP8 confocal microscope (Leica Microsystems) equipped with a 63x/1.32 oil DIC objective and a pulsed excitation white light laser (WLL; ~80-ps pulse width, 80-MHz repetition rate; NKT Photonics).

### 3D-analysis and reconstruction

Samples were scanned using Plan-Apochromat 63×/1.32 Oil DIC objective at a resolution of 1,024 × 1,024 pixels with 8-bit sampling in sequential scanning frame-by-frame mode. Single optical sections were acquired using identical acquisition settings, with the pinhole of 1 Airy Unit. Stacks of 8–29 optical sections yielded voxel dimensions between 100 and 300 nm for the X, Y and Z planes. 3D reconstructions were generated with Amira Software 2020.2 (Thermo Fisher Scientific). First, the surface area of the Bassoon-positive synaptic contacts was reconstructed using the Amira segmentation editor. Bassoon/PRK R1β-positive contacts were defined by color-coding the surface of PRK R1β cells found within 250 nm from Bassoon-positive varicosities. Subsequently, the surface of ‘250-nm-distant’ PRK R1β-positive voxels was mapped onto Bassoon-positive contact using the “surface distance” tool and plotted as a histogram.

### Mouse embryonal fibroblasts (MEF)

MEF cells were maintained in DMEM medium (GIBCO), supplemented 10% FCS (Sigma), 1% Pen Strep (Gibco), 1% MEM NEAA (Gibco). Cells were transfected 24-48 hours after seeding, when the culture reached a confluency of approximately 50-70%, using Lipofectamine™ 3000 Transfection Reagent (Invitrogen) following manufacturers’ guidelines using 5μg Plasmid DNA. After 48 hours, the cells were harvested in RIPA buffer for immunoblotting analysis. PKA signaling was analyzed by detecting pCREB protein level after forskolin stimulation (5 min at 37°C with 10μM Forskolin) or in DMSO treated cells as control.

### NSC34 cells

Cells were maintained in the same media as MEF cells (see above). To induce autophagy by Torin1, the media was supplemented with 250nM Torin1 or exchanged with starvation media (1.8 mM CaCl_2_, 0.8 mM MgSO_4_, 5.3 KCl, 26.2 mM NaHCO_3_, 117.2 mM NaCl, 1.0 mM NaH_2_PO_4_-H_2_O, 5.5 mM D-Glucose). To block lysosomal degradation 67 nM Bafilomycin A was added in the last 4 hours before harvesting for immunoblot analysis.

### Immunogold labelling on brain sections

Mice were perfused with warm Ringer solution, followed by 4% PFA before the brains were sliced at a vibratome (400 μm). Slices were briefly stained with 0.2% methylene blue in PBS buffer. Small blocks with visible CA1 pyramidal cell layer and adjacent Stratum Radiatum were cut out by the razor and placed into the 2.3 M sucrose in 0.1 M PB solution for 24 hours at 4°C. Following sucrose infiltration, blocks were placed at pins and plunge frozen in the liquid Ethan. Ultrathin (80 nm) cryosections were collected and thawed at sucrose drop (as described), placed at EM grids, blocked and washed in PBS supplemented with 1% BSA and 0.1% glycine, stained with R1 alpha or R1 alpha/beta antibodies 1:50. Following secondary antibody labelling 1:50 (12 nm gold goat anti-rabbit antibodies (Dianova), ultrathin sections were embedded and contrasted with 3% Tungstosilicic acid hydrate (w/v) in 2.8% polyvinyl alcohol (w/v), dried and imaged in Zeiss 900 transmission electron microscope. Images of CA1 Stratum radiatum were taken without prior knowledge of genotype or antibody labelling conditions. The reference area of subneuropil structures was determined by superimposing a grid over the neuropil images (volume fraction estimation). The number of gold particles found at those structures was normalized to the volume fraction of corresponding structures in the neuropil. No immunogold labelling was detected in samples where the PKA R1-α/β antibody were omitted (negative control). The preparation of neurons for postsynaptic density thickness detection was described previosly (*18*). Electron micrographs were taken with a JEM-2100 Plus Electron Microscope (JEOL), Camera OneView 4K 16 bit (Gatan), and software DigitalMicrograph (Gatan). The postsynaptic density of ultrathin section (70 nm) was measured using Image J.

### Co-immunoprecipitation experiments

For immunoprecipitation experiments, 20 μL Dynabeads Protein G (Thermo Fischer Scientific) were coated with 2 μg antibody targeting the protein of interest and corresponding IgG as a negative control (see Tables S8, S9). Therefore, the bead’s storing solution was replaced by 100 μL PBS, and 2 μg of antibody was added. The beads were incubated with the antibody for 2-3 hours at 4°C on a shaker before washing once with 200 μL PBS to remove excess antibodies. Neurons were harvested/ tissue was homogenized using a Wheaton Potter-Elvehjem Tissue Grinder in Co-Immunoprecipitation (Co-IP) buffer (50mM Tris-HCl pH = 7.4, 1% NP-40 / Igepal, 100mM NaCl, 2mM MgCl_2_) supplemented with Proteinase Inhibitor (Roche) und Phosphatase Inhibitor (ThermoScientific). Lysates were sonicated, incubated on ice for 45 min, and subsequently centrifuged at 13 000 for 20 min at 4°C. Protein concentration was assessed using Bradford assay (Sigma). An equal amount of protein was added to the beads coated with antibodies against the protein of interest and control IgG for overnight incubation at 4°C. The next day, lysates were removed, and beads were washed 3 times with Co-IP buffer before they were dissolved in 20 μL Co-IP buffer and 20 μL 4s SDS buffer and boiled at 95°C for 5 min. Precipitation of proteins was detected via SDS-page gel.

### Live-imaging of cultured neurons using GCAMP7 imaging

Calcium imaging experiments using GCAMP7^CamKII^ (ssAVV-9/2-mCaMKIIα-jGCaMP7f-WPREbGHp(A), Viral Vector Facility VVF) were performed in *Atg5*flox:tdTomato WT and KO primary cortico-hippocampal neuronal cell culture. The KO was induced as described above. Live-cell imaging and stimulation were performed at DIV15-16 using Zeiss Axiovert 200M microscope (Observer. Z1, Zeiss, USA) equipped with 10x/0.3 EC Plan-Neofluar objective; a pE-4000 LED light source (CoolLED), and a Hamamatsu Orca-Flash4.O V2 CMOS digital camera. Neurons were imaged in an osmolarity adjusted imaging buffer, and time-lapse images were taken within a 1-sec interval. For electrical field stimulation, neurons were stimulated four times with 100 action potentials (100 AP) in a 3-sec interval at 100 Hz using an RC-47FSLP stimulation chamber (Warner Instruments). Coverslips with *Atg5:CamKIIα*-Cre and *Atg5*flox:*Slc32a1*-Cre neurons were only stimulated once and were allowed to rest for 5 min before the buffer was supplemented with 10 μM CNQX for 5 min before imaging. WT neurons transfected with GCAMP7^CamKIIα^ (ssAVV-9/2-mCaMKIIα-jGCaMP7f-WPREbGHp(A), Viral Vector Facility VVF) were incubated with 10μM H89 in DMSO or DMSO as vehicle for 20 minutes before imaging. For analysis, neuronal cell bodies were chosen as ROI, and the GCAMP7 fluorescence response to the stimulus was plotted over time using Image J (Plot Z-axis Profile). Baseline fluorescence value was subtracted after background subtraction, and the traces were normalized to the first peak to visualize the facilitation. The 1^st^ and 4^th^ peak difference was calculated and presented in the bar graphs.

### Stereotactic injections for patch-clamp recordings

Stereotactic injections were performed on 10-week-old ATG5flox/flox mice. The mice were injected with *CamKII*-eGFP (AAV9.CamKII0.4.eGFP.WPRE.rBG, Penn Vector Core) as control or *CamKII*-eGFP-Cre (AAV9.CamKII.HI.eGFP-Cre.WPRESV40, Penn Vector Core) to induce acute ATG5 deletion in layer II/III pyramidal neurons. Mice were weighted and anaesthetized with a mixture of Ketamine (100mg/kg)/ Xylazine (20mg/kg)/ Acepromazine (3 mg/kg) and placed in a stereotactic frame when fully asleep. Local painkiller was injected subcutaneously at the operation field. The skin was opened, and the skull was cleaned using NaCl. Point injection (AP from Bregma: −1.82 mm, LM: +3.8mm) was identified using Bregma and Lambda for navigation. For injection, a small hole was drilled in the skull, and a 1 μL Hamilton syringe filled with the corresponding virus was lowered to the depth of −4.25mm to inject 150 nL over 5 min. This process has been repeated at a depth of −3.5mm to ensure sufficient virus-mediated transfection of neurons. Afterwards, the skull was rehydrated, the wound closed, and the mice were given a dose of carprofen to reduce postsurgical pain (s.c, 5 mg/kg) and 500 μL 5% glucose solution subcutaneously to enhance postoperative recovery.

### Measurements of AMPAR currents using perforated patch configuration and analysis

Whole-cell voltage clamp recordings of individual layer II/III pyramidal neurons were performed at ~32°C. Neurons were visualized with a fixed stage upright microscope (BX51WI; Olympus, Tokyo, Japan) equipped with infrared differential interference contrast optics and fluorescence optics. Layer II/III pyramidal neurons were identified by their anatomical location and eGFP-fluorescent label. Electrodes with tip resistances between 3 and 5 MΩ were fashioned from borosilicate glass (0.86 mm inner diameter; 1.5 mm outer diameter; GB150-8P; Science Products) with a vertical pipette puller (PP-830; Narishige, London, UK). All recordings were performed using a dSEVC (discontinuous single-electrode voltage-clamp) amplifier (SEC-05X, npi Electronics GmbH, Tamm, Germany) controlled by the program Patch-Master (version 2.32; HEKA). AMPA-R currents were recorded in discontinuous voltage clamp mode. The switching frequency and duty cycle of the amplifier were set to 40 – 50 kHz and 1/4 (current injection/potential recording), respectively Data were low-pass filtered at 3 kHz with a four-pole Bessel filter and sampled at 10 kHz. Data were recorded using a micro1410 data acquisition interface and Spike 2 (version 7) (both from CED, Cambridge, UK). Patch pipettes were filled with the following (in mM): 140 KCl, 10 HEPES, 0.1 EGTA, 5 MgCl_2_, 5 K-ATP, 0.3 Na-GTP, adjusted to pH 7.3 with KOH). Neurons were voltage clamped at −70 mV. The calculated liquid junction potential (3.6 mV) was compensated. To isolate AMPA receptor mediated PSCs, GABAergic inhibitory PSCs were blocked with picrotoxin (100 μM; Sigma) and NMDA receptor mediated PSCs were blocked with DL-2-amino-5-phosphonopentanoic acid (D-AP5; 50 μM; Sigma). The AMPA-R PSC frequency and amplitude was determined from a 1 min interval after the recording had stabilized (~10 min after wash-in of picrotoxin/D-AP5). The data were digitally filtered offline using the smooth (1 msec time period) and the DC remove (500 msec time period) functions in the Spike2 software (CED, Cambridge, UK). Data analysis was performed with Spike2 (version 7; Cambridge Electronic Design Ltd., Cambridge, UK) and the Pandas software library for Python 3.x (https://pandas.pydata.org/). Estimation statistics and plots were generated with the DABEST library for Python 3.x. The final figure was made with Adobe Illustrator 2022 (Adobe, San Jose, CA, USA) running on an Apple Macintosh.

### Fluorescence-activated cell sorting (FACS) of isolated neurons

Neurons for FACS sorting were isolated from 3-week-old *Atg5*flox:*CamkIIα*-Cre:tdTomato and *Atg5*flox:*Slc32a1*-Cre:tdTomato reporter mice. Mice were transcardially perfused with ACSF to remove blood cells. Isolated brains were dissected in HibernateA (HA, Gibco) and cortex, hippocampus and striatum were isolated and transferred into a fresh dish containing HABG (Hibernate-A, 2% B27, 1% GlutaMax). The tissue was cut into pieces and digested in activated Papain (Worthington) for 40 min at 37°C. Afterwards, tissue pieces were transferred back into fresh HABG. They were homogenized with a fire-polished Pasteur pipette (triturate approximately ten times in 1 min). The cell suspension was applied on an OptiPrep®Density Gradient Medium (Sigma) centrifuged at 800 xg for 15 min at 22°C to force cell type separation. Neuron enriched fractions were maintained for further processing. Gradient material was diluted with 10 mL HABG and cells were pelleted down at 3000 xg for 3 min at 22°C to clean cell suspension from debris contaminations. This step was repeated once before the cells were resuspended in NBA supplemented with 2% B27. To obtain purified lysates from autophagy-deficient neurons, cells in suspension were stained with DAPI and DRAQ5 ((see Tables S8, S9). Cell sorting was performed using BD FACSAria Fusion IIu and BD FACSAria IIIu with FACSDiva 8.0.1 software. Neurons were sorted at 4°C using a 100 μm nozzle and sheath pressure was set at 20 psi. 0.9% NaCl was used as sheath fluid. The tdTomato highly positive/ DAPI negative/ DRAQ5 positive cell population was selected. Cells were collected in chilled 1.5 mL Eppendorf tubes containing DPBS. After sorting, the cells were centrifuged at 3000 xg for 3 min and lysed in buffers depending on subsequent analysis.

### RNA Sequencing

RNA of FACS sorted neurons were isolated using the RNeasy® Plus Micro Kit (Qiagen) following manufacturer’s instructions. Isolated RNA was forwarded to RNA sequencing. Library preparation was performed with the NEBNext® Ultra Directional RNA Library Prep Kit for Illumina. The first step involves the removal of ribosomal RNA using biotinylated target-specific oligos combined with rRNA removal beads from 1ug total RNA input. The NEBNext® rRNA Depletion Kit (Human/Mouse/Rat) depletes samples of cytoplasmic rRNA. Following purification, the RNA is fragmented into small pieces using divalent cations under elevated temperatures. The cleaved RNA fragments are copied into first-strand cDNA using reverse transcriptase and random primers, followed by second-strand cDNA synthesis using DNA Polymerase I and RNase H. These cDNA fragments then have the addition of a single‘A’ base and subsequent ligation of the adapter. The products are purified and enriched with PCR (20ul template, 15cycles) to create the final cDNA library. After library validation and quantification (Agilent Tape Station), equimolar amounts of the library were pooled. Pools of 4-6 libraries were quantified using the Peqlab KAPA Library Quantification Kit and the Applied Biosystems 7900HT Sequence Detection System and sequenced on a NovaSeq 6000 S4 flowcell 2×100bp. Bioinformatic analysis of RNA sequencing raw data processing can be found in the supplementary material. The p-value of the Kallisto gene counts was calculated, and all significant hits were forwarded to analysis. The means were calculated between the genotypes and the genes sorted by the difference between WT and KO. The dataset was divided into up- and downregulated hits and both datasets were analyzed using Enrichr (Ma’ayan Lab, Icahn School of Medicine at Mount Sinai, New York) gene set analysis. Results from ChEA 2016 analysis are presented as Clustergrams.

### High content screening microscopy

Cells were isolated from adult 12-13-week-old mice as described for FACS sorting. Afterwards, the cells were resuspended in supplemented NBA medium, were seeded 100 μl/well in clear 96-well plates (Greiner #655896) and were incubated for 3 hours (37 C, 5% CO2) before stimulation. Using an electronic 8-channel pipette (Eppendorf) at low dispense speed, the cells were stimulated for 15 min. The stimulant compound was prepared 10-fold concentrated in 12.5 μl of PBS (PAA #H15-002) in 96-well V-bottom plates (NerbePlus, #10-111-000). Half of the media (50 μl) was removed from the wells in the cell culture plate, mixed with a stimulant in the V-bottom plate, and added back to the well. Negative controls were stimulated with the vehicle only. The cells were fixed by adding 100 μl 8% PFA (4% final) for 10 min followed by 2X wash with 100 μl PBS. Then the cells were blocked and permeabilized with 50 μl normal goat serum blocking (NGSB) for 1 h at room temperature. Afterwards, the cells were incubated with 30 μl primary antibodies diluted in 1% BSA overnight at 4°C. Rinsed 3X with 100 μl PBS for 10 min each and were incubated with 50 μl secondary antibodies diluted 1:1000 in PBS for 1 h at room temperature in the dark and rinsed 3X with 100 μl PBS for 10 min each ((see Tables S8, S9). Finally, the wells were filled with 200 μl PBS and sealed with aluminum. For imaging, a CellInsight CX7 LZR (ThermoFisher Scientific) microscope with a laser light source. We used the 20X objective and acquired images of 1024 * 1024 pixels. Image analysis was performed using the Cellomics software package. Briefly, images of tdTomato stainings were background corrected (low pass filtration), converted to binary image masks (fixed threshold), and segmented (geometric method). Inhibitory neurons were then identified by appropriate object selection parameters such as size, circularity, and average tdTomato intensity. These image masks were then overlaid on images obtained at other fluorescence wavelengths to quantify signal intensities. To later determine spillover between fluorescence channels, three respective controls for each triple staining: (1) tdTomato alone, (2) tdTomato + antibody 1, and (3) tdTomato + antibody 2 were included. Single-cell data containing the raw fluorescence data of single cells from the Cellomics software was exported as spreadsheets and analyzed by open-source statistical language R. The raw fluorescence data of the controls were used to calculate the slope of best fit straight lines by linear regression, which were then applied to compensate. Compensated data were scaled to a mean value of 1 for the vehicle-stimulated cells to adjust for variability between experimental treatments. To visualize the data, we used two-dimensional probability density plots generated using R. For gating of subpopulations, we set thresholds at local minima of probability density plots.

### Proteome analysis using LC-MS/MS

For total proteome analysis *Atg5*flox:CAG-Cre^Tmx^ and Atg16L1flox:CAG-Cre^Tmx^ cultured cortico-hippocampal neurons were harvested at DIV15-16 in RIPA buffer and processed as described for Immunoblotting (see above). A total of 50 μg were forwarded for proteomic analysis using an in-gel digestion protocol for sample preparation. In brief, samples were loaded onto an SDS-PAGE gel and run for 1-2 cm. The whole running line was chopped into small pieces and transferred in a 1.5 mL Eppendorf tube. Samples were reduced (10 mM DTT) and alkylated (55 mM CAA). Digestion of proteins were done overnight using Trypsin (10 ng/μL 90% Trypsin) and LysC (50 mM). Peptides were extracted and purified using Stagetips. Eluted peptides were dried in vacuo, resuspended in 1% formic acid/4% acetonitrile and stored at −20°C prior MS measurement. For proteome analysis of FACS-sorted neurons, the cells were pelleted and resuspended in SP3 lysis buffer (5% SDS in PBS). Chromatin was degraded using a Biorupter (10 min, cycle 30/30). The samples were reduced (5 mM DTT) and alkylated (40 mM CAA) and protein content was measured with the Pierce BCS protein assay. 20 μg were used for protein digestion according to the single-pot solid-phase-enhanced sample preparation (Hughes et al. 2019). Precise description of LC-MS/MS and data analysis for each dataset can be found in the supplementary material. Link to the data is provided in the data availability section.

### Phosphoproteome analysis using LC-MS/MS

Experiments were performed with *Atg5*flox:*CamKIIα*-Cre and *Atg5*flox:*Slc32a1*-Cre WT/KO 12-14-week-old brain tissue. Brains from SILAC-labelled mice (*27*), and ATG5 KO mice of both lines were homogenized using a Wheaton Potter-Elvehjem Tissue Grinder (VWR) in 6 M Urea/ 2 M Thiourea. Lysates were sonicated and centrifuged at maximum speed. The supernatant was forwarded to protein concentration assessment using Pierce™ 660nm Protein Assay Reagent (Thermo Scientific™). Lysates of ATG5 KO mice were mixed with brain lysates of isotope labelled mice in a ratio of 1:1 (1 mg:1 mg). Samples were incubated with 10 mM DTT for 30 min at RT, followed by 45 min incubation with 55 mM IAA in the dark. Afterwards, samples were pre-digested with LysC in an enzyme:substrate ratio of 1:50 for 2 hours. The samples were diluted three times with 50 mM ABC buffer to reach a concentration of 2 M Urea before LysC was added again in an enzyme:substrate ratio of 1:100 for overnight digestion at RT. The next day, desalting and phosphopeptide enrichment were performed. Experiments with acute brain of slices of *Atg5*flox:*CamKIIα*-Cre WT and KO treated with the cAMP-inducing agent Forskolin were performed with 12-14 weeks old mice. The brain was isolated and chopped into 400μm thick sections. The sections were washed with HBSS, transferred into media, and placed in the incubator for 1 hour. Afterwards, they were treated with 50μM Forskolin for 15 min before being washed with ice-cold PBS for albumin removal. The sections were snap-frozen and stored at −80°C till further use. On the day of the experiment, slices were homogenized in 1.5 mL 8 M Urea buffer using a Wheaton Potter-Elvehjem Tissue Grinder (VWR). The lysates were sonicated and centrifuged at 13000 rpm for 10 min. Protein concentration was assessed using Pierce™ 660nm Protein Assay Reagent (Thermo Scientific™) and 1 mg protein of each sample taken for the experiment. The samples were predigested in 5 mM TCEP, 27.5 mM CAA and LysC, in an enzyme:substrate ratio of 1:50 for 3 hours at RT after vortexing. Afterwards, the samples were diluted to a concentration of 2 M Urea using 50 mM ABC buffer. Trypsin was added to an enzyme:substrate ratio of 1:100 and incubated overnight at RT. The next day, desalting and phosphopeptide enrichment were performed. The next day, after Trypsin digestion, the samples of both datasets were acidified with TFA (1:100 [v/v]) before 10 min centrifugation. Desalting of the samples was performed with 50 mg C18 SepPak®Vac cartridges (Waters™). Therefore, the columns were activated with 2 mL 100% CAN and washed three times with 1 mL 0.1% TFA before loading samples. Samples were washed three times with 1 mL 0.1% TFA prior to elution with 0.6 mL elution buffer (60% ACN, 0.1% TFA). 30μg of the sample were taken, desalted, and stored on stage tips for total proteome analysis (described above) before the remaining sample was dried in a vacuum concentrator. TiO2 Phosphopeptide extraction was performed using the High-Select™ TiO2 Phosphopeptide Enrichment Kit (Thermo Scientific™) following the manufacturers’ instructions. In brief, a lyophilised peptide sample was resuspended in 150 μL Binding/Equilibration Buffer, columns prepared using 20 μL Wash Buffer followed by 20 μL Binding/ Equilibration Buffer. The samples were run twice through the column before the column was washed with 20 μL Binding/ Equilibration Buffer followed by 20 μL Wash Buffer. Afterwards, the columns were washed with 20 μL of LC-MS grade water before the phosphopeptides were eluted from the column in 50 μL Phosphopeptide Elution Buffer. Samples were dried immediately using a vacuum concentrator and subsequently resuspended in 15 μL resuspension buffer (5% FA and 2% ACN). Phosphopeptide samples were stored at −20°C until measurement. Precise description of LC-MS/MS and data analysis for each dataset can be found in the supplementary material. Data availability described in data availability section.

### Pathway analysis

Pathway analysis of proteomic approaches was performed using WebGestalt (WEB-based GEne SeT AnaLysis Toolkit, developed and maintained by the Zhang Lab, Baylor College of Medicine, Houston, Texas) and ShinyGO v0.741: Gene Ontology Enrichment Analysis (http://bioinformatics.sdstate.edu/go/). Venn diagrams were created using Gene List Venn Diagram, an interactive tool for comparing lists with Venn’s diagrams (http://genevenn.sourceforge.net/). GO functional analysis and human phenotype ontology analysis was done using g:Profiler (https://biit.cs.ut.ee/gprofiler/gost).

### Statistical analyses

Statistical analyses were done on cell values (indicated by data points) from at least 3 independent experiments (indicated by “n”, biological replicates). MS Excel (Microsoft, USA) and GraphPad Prism version 9 (GraphPad Software, Inc., USA) were used statistical analysis and result illustration (unless otherwise stated). Statistical analysis of normalized data between the two groups was performed using one-sample student’s t-test. Statistical significance between two groups for normally distributed non-normalized data was evaluated with a two-tailed unpaired student’s t-test. Two-tailed unpaired Mann-Whitney test was used for analysis between two groups for non-normally distributed non-normalized data. Statistical difference between more than two groups and two conditions was evaluated using Two-Way ANOVA (Tukey posthoc test was used to determine the statistical significance between the groups). Statistical difference between more than two groups was calculated using One-way ANOVA with Dunnett’s multiple comparison test. One-way ANOVA with Dunn’s multiple comparison (Kruskal-Wallis test) was used for comparison between more than two groups for not normally distributed data. In experiments using the GCAMP probe, ROUT-based outlier analysis was applied to all data sets before statistical analysis. Significant differences were accepted at p<0.05 indicated by asterisks: * p<0.05; ** p<0.01; *** p<0.001.

## Acknowledgements

We thank Marvin Schäfer for expert technical assistance. We are indebted to Dr. Ulrike Göbel (CECAD Bioinformatics facility), Dr. Christian Jüngst (CECAD Imaging Facility), Dr. Stephan Müller and Dr. Jan-Wilm Lackmann (CECAD Proteomic Facility) for their help and expert assistance. We thank Prof. Matteo Bergami (Univ. of Cologne) for providing the Ai9(RCL-tdT) transgenic mice and Prof. Matthew Poy (Johns Hopkins University School of Medicine, USA) for providing *Slc32a1*-Cre mice. We are indebted to Dr. Paul Turko (Charité Belrin) for a valuable advice on FACS sorting protocol. Flow cytometry experiments were performed with help of the FACS & Imaging Core Facility at the Max Planck Institute for Biology of Ageing. We thank Eike Strathmann (Institute of Human Genetics, Medical Facultyand University Hospital Cologne, University of Cologne) for the help with the pilot RNAseq data visualization, as well as Martina Ringling (FMP Core cellular imaging facility, Berlin) for excellent technical assistance with EM data processing. We thank Dr. Janine Altmüller and the Cologne Center for Genomics (CCG) for the help with NGS sequencing.

## Author contributions

Conceptualization: MO, NLK

Methodology: MO, NLK, DI, TH, MK, PK, MF, VN, ADV, JI

Investigation: MO, FT, SH, JT, MT, MF, MM, EDB, EK, VN

Visualization: MO, FT, SH, JT, MT, MF, MM, EDB, EK

Supervision: MK, PK, NLK, TH, BW, VN

Writing—original draft: MO, NLK

Writing—review & editing: MO, NLK

## Data and materials availability

All data needed to evaluate the conclusions in the paper are present in the paper and/or the Supplementary Materials. Proteome data of all experiments are deposited in the database PRIDE and accessible for public after publishing. Additional data related to this paper may be requested from the corresponding author.

