## Supplementary material for "Autophagy regulates neuronal excitability by controlling cAMP/Protein Kinase A signaling"

**This PDF file includes:**

Supplementary Text
Figs. S1 to S6
Table1 S5-S9
Table description S1 to S9
Movies S1 description

**Other Supplementary Materials for this manuscript include the following:**

Tables S1-S4
Movies S1
Data S1
Data S2

### Supplementary Text

### Materials & Methods

#### Primary mouse neurons

*Neuronal cell culture.* Cortical and hippocampal neurons from postnatal pups (P1-5) were isolated as previously described (*18*) and plated on PDL-coated coverslips or dishes. Homologous recombination in neurons expressing *Atg5*flox:*CAG-Cre*$^{Tmx}$ was induced by applying 0.4 µM (Z)-4-hydroxytamoxifen (Sigma) immediately after plating. After 24 and 48 h, cells were treated with 0.2 µM and 0.4 µM of tamoxifen, respectively, during medium renewal. Ethanol was added to control neurons (WT) equal to tamoxifen.

*Plasmid transfection of primary neurons.* Neurons were transfected at DIV7 via calcium phosphate transfection using the ProFection Mammalian Transfection System- Calcium Phosphate Kit (Promega) following manufacturers' guidelines. In brief, plasmid DNA was mixed with 2 M $CaCl_2$, water and HEPES buffer (see Table S6). Neurons were covered with transfection solution in osmolarity adjusted NeurobasalA medium (Gibco), washed and transferred back into their original medium. Neurons were used at DIV14-16.

*Nutrient deprivation of primary neurons.* Neurons were starved by replacing regular culturing media with homemade, osmolarity-adjusted starvation media (1.8 mM $CaCl_2$, 0.8 mM $MgSO_4$, 5.3 KCl, 26.2 mM $NaHCO_3$, 117.2 mM NaCl, 1.0 mM $NaH_2PO_4$-$H_2O$, 5.5 mM D-Glucose). Cells were starved overnight (16 hours) and harvested/ PFA fixated the following day. Bafilomycin A was supplemented in a concentration of 67 nM, H89 dihydrochloride was used in a 1 µM concentration in the media.

*AAV-mediated transduction of primary neurons.* AAVs (see Table S7) were added to the cell media at DIV5. For induction of GCAMP7$^{CamKII\alpha}$ expression, this AAV (ssAVV-9/2-mCaMKIIα-jGCaMP7f-WPREbGHp(A), Viral Vector Facility VVF) was added to the cell culture at DIV7.

*Immunocytochemistry on cultured neurons.* Neurons were fixed in 4% PFA/sucrose in phosphate-buffered saline (PBS) for 15 min at room temperature (RT) at DIV 14-16 and processed as described previously (*17*) (see Tables S8, S9 for the list of antibodies). Fixed neurons were imaged using either Zeiss Axiovert 200M microscope equipped with 40x/1.4 oil DIC objective and the Micro-Manager software (Micro-Manager1.4, USA) or with Leica SP8 confocal microscope (Leica Microsystems) equipped with a 63x/1.32 oil DIC objective and a pulsed excitation white light laser (WLL; ~80-ps pulse width, 80-MHz repetition rate; NKT Photonics).

*Colocalisation analysis.* Colocalization of AMPA receptor subunit GLUR1 and postsynaptic density marker protein PDS95 were assessed in *Atg5*flox:*CAG*-Cre$^{Tmx}$ neurons in a semi-automated fashion using a macro-tool. For colocalization analysis an ROI were set in an area devoid of somata. Each channel was individually smoothened and binarized with a user-defined threshold before calculating the overlap area by dividing the overlap between PSD95 and GLUR1 by total GLUR1 signal (custom written Image J script is available as Data S1).

*Analysis of pCREB positive neurons after starvation.* The percentage of pCREB positive neurons in different culture media was calculated from DAPI positive cells.

*Live-imaging of cultured neurons using GLUR1-pHlourin.* Neurons were imaged 7-8 days after transfection at DIV14-15 using Zeiss Axiovert 200M microscope (Observer. Z1, Zeiss, USA) equipped with 40x/1.4 oil DIC objective; a pE-4000 LED light source (CoolLED), and a Hamamatsu Orca-Flash4.O V2 CMOS digital camera. Time-lapse images were acquired every second using Micro-Manager software (Micro-Manager1.4, USA). The imaging chamber was connected to aVC-6 (six-channel valve controller, Warner Instrument Corporation) and plugged into a pump (Mini-Peristaltic Pump II, Harvard Apparatus) to ensure a constant media flow over the neurons. Note that all used media were osmolarity adjusted using D-Mannitol before the experiment. Neurons were imaged in imaging buffer (170 mM NaCl, 3.5 mM KCl, 0.4 mM $KH_2PO_4$, 20 mM N-Tris[hydroxyl-methyl]-methyl-2-aminoethane-sulphonicacid (TES), 5 mM $NaHCO_3$, 5 mM glucose, 1.2 mM $Na_2SO_4$, 1.2 mM $MgCl_2$, 1.3 mM $CaCl_2$, pH = 7.4) for 1 minute to establish baseline GLUR1-pHlourin level. Afterwards, media conditions were switched to an acidic buffer (pH = 5.5) to quench the GLUR1-pHlourin signal from membrane inserted receptors until a plateau was reached. For dequenching, the media was switched to a standard imaging buffer again. The total amount of GLUR1-pHlourin in the neurons was assessed by a media exchange to an alkaline buffer ($NH_4Cl$-buffer). To calculate the surface fraction of GLUR1-pHlourin receptors, 10 ROIs per neuron along the processes were chosen, and the mean fluorescence intensity signal was plotted over time using Image J (Plot Z-axis Profile). The background was subtracted for each single time point, and the mean of 30 images in the plateau phase of the single conditions was determined. The surface fraction was calculated as demonstrated in the formula for every single neuron:

$$Surface\ GLUR1\ (\%) = \frac{F(basal) - F(acidic)}{F(alkaline) - F(acidic)} * 100$$

### Preparation of acute slices for patch-clamp recordings

Experiments were performed on brain slices containing the perirhinal cortex from 12-week-old *Atg5*flox:flox mice, which were stereotactically injected either with a control virus AAV expressing eGFP or with *CamKIIα*-Cre-eGFP AAV (see above). Animals were kept under standard laboratory conditions, with tap water and chow available ad libitum, on a 12h light/dark cycle. The animals were lightly anesthetized with isoflurane (B506; AbbVie Deutschland GmbH and Co KG, Ludwigshafen, Germany) and decapitated. Coronal slices (300 µm) were cut with a vibration microtome (VT1200 S; Leica, Germany) under cold (4°C), carbogenated (95% O2 and 5% CO2), glycerol-based modified artificial cerebrospinal fluid (GaCSF) (*81*). GaCSF contained (in mM): 244 glycerol, 2.5 KCl, 2 $MgCl_2$, 2 $CaCl_2$, 1.2 $NaH_2PO_4$, 10 HEPES, 21 $NaHCO_3$, and 5 Glucose adjusted to pH 7.2 with NaOH. Brain slices were transferred into carbogenated artificial cerebrospinal fluid (aCSF). First, they were kept for 20 min in a 35°C 'recovery bath' and then stored at room temperature (24°C) for at least 30 min prior to recording. aCSF contained (in mM): 125 NaCl, 2.5 KCl, 2 $MgCl_2$, 2 $CaCl_2$, 1.2 $NaH_2PO_4$, 21 $NaHCO_3$, 10 HEPES, and 5 Glucose adjusted to pH 7.2 with NaOH. Slices were transferred to a recording chamber (~3 ml volume) and continuously superfused with carbogenated aCSF at a flow rate of ~2 $ml\cdot min^{-1}$.

**Cresyl-violet staining**

Horizontal brain sections were obtained from perfused 13-week-old Atg5flox:Slc32a1-Cre mice and processed for cresyl violet staining as previously described (*17*).

**Immunoblotting using brain lysates**

Mice were sacrificed at the age of 10-13-week via cervical dislocation. Brains were isolated, dissected, shock frozen in liquid nitrogen, and stored at -80°C till further use. Brain tissue was homogenized in radioimmunoprecipitation assay (RIPA) buffer (50 mM Tris pH 8.0, 150 mM NaCl, 1.0% IGEPAL CA-630, 0.5% Sodium deoxycholate, 0.1% SDS) containing Protease Inhibitor (Roche) and Phosphatase Inhibitor (ThermoScientific) using a Wheaton Potter-Elvehjem Tissue Grinder. Primary neurons, MEF cells, and NSC34 cells were harvested in RIPA buffer containing Protease Inhibitor (Roche) and Phosphatase Inhibitor (ThermoScientific) using a cell scraper. Samples were sonicated and incubated on ice for 45 min. Lysates were centrifuged at 13 000 rpm for 20 min at 4°C and supernatant concentration was assessed using Bradford assay (Sigma). 10-20 µg of protein per sample was loaded onto SDS-page gels for protein separation. Proteins were transferred to Nitrocellulose or methanol-activated PVDF membranes (LC3). Membranes were blocked for 1 hour with 5% skim milk (or BSA) in TBS (20 mM Tris pH=7.6, 150 mM NaCl) containing 1% Tween (TBS-T) before incubation with the primary antibodies overnight at 4°C (see Table S8). The next day, the membranes incubation with secondary HRP-tagged secondary antibody (see Table S9) for 1 hour at RT. Protein levels were visualized using an ECL-based autoradiography film system for documentation. Analysis was done using the Gel Analyzer plugin from Image J.

**Behavioral analysis**

The SmithKline, Harwell, Imperial College, Royal Hospital, Phenotype (SHIRPA) test was performed to determine behavioral or motoric alterations in WT and conditional ATG5-deficient mice (*82*). The mice were weighted and placed in a glass beaker (15 cm diameter, 19 cm height) for 2 minutes. During this time, the number of rears (Rearing), the level of arousal, and deviating external appearance characteristics were noted. Afterwards, the mice were transferred into an arena (55 cm x 33cm, 18cm height) in which the floor was subdivided into uniform squares. The number of squares passed during 1 minute was noted and displayed as locomotion (# of squares/min). After 1 minute, the response to a loud, spontaneous noise was measured and ranked from 0-2 (0 – no response; 1 – Preyer reflex; 2 –Reaction in addition to Preyer reflex) and displayed as startle response.

**LC-MS/MS acquisition and data analysis**

*Atg5*flox:*CAG*-Cre$^{Tmx}$ *cultured neurons*. All samples were analysed on a Q Exactive Plus Orbitrap (Thermo Scientific) mass spectrometer coupled to an EASY nLC (Thermo Scientific). Peptides were loaded with solvent A (0.1% formic acid in water) onto an in-house packed analytical column (50 cm — 75 µm I.D., filled with 2.7 µm Poroshell EC120 C18, Agilent). Peptides were chromatographically separated at a constant flow rate of 250 nL/min using the following gradient: 4-5% solvent B (0.1% formic acid in 80 % acetonitrile) within 1.0 min, 5-28% solvent B within 200.0 min, 28-50% solvent B within 28.0 min, 50-95% solvent B within 1.0 min, followed by washing and column equilibration. The mass spectrometer was operated in data-dependent acquisition mode. The MS1 survey scan was acquired from 300-1750 m/z at a resolution of 70,000. The top 10 most abundant peptides were isolated within a 2.0 Th window and subjected to HCD fragmentation at a normalised collision energy of 27%. The AGC target was set to 5e5 charges, allowing a maximum injection time of 55 ms. Product ions were detected in the Orbitrap at a resolution of 17,500. Precursors were dynamically excluded for 40.0 sec. Each sample was injected twice.
*Data analysis.* All mass spectrometric raw data were processed with Maxquant (version 1.5.3.8) using default parameters. Double injections from each sample were combined in MaxQuant by naming them identically. MS2 spectra were searched against the Uniprot mouse reference proteome containing isoforms (UP000000589, downloaded at 26.08.2020), including a list of common contaminants. The target-decoy approach estimated false discovery rates on protein and PSM levels to 1% (Protein FDR) and 1% (PSM FDR), respectively. The minimal peptide length was set to 7 amino acids, and carbamidomethylation at cysteine residues was considered a fixed modification. Oxidation (M), Phospho (STY), and Acetyl (Protein N-term) were included as variable modifications. The match-between runs option was enabled. LFQ quantification was enabled using default settings.

*Atg16L1flox:CAG-Cre$^{Tmx}$ cultured neurons*. Peptide digests were analyzed on a Q Exactive plus Orbitrap (Thermo Scientific) mass spectrometer coupled to an EASY nLC (Thermo Scientific). Samples were loaded onto an in-house packed analytical column (50 cm — 75 µm I.D., filled with 2.7 µm Poroshell EC120 C18, Agilent). Peptides were separated at a 250 nL/min flow rate using 2 hours runs with data-independent acquisitions (DIA) or 4 hours runs with the data-dependent acquisition (DDA). The gradients were: (2 hours) 3-5% solvent B (0.1% formic acid in 80 % acetonitrile) within 1.0 min, 5-30% solvent B within 91.0 min, 30-50% solvent B within 17.0 min,

50-95% solvent B within 1.0 min, followed by washing with 95 % solvent B for 10 min or (4h) 4-5% solvent B (0.1% formic acid in 80 % acetonitrile) within 1.0 min, 5-28% solvent B within 200.0 min, 28-50% solvent B within 28.0 min, 50-95% solvent B within 1.0 min, followed by washing with 95 % solvent B for 10 min. DDA runs for spectrum library generation were acquired from each sample. MS1 survey scans were acquired at a resolution of 70,000. The top 10 most abundant peptides were isolated within a 2.0 Th window and subjected to HCD fragmentation with a normalized collision energy of 27. The AGC target was set to 5e5 charges, allowing a maximum injection time of 55 ms. Product ions were detected in the orbitrap at a resolution of 17,500. Precursors were dynamically excluded for 20.0 s. Sample runs were acquired in DIA mode using 10 variable windows covering the mass range from m/z 450 to m/z 1200. MS1 scans were acquired at 140,000 resolution, and maximum IT restricted to 120 ms and an AGC target set to 5e6 charges. The settings for MS2 scans were 17,500 resolution, maximum IT restricted to 60 ms and AGC target set to 5e6 charges. The default charge state for the MS2 was set to 4. Stepped normalized collision energy was set to 27. All spectra were acquired in profile mode.
*Data analysis.* An assay-specific hybrid spectrum library was generated in Spectronaut 13 (*83*) using DDA library runs, DIA sample runs, and a mouse sequence file (up000000589) downloaded from Uniprot. Spectronaut default settings were used for the analysis of the DIA runs. Protein identifications were filtered for q-values below 0.01, and normalized intensities were exported for subsequent statistical analysis in Perseus 1.6.1.1 (*84*). Intensities were transformed to log2 values, and the dataset was filtered for at least 3 out of 3 values in at least one condition. The remaining missing values were imputed with random values from the left end of the intensity distribution (with 0.3 sd, downshift 2 sd). Two runs were removed from the analysis since the projection in the principal component analysis were outside median plus/minus 4 times inter quartile range for component 1 or 2. Two sample Student´s T-tests were calculated using permutation-based FDR estimation.

<u>*FACS-sorted neurons from Atg5*flox:*CamKIIα*-Cre/tdTomato mice.</u>
The LCMS analysis approach was adjusted to handle low input volume. Samples were analysed on a Q Exactive Exploris 480 (Thermo Scientific) mass spectrometer that was coupled to an Evosep ONE (Evosep) in the recommended “whisper” setup. Samples were loaded onto EvoTips following the manual instructions (Evosep). Peptides were chromatographically separated by the predefined “whisper 20 SPD” setup on a 58 min gradient on a PepSep 15 cm column with a 75 µm inner diameter filled with 1.9 µm Dr. Maisch resin. MS1 scans were acquired from 380 m/z to 900 m/z at 45k resolution. Maximum injection time was set to 60 ms and the AGC target to 100%. MS2 scans ranged from 400 m/z to 880 m/z and were acquired at 45 k resolution with a maximum injection time of 84 msec and an AGC target of 1000%. DIA scans covering the precursor range from 400 - 880 m/z were acquired in 15 x 30 m/z staggered windows resulting in 30 nominal 15 m/z windows after demultiplexing. All scans were stored as a centroid.
*Data analysis.* Thermo raw files were demultiplexed and transformed to mzML files using the msconvert module in Proteowizard. MzML files were converted to dia file format in DIA-NN

1.7.15. A Mouse canonical Swissprot fasta file was converted to a Prosit upload file with the convert tool in encyclopedia 0.9.0 (*85*) using default settings: Trypsin, up to 1 missed cleavage, range 396 m/z – 1004 m/z, charge states 2+ and 3+, default charge state 3 and NCE 33. The csv file was uploaded to the Prosit webserver and converted to a spectrum library in generic text format (*86*). The resulting library (16998 protein isoforms, 21694 protein groups and 1404872 precursors was used in DIA-NN 1.7.15 (*87*) to search acquired data in double-pass mode. The applied settings were: Output will be filtered at 0.01 FDR, N-terminal methionine excision enabled, Maximum number of missed cleavages set to 1, Min peptide length set to 7, Max peptide length set to 30, Min precursor m/z set to 400, Max precursor m/z set to 1000, Cysteine carbamidomethylation enabled as a fixed modification.

*Atg5*flox:*Slc32a1*-Cre/tdTomato

*LCMS data acquisition.* Samples were analysed on a Q Exactive Exploris 480 (Thermo Scientific) mass spectrometer equipped with a FAIMSpro differential ion mobility device coupled to an EASY-nLC 1200 (Thermo Scientific). Samples were loaded onto an in-house packed analytical column (30 cm — 75 µm I.D., filled with 2.7 µm Poroshell EC120 C18, Agilent). Peptides were chromatographically separated at a constant flow rate of 300 nL/min and the following gradient: initial 4% B (0.1% formic acid in 80 % acetonitrile), up to 30% B in 74 min, up to 55% B within 8.0 min and up to 95% solvent B within 2.0 min, followed by a 6 min column wash with 95% solvent B. The FAIMS pro was operated at -47V compensation voltage and electrode temperatures of 99.5 °C for the inner and 85 °C for the outer electrode. Identical HPLC settings were used for library generation, and sample runs.

*Spectrum library generation by Gas-phase fractionation.* Aliquots from each sample were pooled, and the pool was used for spectrum library generation by narrow window DIA of six 100 m/z gas phase fractions (GPF) covering the range from 400 m/z to 1000 m/z (*88*). The Orbitrap was operated in DIA mode. MS1 scans of the respective 100 m/z gas-phase fraction were acquired at 60k resolution. Maximum injection time was set to 60 ms and the AGC target to 100%. MS2 scans of the corresponding 100 m/z regions were acquired in 24 x 4 m/z staggered windows resulting in 48 nominal 2 m/z windows after demultiplexing. MS2 settings were 30 k resolution, 60 ms maximum injection time and an AGC target of 100%. All scans were stored as a centroid.

*Data independent acquisition of samples.* MS1 scans were acquired from 390 m/z to 1010 m/z at 60k resolution. Maximum injection time was set to 60 ms and the AGC target to 100%. MS2 scans ranged from 300 m/z to 1500 m/z and were acquired at 15 k resolution with a maximum injection time of 22 msec and an AGC target of 100%. DIA scans covering the precursor range from 400 - 1000 m/z were acquired in 50 x 12 m/z staggered windows resulting in 100 nominal 6 m/z windows after demultiplexing. All scans were stored as a centroid.

*Data analysis.* Thermo raw files were demultiplexed and transformed to mzML files using the msconvert module in Proteowizard. MzML files were converted to dia file format in DIA-NN 1.7.11.

*Spectral Library.* A Mouse canonical Swissprot fasta file was converted to a Prosit upload file with the convert tool in encyclopedia 0.9.0 (*85*) using default settings: Trypsin, up to 1 missed cleavage, range 396 m/z – 1004 m/z, charge states 2+ and 3+, default charge state 3 and NCE 33. The csv file was uploaded to the Prosit webserver and converted to a spectrum library in generic text format (*86*). The resulting library (16998 protein isoforms, 21698 protein groups and 1404872 precursors) was searched in DIA-NN 1.7.11 (*87*) with the 6 GPF runs to generate a project-specific library (6275 protein isoforms, 6523 protein groups and 26215 precursors). The applied settings were: Output will be filtered at 0.01 FDR, N-terminal methionine excision enabled, the maximum number of missed cleavages set to 1, min peptide length set to 7, max peptide length set to 30, min precursor m/z set to 400, Max precursor m/z set to 1000, cysteine carbamidomethylation enabled as a fixed modification,
*Samples.* 10 sample files were searched with DIA-NN 1.7.11 and the project library. In addition to the settings used for library generation, Rt dependent normalization was used.

**Phosphoproteomic analysis**

*Atg5*flox:*CamKIIα*-Cre & *Atg5*flox:*Slc32a1*-Cre mice brain tissue. *LCMS data acquisition.* Proteome samples were eluted from stage tips with elution buffer (60% ACN, 1% ammonia), dried in a vacuum concentrator and resuspended in 12µl resuspension buffer. Proteome and phosphoproteome samples were measured on a Thermo Orbitrap Eclipse mass spectrometer with an AIMS interface coupled to a Thermo EASY-nLC system. Each sample was measured in a 90min reverse phase separation using two FAIMS CVs -45V and -65V.
*Data analysis.* All raw files were demultiplexed for FAIMS CVs using the Coons lab “FAIMS to Mzxml generator” (https://github.com/coongroup/FAIMS-MzXML-Generator). Mzxml files were analysed with MaxQuant 1.6.14.0 using a Uniprot mouse protein database (release July 3, 2020). Default settings were used plus: Multiplicity set to 2, label: lys6, enzyme: LysC/P, p(STY) enabled as variable modification, a match between runs were enabled, unmodified counterpart peptides were not discarded. The resulting MaxQuant (*89*) output was further analysed in Perseus 1.6.14.0 (*90*) and InstantClue 0.10.10 (*91*).

**Immunoprecipitation of PKA Substrates**

WT brains were homogenized in Co-IP buffer and processed as described above. A total amount of 2 mg protein was provided to the beads. The samples were loaded onto an SDS-PAGE gel and sample preparation followed in-gel digestion protocol (described above). Eluted peptides were dried in vacuo, resuspended in 1% formic acid/4% acetonitrile and stored at -20°C prior MS measurement. Precise description of LC-MS/MS and data analysis for each dataset can be found in the supplementary material. Data availability described in data availability section. All samples were analyzed on a Q Exactive Plus Orbitrap mass spectrometer that was coupled to an EASY nLC (both Thermo Scientific). Peptides were loaded with solvent A (0.1% formic acid in water) onto an in-house packed analytical column (50 cm, 75 µm I.D., filled with 2.7 µm Poroshell EC120

C18, Agilent). Peptides were chromatographically separated at a constant flow rate of 250 nL/min using the following gradient: 3-5% solvent B (0.1% formic acid in 80 % acetonitrile) within 1.0 min, 5-30% solvent B within 40.0 min, 30-50% solvent B within 8.0 min, 50-95% solvent B within 1.0 min, followed by washing and column equilibration. The mass spectrometer was operated in data-dependent acquisition mode. The MS1 survey scan was acquired from 300-1750 m/z at a resolution of 70,000. The top 10 most abundant peptides were isolated within a 1.8 Th window and subjected to HCD fragmentation at a normalized collision energy of 27%. The AGC target was set to 5e5 charges, allowing a maximum injection time of 110 ms. Product ions were detected in the Orbitrap at a resolution of 35,000. Precursors were dynamically excluded for 10.0 s. All mass spectrometric raw data were processed with Maxquant (version 1.5.3.8) using default parameters. Briefly, MS2 spectra were searched against the Uniprot mouse reference proteome containing isoforms (UP000000589, downloaded at 26.08.2020), including a list of common contaminants. False discovery rates on protein and PSM level were estimated by the target-decoy approach to 1% (Protein FDR) and 1% (PSM FDR) respectively. The minimal peptide length was set to 7 amino acids and carbamidomethylation at cysteine residues was considered as a fixed modification. Oxidation (M), Phospho (STY), and Acetyl (Protein N-term) were included as variable modifications. The match-between runs option was enabled. LFQ quantification was enabled using default settings.

**RNA sequencing data analysis**

*Read Trimming.* Illumina adapters were clipped off the raw paired-end reads using cutadapt v2.10 (*92*) with standard parameters and a minimum read length of 35 base pairs after trimming (shorter reads were discarded).

For read 1, the adapter sequences trimmed were
AGATCGGAAGAGCACACGTCTGAACTCCAGTCA,
AGATCGGAAGAGCACACGTCTGAAC, TGGAATTCTCGGGTGCCAAGG,
AGATCGGAAGAGCACACGTCT, CTGTCTCTTATACACATCT, and
AGATGTGTATAAGAGACAG.

For read 2, the sequences were AGATCGGAAGAGCGTCGTGTAGGGAAAGAGTGT,
AGATCGGAAGAGCGTCGTGTAGGGA, TGGAATTCTCGGGTGCCAAGG,
AGATCGGAAGAGCACACGTCT, CTGTCTCTTATACACATCT and
AGATGTGTATAAGAGACAG.

*Transcript Quantification.* Transcript abundance was quantified using kallisto v0.46.1 (*93*). The transcriptome index was built by kallisto index from file Mus_musculus.GRCm38.rna.fa of Ensembl release 100. Subsequently, the adapter-trimmed RNA-seq reads were matched against the index using kallisto quant with standard parameters and 100 bootstrap samples (-b 100). Transcript abundances (=TPMs, =estimated Transcripts Per Million) were extracted from the primary output of kallisto using package tximport v1.16.1 of the Bioconductor v3.11 (*94*) software project, in an environment of the R v4.0.0 programming language. In the same environment, Ensembl transcript ids were mapped to Ensembl gene ids and gene symbols, using package biomaRt v2.44.4 to access the Ensembl v100 database. A table containing the gene ids, gene symbols, and kallisto TPMs was output as kallisto_tx_abundance.xlsx (141450 transcripts). In table kallisto_tx_abundance_1TPM.xlsx, transcripts represented by less than 1 TPM were filtered out (72659 remaining transcripts).

For downstream gene-based analysis, the TPM values of transcript ids associated with the same Ensembl gene id were summed up. This resulted in 53028 unique Ensembl gene ids in the full matrix and 23669 unique Ensembl gene ids in the 1TPM-filtered matrix. The Ensembl ids were associated with 52263 unique gene symbols in the full matrix and 23380 unique gene symbols in the filtered matrix. To resolve cases where >1 Ensembl gene id mapped to the same symbol, we discarded Ensembl gene ids which were not represented in the Ensembl v100 genome annotation file Mus_musculus.GRCm38.100.gtf. (Note that unrepresented Ensembl gene ids were not removed if they did not cause a mapping conflict. As all unrepresented ids came from PATCH contigs rather than from the canonical chromosomes, this may be an issue. On the other hand, these contigs had been part of the kallisto index used and thus have had an influence on the mapping process, so they should be represented in the output.) After removal of 873 Ensembl gene ids due to conflicts, the final number of genes represented in the full matrix was 52155, and 23319 in the 1TPM-filtered matrix. The full and the 1TPM-filtered gene collapsed matrices are reported in files kallisto_gene_abundance.xlsx and kallisto_gene_abundance_1TPM.xlsx, respectively. In addition to the TPM values, kallisto also outputs estimated read counts per transcript. We converted these to read counts per gene by summing over associated transcripts as above, and retaining the same genes as in the gene-collapsed TPM matrices. Summed estimated counts are reported in files kallisto_gene_counts.xlsx and kallisto_gene_counts_1TPM.xlsx.

*Differential Gene Expression Analysis.* We used the R package sleuth v0.30.0 (*95*) for differential gene expression, because sleuth is explicitly designed to work with kallisto output. Because the experimental samples had been sequenced in two batches, we employed an additive linear model with categorical factors Batch and Genotype. Two of the original samples (BKC515 and BKC437 were left out of the analysis because their genotypic states were ambiguous. (Note that these samples are also left out of the reported kallisto TPM and count matrices.)
File newDGE_sleuth_BG_reproduced.xlsx reports the result of the sleuth analysis for model coefficient Genotype, that is, for the effect of genotype state (WT or KO) on transcript expression, corrected for batch membership of the samples.
Sleuth was configured to run Wald tests on the coefficients of the linear model and to not aggregate the p-values from the individual transcripts contributing to a gene's expression. Therefore, a given Ensembl gene id is usually represented in more than one row of the table. The original sleuth output column names "b" (effect strength of the model coefficient), "pval" (p-value = probability of the effect being equal to zero), and "qval" (FDR adjusted p-value using the Benjamini-Hochberg procedure) have been replaced by "logFoldChange", "p-value", and "FDR". (Note that the effect strength "b" is only roughly comparable to the natural logarithm of fold change, and that it's not

even clear whether in sleuth v0.30.0 the definition has not already been changed to log2 – see https://github.com/pachterlab/sleuth/issues/59). The reported table contains in addition columns for the Ensembl gene id ("geneid") and transcript id ("txid"), plus the gene symbol and a description.

#### cAMP Direct Immunassay

cAMP concentration in brain tissue of autophagy-defcient mice was measured using the cAMP Direct Immunoassays Kit (Flourometric) from abcam (ab138880) following the manufacturers instructions. Brains of *Atg5*flox:*CamKIIα*-Cre WT and KO mice were dissected at 12-weeks of age, snap-frozen and stored at -80°C. Hippocampi were weighted and homogenized in lysis buffer depending tissue weight using a Wheaton Potter-Elvehjem Tissue Grinder (VWR). Insoluble material was removed by centrifugation before supernatant was pipetted into the anti-cAMP coated 96-well plate. A standard with pre-defined cAMP concentrations was loaded in parallel. Total cAMP content was calculated using a set amount of HRP*-cAMP conjugate and HRP-dependent fluorescent substrate detected by a microplate reader at Ex/Em=540/590 nm.

### Supplementary figures

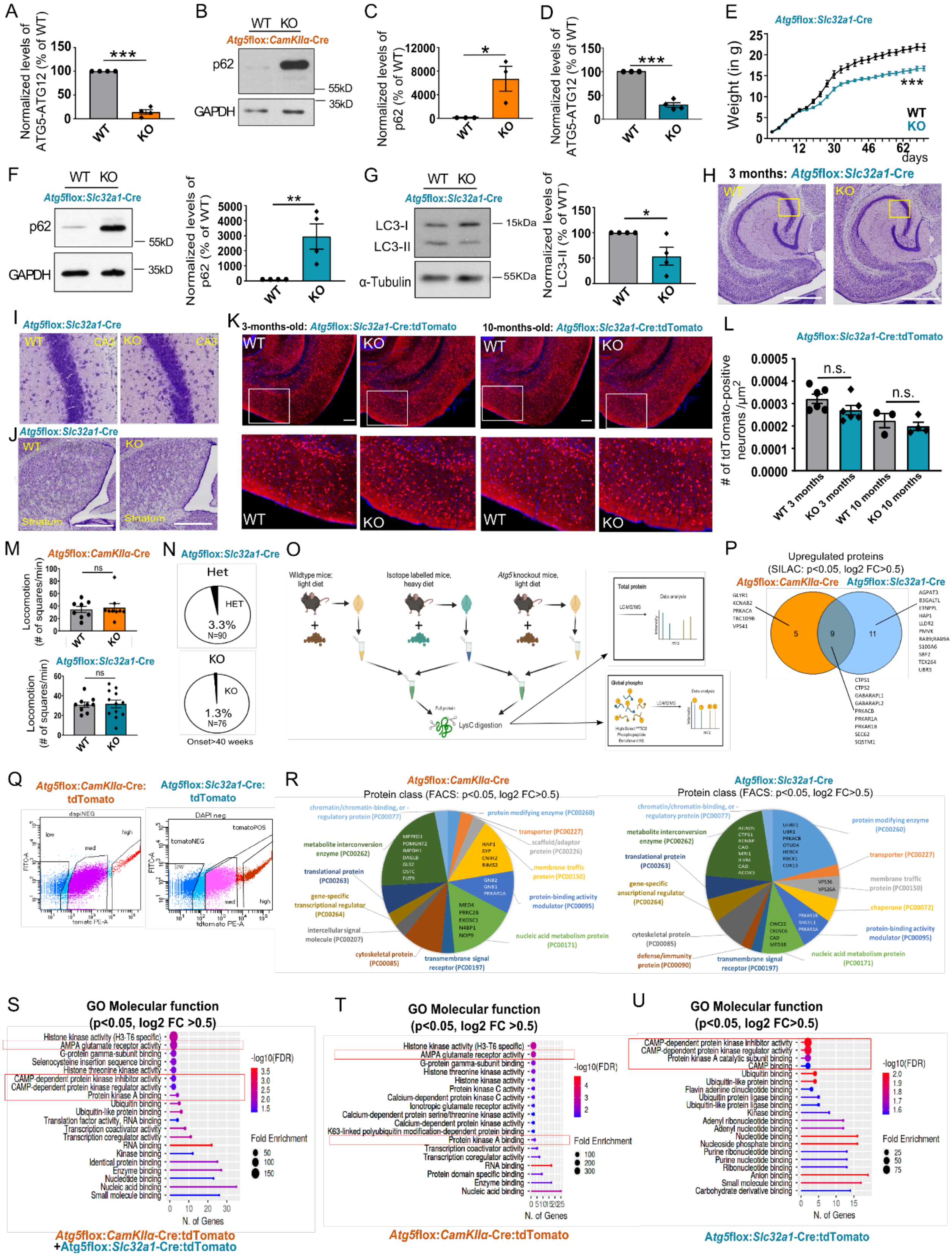

**Supplementary figure 1 | Survival-independent role of ATG5 in regulation of cAMP/PKA signaling in excitatory and inhibitory neurons.**

**(A)** Protein levels of ATG5 are significantly decreased in cortical lysates of *Atg5*flox:*CamKIIα*-Cre KO mice compared to the WT set to 100% (KO: 14.22± 4.80%). p<0.0001, n=4. Performed one-sample t-test. **(B,C)** Protein levels of p62 are significantly increased in cortical lysates of *Atg5*flox:*CamKIIα*-Cre KO mice compared to the WT set to 100% (KO: 6697± 2105%). p=0.0175, n=3. Performed one-sample t-test. **(D)** Protein levels of ATG5 are significantly decreased in striatal lysates of *Atg5*flox:*Slc32a1*-Cre KO mice compared to the WT set to 100% (KO: 29.70± 5.043%). p<0,0001, n=4. Performed one-sample t-test. **(E)** Bodyweight of *Atg5*flox:*Slc32a1*-Cre KO mice is decreased compared to WT littermates starting at 3 weeks of age (at 2 month of age; WT: 20.6± 0.68 g; KO: 15.6± 0.43 g). p=<0.0001, $n_{WT}$=18, $n_{KO}$=20. Performed unpaired t-test for each time point. **(F)** Protein levels of p62 are significantly increased in striatal lysates of *Atg5*flox:*Slc32a1*-Cre KO mice compared to the WT set to 100% (KO: 2956± 839.3%). p=0.0072, n=4. Performed one-sample t-test. **(G)** Protein levels of lipidated LC3 (LC3-II) are reduced in striatal lysates of *Atg5*flox:*Slc32a1*-Cre KO mice compared to the WT set to 100% (WT set to 100%, KO: 53.69 ± 17.69%). p=0.0199; n=4. Performed one-sample t-test. **(H-J)** Nissl staining of 40 µM thick horizontal brain sections reveals no signs of neuronal cell loss in 12-week-old *Atg5*flox:*Slc32a1*-Cre KO mice neither in the CA1 region of the hippocampus (H-I) nor in striatum (J). Scale bar: 200µm. **(K-L)** The number of tdTomato-expressing neurons was not altered in the entorhinal cortex (white rectangle) of *Atg5*flox:*Slc32a1*-Cre:tdTomato KO mice compared to the WT at 3 and 10 months of age ($WT_{3\ months}$: 0.00032± 0.00002079, n=6; $KO_{3\ months}$: 0.00027± 0.00002083, n=6; $WT_{10\ months}$: 0.00022± 0.00003156, n=3; $KO_{10\ months}$: 0.00020± 0.000018, n=4). Performed Two-Way ANOVA with Tukey's multiple comparisons test. Scale bar: 200µm **(M)** No alterations in locomotion (# of passed squares per 1 min) during SHIRPA analysis in *Atg5*flox:*CamKIIα*-Cre KO mice (WT: 34.38± 4.931, KO: 37.22± 6.606; $n_{WT}$=8, $n_{KO}$=9), as well as in *Atg5*flox:*Slc32a1*-Cre KO mice compared to the WT (WT: 30.89± 2.679, KO: 31.67± 3.938; $n_{WT}$=9, $n_{KO}$=12). **(N)** Percentages of *Atg5*flox:*Slc32a1*-Cre KO and HET mice that developed seizures at the age of 40 weeks and later. Animal number and the percentage of affected animals are displayed in the graphs (WT is set to 100%). **(O)** Schematic illustration of SILAC spike-in approach for global proteome and phosphoproteome analysis. Brain regions of interest from *Atg5*flox:*CamKIIα*-Cre (cortex) WT and KO mice, *Atg5*flox:*Slc32a1*-Cre (striatum) WT and KO mice and isotope-labeled mice (brain tissue) were dissected and homogenized. Unlabeled and isotope-labeled brain lysates were mixed in a 1:1 ratio and proteins were digested into peptides using LysC. Samples were divided to detect global proteome phosphoproteome alterations. **(P)** Venn diagram showing the number of highly upregulated proteins (log2FC>0.5, p<0.05) detected in SILAC spike-in global proteome analysis of *Atg5*flox:*CamKIIα*-Cre KO and *Atg5*flox:*Slc32a1*-Cre KO mice. **(Q)** Exemplary illustration of fluorescence-based cell sorting (FACS) of *Atg5*flox:*CamKIIα*-Cre:tdTomato and *Atg5*flox:*Slc32a1*-Cre:tdTomato neurons. The cells were categorized in three populations depending on their tdTomato fluorescence intensity expression. Only the populations of highly tdTomato-positive cells were used for subsequent proteomic analysis. **(R)** Pie chart of protein classes upregulated (p<0.05, log2FC>0.5) in *Atg5*flox:*CamKIIα*-Cre:tdTomato and *Atg5*flox:*Slc32a1*-Cre FACS-sorted KO neurons, analyzed using Panther classification system (http://www.pantherdb.org/panther/ontologies.jsp). **(S-U)** ShinyGO v0.741-based gene ontology (GO) analysis of "molecular function"-enriched terms in the proteome of either pooled *Atg5*flox:*CamKIIα*-Cre:tdTomato and *Atg5*flox:*Slc32a1*-Cre:tdTomato datasets (S) or in the dataset obtained from *Atg5*flox:*CamKIIα*-Cre:tdTomato (T) and *Atg5*flox:*Slc32a1*-Cre:tdTomato (U) mice separately (cut-off p<0.05, log2 FC>0.5).

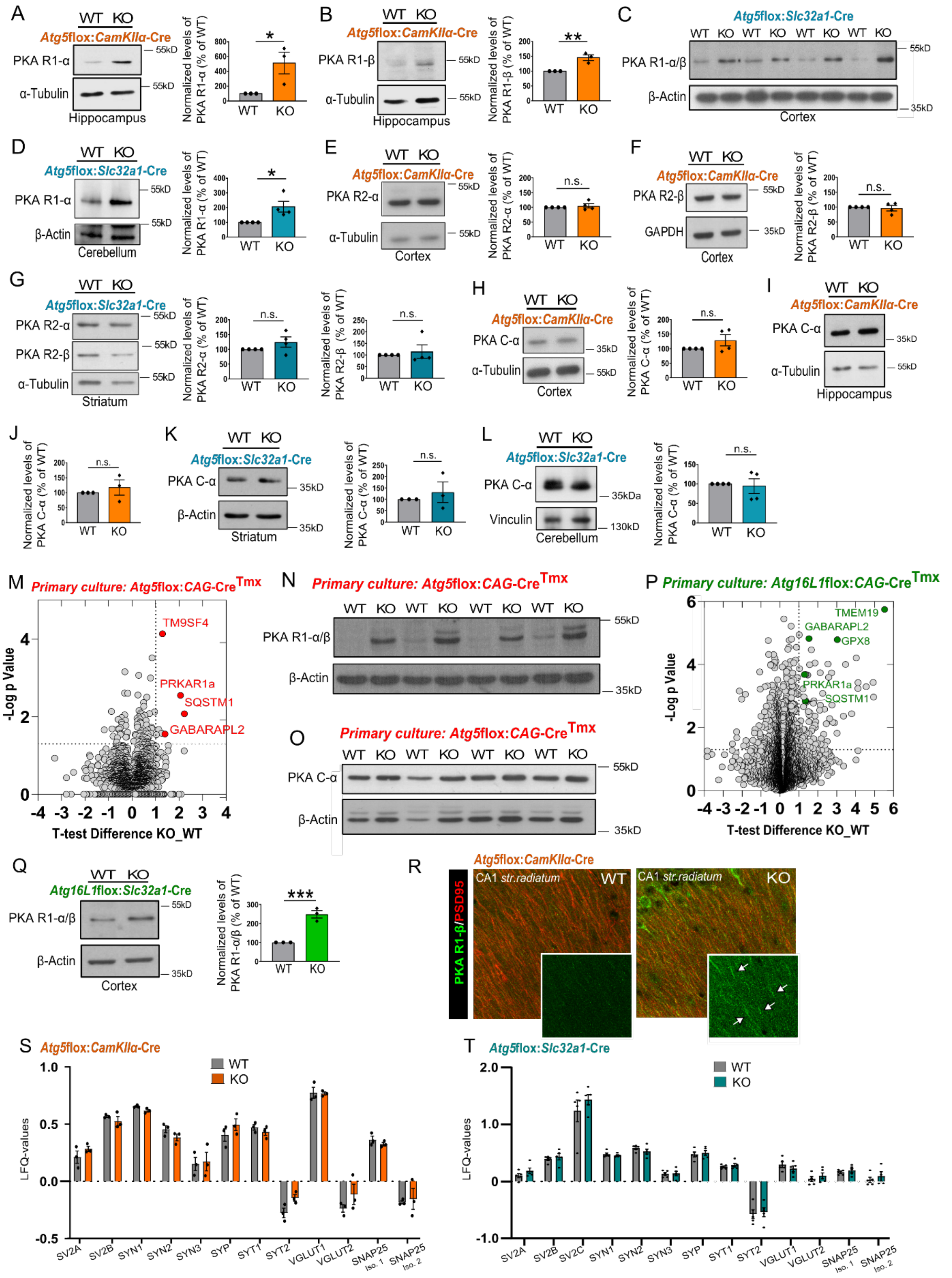
A
WT KO
Atg5flox:CamKIIα-Cre
PKA R1-α
α-Tubulin
Hippocampus
Normalized levels of PKA R1-α (% of WT)
B
Atg5flox:CamKIIα-Cre
PKA R1-β
Normalized levels of PKA R1-β (% of WT)
C
Atg5flox:Slc32a1-Cre
PKA R1-α/β
β-Actin
Cortex
D
Atg5flox:Slc32a1-Cre
Cerebellum
E
PKA R2-α
F
PKA R2-β
GAPDH
G
Striatum
H
PKA C-α
I
J
K
L
Vinculin
M
Primary culture: Atg5flox:CAG-Cre Tmx
TM9SF4
PRKAR1a
SQSTM1
GABARAPL2
-Log p Value
T-test Difference KO_WT
N
O
P
Primary culture: Atg16L1flox:CAG-Cre Tmx
TMEM19
GPX8
Q
Atg16L1flox:Slc32a1-Cre
Normalized levels of PKA R1-α/β (% of WT)
R
CA1 str.radiatum
PKA R1-β/PSD95
S
T
LFQ-values
SV2A
SV2B
SV2C
SYN1
SYN2
SYN3
SYP
SYT1
SYT2
VGLUT1
VGLUT2
SNAP25 Iso. 1
SNAP25 Iso. 2

**Supplementary figure 2 | Levels of PKA R1-α/β protein are upregulated in ATG5 and ATG16L1 KO neurons *in-vivo* and *in-vitro*.**

**(A)** Protein levels of PKA R1-α are significantly upregulated in hippocampal brain lysates of *Atg5*flox:*CamKIIα*-Cre KO mice compared to the WT set to 100% (KO: 513.1± 146.8%). p= 0.0241, n=3. Performed one-sample t-test. **(B)** Protein levels of PKA R1-β are significantly upregulated in hippocampal brain lysates of *Atg5*flox:*CamKIIα*-Cre KO mice compared to the WT set to 100% (WT set to 100%, KO: 145.3± 9.578%). p= 0.0046, n=3. Performed one-sample t-test. **(C)** Cortical lysates of *Atg5*flox:*Slc32a1*-Cre KO mice show increased PKA R1-α/β protein levels in compared to the WT. **(D)** Significantly upregulated protein level of PKA R1-α was detected in cerebellar lysates of *Atg5*flox:*Slc32a1*-Cre KO mice (WT set to 100%, KO: 206.6± 36.70%). p=0.0136, n=4. Performed one-sample t-test. **(E)** Protein levels of PKA R2-α are unaltered in cortical brain lysates of *Atg5*flox:*CamKIIα*-Cre KO mice compared to the WT set to 100% (KO: 105.3± 7.399%). n=4. **(F)** Protein levels of PKA R2-β are unaltered in cortical brain lysates of *Atg5*flox:*CamKIIα*-Cre KO mice compared to the WT set to 100% (KO: 96.40± 9.226%). n=4. **(G)** Protein levels of PKA R2-α/β are unaltered in striatal brain lysates of *Atg5*flox:*Slc32a1*-Cre KO mice (PKA R2-α KO: 124.6± 17.45%; PKA R2-β KO: 114.6± 28.94%). n=4. **(H)** No upregulation of PKA catalytic subunit α could be observed in *Atg5*flox:*CamKIIα*-Cre KO neurons compared to the WT, neither in cortical (H) (KO: 129.1±19.39%, p=0.0921, n=4) nor hippocampal (I) brain lysates (KO: 117.6± 25.45%, p=0.2637, n=3). **(K)** In striatal brain lysates of *Atg5*flox:*Slc32a1*-Cre mice the PKA C-α was not changed compared to the WT (KO: 131.2± 45.27%). p=0.2644, n=3. **(L)** The protein level of PKA C-α is unaltered in cerebellar brain lysates of *Atg5*flox:*Slc32a1*-Cre KO mice (KO: 94.36± 19.02%). p=0.3884, n=4. For analysis in (E-L), the KO was compared to the WT set to 100% and one-sample t-test was used for statistical analysis. **(M)** Volcano plot of differentially expressed proteins in *Atg5*flox:*CAG*-Cre$^{Tmx}$ primary cortical and hippocampal WT and KO neurons at DIV15-16 (n=3). **(N)** Western Blot analysis of PKA R1-α/β protein levels in *Atg5*flox:*CAG*-Cre$^{Tmx}$ KO cortico-hippocampal primary neurons at DIV 15-16. **(O)** Western Blot analysis of PKA C-α protein levels in *Atg5*flox:*CAG*-Cre$^{Tmx}$ primary cortical and hippocampal WT and KO neurons at DIV15-16 (n=4). **(P)** Volcano plot of differentially expressed proteins in *Atg16L1*flox:*CAG*-Cre$^{Tmx}$ primary cortical and hippocampal WT and KO neurons at DIV15-16 (n=3). **(Q)** Protein levels of PKA R1-α/β are upregulated in cortical brain lysates of *Atg16L1*flox:*Slc32a1*-Cre KO mice compared to the WT (WT set to 100%, KO: 248.4±

19.64%). p=0.0008, n=3. Performed one-sample t-test. **(R)** Uncropped confocal images of brain sections from *Atg5*flox:*CamKIIα*-Cre WT and KO CA1 region of the hippocampus, immunostained for PKA R1-β and PSD95 and used for analysis in Fig. 2K. **(S,T)** LFQ values of synaptic vesicle proteins from global proteome analysis of *Atg5*flox:*CamKIIα*-Cre (S) and *Atg5*flox:*Slc32a1*-Cre (T) WT and KO mice.

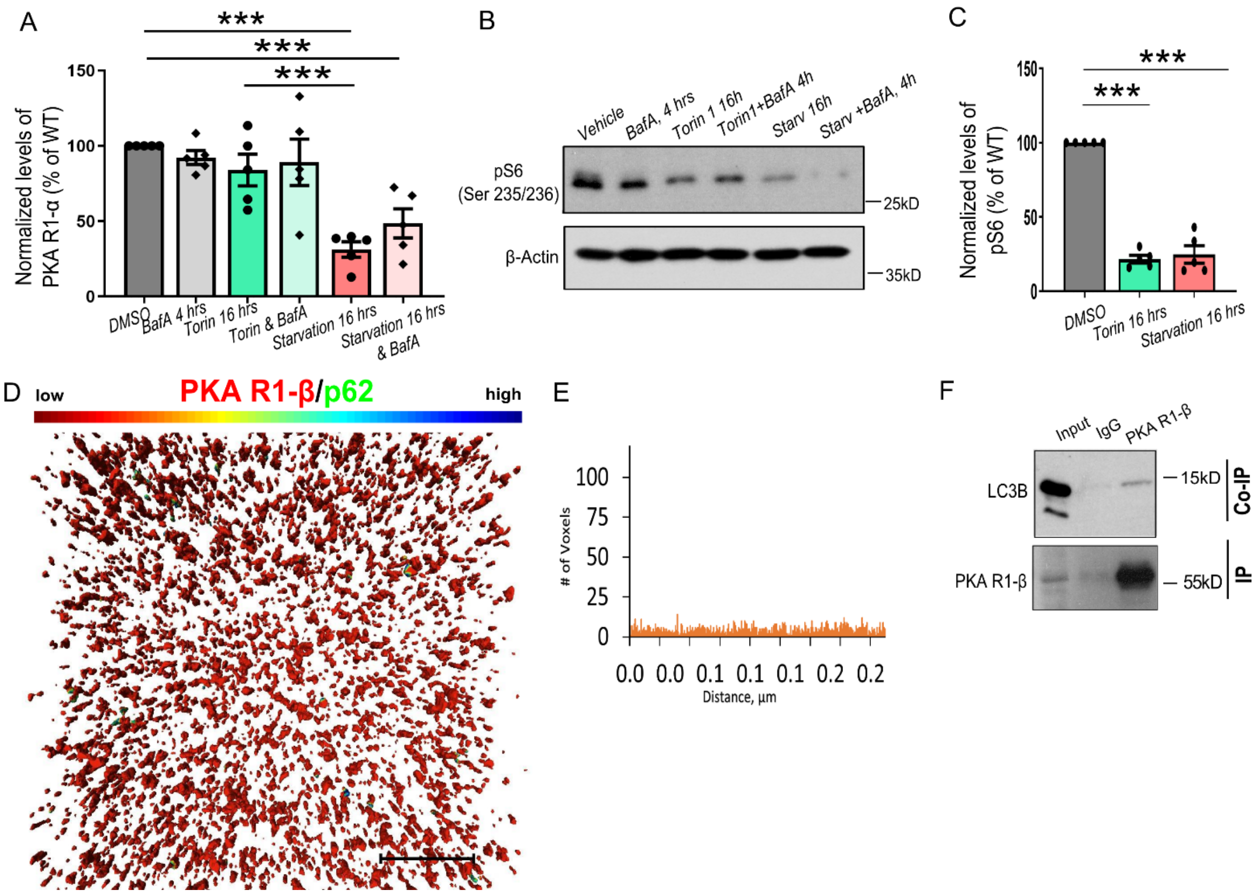


**Supplementary figure 3 |PKA R1 subunits degradation is controlled by starvation-induced LC3-mediated autophagy.**

**(A)** Amino acids and serum starvation (16 hours), but not the mTORC1 inhibition by the Torin 1 (250nM, 16 hours) induces a decrease in PKA R1-α total protein levels in NSC34 cells (DMSO: set to 100%, BafA 4 hours: 92.28± 4.612%, Torin 16 hours: 83.90± 10.48%, Torin & BafA: 89.15± 15.47%, Starvation 16 hours: 31.19± 5.085%, Starvation 16 hours & BafA: 48.52± 9.627%). $p_{DMSO/starvation\ 16\ hours}<0,0001$, $p_{Torin\ 16\ hours/starvation\ 16\ hours}=0.0007$, $p_{DMSO/starvation\ 16\ hours\ \&\ BafA}=0.0009$, n=5. Performed Two-Way ANOVA with Tukeys' multiple comparison test. **(B-C)** The levels of pS6 protein are equally reduced n NSC34 cells upon starvation and Torin 1 treatment (both 16h) (DMSO set to 100%, Torin 16 hours: 21.65± 2.541%, Starvation 16 hours: 24.82± 5.819%). $p_{DMSO/Torin\ 16\ hours}<0.0001$, $p_{DMSO/starvation\ 16\ hours}<0.0001$, n=5. Performed Two-Way ANOVA with Tukeys' multiple comparison test. **(D-E)** Uncropped image of the 3D-reconstruction shown in Fig. 3M (most right). Scale bar: 10 µm. The histogram in (E) shows the number of p62-labeled voxels (3D pixels) found within 250 nm of the PKA R1-β neuropil staining. **(F)** Co-immunoprecipitation of endogenous PKA R1-β with LC3 form *Atg5*flox:*CamKIIα*-Cre WT brain lysates. Input, 1.5% of the total lysate was added to the assay.

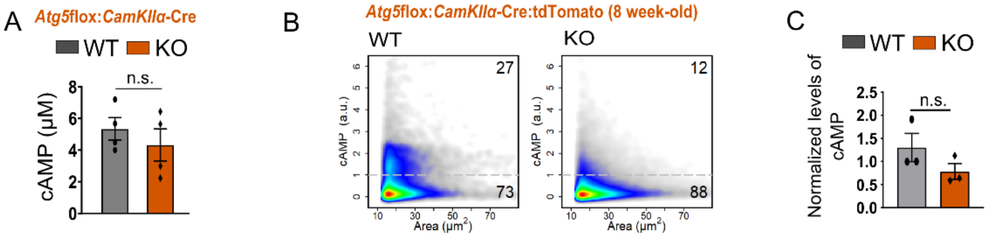


D

| log2 FoldChange | p-value | FDR | GENE ID | Description |
|---|---|---|---|---|
| 7.485700997 | 6.36852E-17 | 4.29614E-12 | *Gtf3c4* | general transcription factor IIIC, polypeptide 4 |
| 6.867994145 | 6.6644E-09 | 0.000149858 | *Srcap* | Snf2-related CREBBP activator protein |
| 6.625298107 | 3.12171E-07 | 0.00421175 | *Rev3l* | REV3 like, DNA directed polymerase zeta catalytic subunit |
| 6.60154495 | 4.4295E-08 | 0.000747025 | *Pdpk1* | 3-phosphoinositide dependent protein kinase 1 |
| 6.107961585 | 9.35384E-06 | 0.057363684 | *Brpf3* | bromodomain and PHD finger containing, 3 |
| 5.887685285 | 2.81595E-06 | 0.031660164 | *Adamtsl4* | ADAMTS-like 4 |
| 5.80575281 | 1.82371E-05 | 0.09463526 | *Dynlt1c* | dynein light chain Tctex-type 1C |
| 5.087358681 | 1.59584E-05 | 0.08971142 | *Eif4enif1* | eukaryotic translation initiation factor 4E nuclear import factor 1 |
| 2.267973449 | 3.78227E-12 | 1.27574E-07 | *9530068E07Rik* | RIKEN cDNA 9530068E07 gene |
| 1.89063498 | 8.09421E-06 | 0.054602732 | *Mylk* | myosin, light polypeptide kinase |
| 1.184307257 | 2.05084E-05 | 0.098819929 | *Sltm* | SAFB-like, transcription modulator |
| 1.169578801 | 7.58039E-06 | 0.054602732 | *Uba1* | ubiquitin-like modifier activating enzyme 1 |
| 1.125582966 | 4.30911E-06 | 0.036335998 | *Bahcc1* | BAH domain and coiled-coil containing 1 |
| -1.28013971 | 3.83611E-06 | 0.036335998 | *Cyld* | CYLD lysine 63 deubiquitinase |

**Supplementary figure 4 | Levels of cAMP are unaltered in ATG5 KO neurons.**

**(A)** Levels of cAMP were unchanged in hippocampal brain lysates of *Atg5*flox:*CamKIIα*-Cre KO mice compared to the WT (WT: 5.34±0.711, KO: 4.33±1.01). p=0.445, n=4. Performed Two-tailed unpaired t-test. **(B-C)** High-content screening microscopy analysis detects no alterations in cAMP fluorescence intensity in neurons isolated from 8-12-week-old *Atg5*flox:*CamKIIα*-Cre:tdTomato WT and KO mice (WT: 1.30±0.305, KO: 0.784±0.172). p= 0.213, n=3. Performed two-tailed unpaired t-test. **(D)** Differentially expressed genes (FDR<1) identified in mRNA sequencing analysis of 3-week- old *Atg5*flox:*CamKIIα*-Cre:tdTomato FACS sorted WT and KO neurons (Sleuth-based analysis of Kalisto transcriptome output, see also Suppl. Table 2).

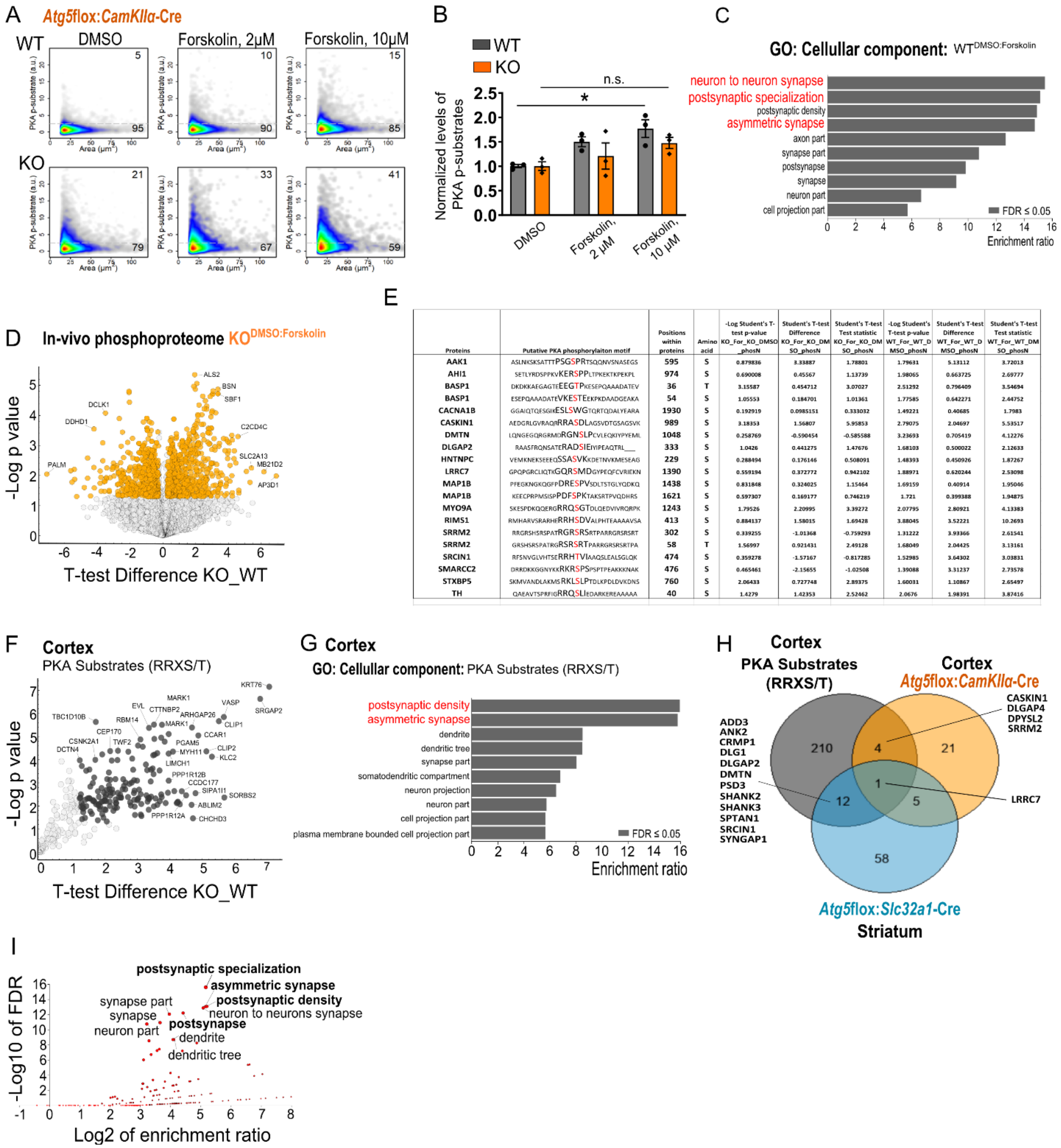


| Proteins | Putative PKA phosphorylaiton motif | Positions within proteins | Amino acid | -Log Student's T-test p-value KO_For_KO_DMSO_phosN | Student's T-test Difference KO_For_KO_DMSO_phosN | Student's T-test Test statistic KO_For_KO_DMSO_phosN | -Log Student's T-test p-value WT_For_WT_DMSO_phosN | Student's T-test Difference WT_For_WT_DMSO_phosN | Student's T-test Test statistic WT_For_WT_DMSO_phosN |
|---|---|---|---|---|---|---|---|---|---|
| AAK1 | ASLNKSKSATTTPSGSPRTSQQNVSNASEGS | 595 | S | 0.879836 | 3.33887 | 1.78801 | 1.79631 | 5.13112 | 3.72013 |
| AHI1 | SETLYRDSPPKVKERSPPLTPKEKTKPEKPL | 974 | S | 0.690008 | 0.45567 | 1.13739 | 1.98065 | 0.663725 | 2.69777 |
| BASP1 | DKDKKAEGAGTEEEGTPKESEPQAAADATEV | 36 | T | 3.15587 | 0.454712 | 3.07027 | 2.51292 | 0.796409 | 3.54694 |
| BASP1 | ESEPQAAADATEVKESTEEKPKDAADGEAKA | 54 | S | 1.05553 | 0.184701 | 1.01361 | 1.77585 | 0.642271 | 2.44752 |
| CACNA1B | GGAIQTQESGIKESLSWGTQRTQDALYEARA | 1930 | S | 0.192919 | 0.0985151 | 0.333032 | 1.49221 | 0.40685 | 1.7983 |
| CASKIN1 | AEDGRLGVRAQRRRASDLAGSVDTGSAGSVK | 989 | S | 3.18353 | 1.56807 | 5.95853 | 2.79075 | 2.04697 | 5.53517 |
| DMTN | LQNGEGQRGRMDRGNSLPCVLEQKIYPYEML | 1048 | S | 0.258769 | -0.590454 | -0.585588 | 3.23693 | 0.705419 | 4.12276 |
| DLGAP2 | RAASFRQNSATERADSIEIYIPEAQTRL___ | 333 | S | 1.0426 | 0.441275 | 1.47676 | 1.68103 | 0.500022 | 2.12633 |
| HNTNPC | VEMKNEKSEEEQSSASVKKDETNVKMESEAG | 229 | S | 0.288494 | 0.176146 | 0.508091 | 1.48393 | 0.450926 | 1.87267 |
| LRRC7 | GPQPGRCLIQTKGQRSMDGYPEQFCVRIEKN | 1390 | S | 0.559194 | 0.372772 | 0.942102 | 1.88971 | 0.620244 | 2.53098 |
| MAP1B | PFEGKNGKQGFPDRESPVSDLTSTGLYQDKQ | 1438 | S | 0.831848 | 0.324025 | 1.15464 | 1.69159 | 0.40914 | 1.95046 |
| MAP1B | KEECPRPMSISPPDFSPKTAKSRTPVQDHRS | 1621 | S | 0.597307 | 0.169177 | 0.746219 | 1.721 | 0.399388 | 1.94875 |
| MYO9A | SKESPDKQQERGRRQSGTDLQEDVIVRQRPK | 1243 | S | 1.79526 | 2.20995 | 3.39272 | 2.07795 | 2.80921 | 4.13383 |
| RIMS1 | RMHARVSRARHERRHSDVALPHTEAAAAVSA | 413 | S | 0.884137 | 1.58015 | 1.69428 | 3.88045 | 3.52221 | 10.2693 |
| SRRM2 | RRGRSHSRSPATRGRSRSRTPARRGRSRSRT | 302 | S | 0.339255 | -1.01368 | -0.759293 | 1.31222 | 3.93366 | 2.61541 |
| SRRM2 | GRSHSRSPATRGRSRSRTPARRGRSRSRTPA | 58 | T | 1.56997 | 0.921431 | 2.49128 | 1.68049 | 2.04425 | 3.13161 |
| SRCIN1 | RFSNVGLVHTSERRHTVIAAQSLEALSGLQK | 474 | S | 0.359278 | -1.57167 | -0.817285 | 1.52985 | 3.64302 | 3.03831 |
| SMARCC2 | DRRDKKGGNYKKRKRSPSPSPTPEAKKKNAK | 476 | S | 0.465461 | -2.15655 | -1.02508 | 1.39088 | 3.31237 | 2.73578 |
| STXBP5 | SKMVANDLAKMSRKLSLPTDLKPDLDVKDNS | 760 | S | 2.06433 | 0.727748 | 2.89375 | 1.60031 | 1.10867 | 2.65497 |
| TH | QAEAVTSPRFIGRRQSLIEDARKEREAAAAA | 40 | S | 1.4279 | 1.42353 | 2.52462 | 2.0676 | 1.98391 | 3.87416 |

**Supplementary figure 5 |Neuronal PKA targets are enriched at the postsynaptic density of excitatory synapses.**

**(A,B)** High-content screening microscopy analysis of cortical neurons isolated from 8-10-week-old *Atg5*flox:*CamKIIα*-Cre:tdTomato mice and immunostained with RRXS/T motif antibody to visualize phosphorylated PKA substrates. Forskolin stimulation increased levels of RRXS/T motif-carrying proteins in WT, but not in KO neurons ($WT_{DMSO}$: 1.005±0.034, $KO_{DMSO}$:

1.004±0.086, $WT_{Forskolin\ 2\mu M}$: 1.502±0.098, $KO_{Forskolin\ 2\mu M}$: 1.209±0.269, $WT_{Forskolin\ 10\mu M}$: 1.770±0.182, $KO_{Forskolin\ 10\mu M}$: 1.476±0.116). $p_{WT\ DMSO/WT\ Forskolin\ 10\mu M}$= 0.034, n=3. Performed Two-Way ANOVA with Tukeys' multiple comparison test. **(C)** WebGestalt-based gene ontology (GO) analysis of "cellular component"-enriched terms in the phosphoproteome of Forskolin-treated WT brain slices. **(D)** Volcano plot of differentially expressed phosphopeptides in acute Atg5flox:*CamKIIα*-Cre KO brain slices treated either with DMSO or 50µM Forskolin for 15 min. Highlighted are the proteins with $p<0.05$ and a 1.2/-1.2 minimal fold-change (log2FC ±0.58). n=5 for each genotype. **(E)** Table overview of putative PKA targets downregulated in ATG5 KO neurons in-vivo, which are shown in Fig. 5G. **(F)** Volcano plot of co-immunoprecipitated PKA substrates in WT brain using the RRXS/T motif antibody. Highlighted are significant proteins ($p<0.05$) with a FC>1.2. n=4. **(G)** WebGestalt-based gene ontology (GO) analysis of "cellular component"-enriched terms in all proteins highlighted in (F). **(H)** Venn diagram showing the number of putative PKA substrates detected using the co-immunoprecipitation analysis in (F) and phosphopeptides significantly downregulated in *Atg5*flox:*CamKIIα*-Cre KO and *Atg5*flox:*Slc32a1*-Cre KO brains (cut-off $p<0.05$, Fold change <-1.2). **(I)** WebGestalt- based gene ontology (GO) analysis of "cellular component" enriched terms among common proteins identified (H).

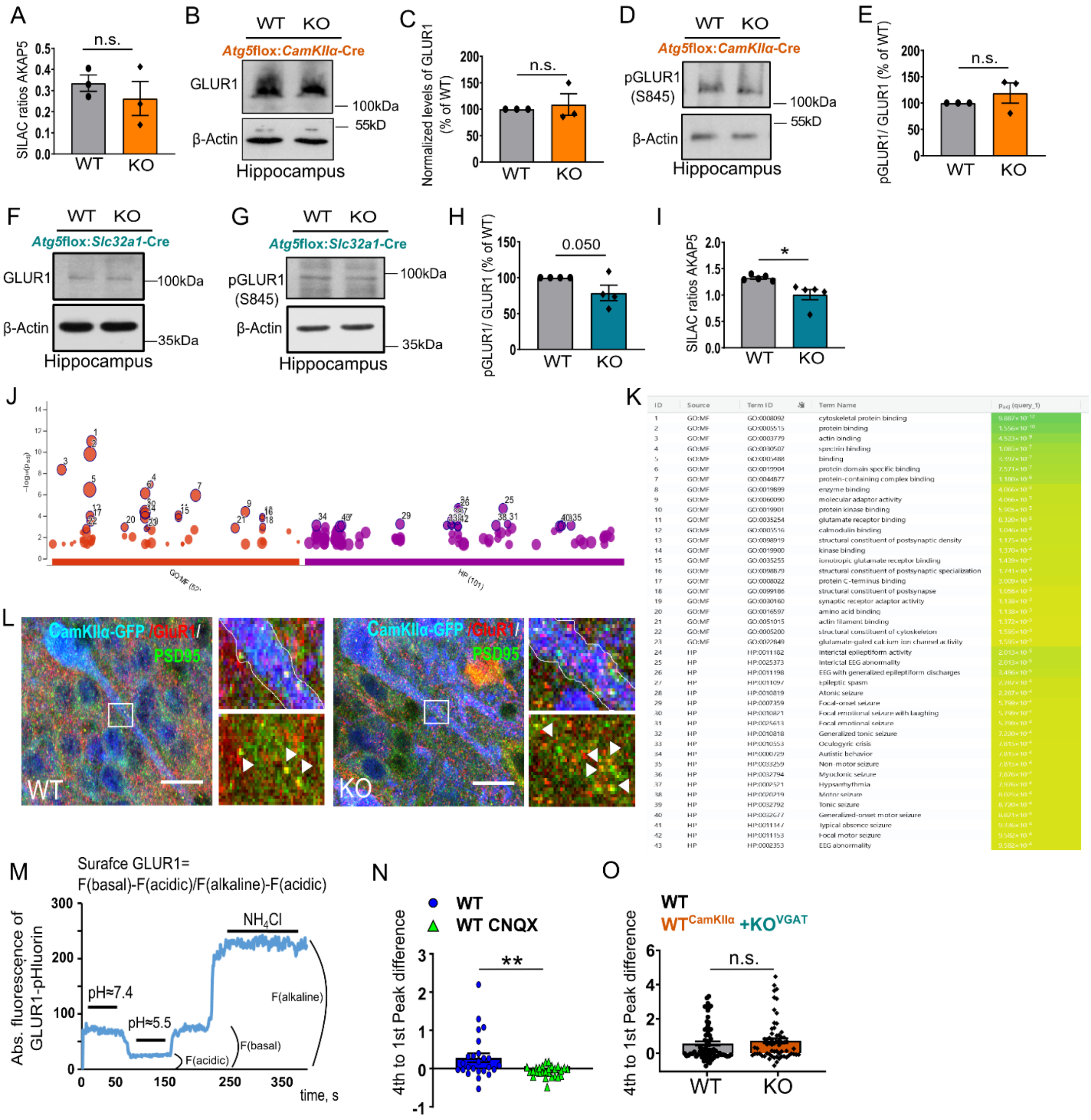

**Supplementary figure 6 | GLUR1 total levels or its phosphorylation state of GLUR1 are not altered in ATG5 KO neurons.**

**(A)** SILAC-based proteome analysis does not reveal changes in AKAP5 (AKAP79) protein levels in *Atg5*flox:*CamKIIα*-Cre KO mice comparing to the WT (SILAC ratios, WT: 0.336±0.039, KO: 0.262±0.0805). p=0.458, n=3. Performed two-tailed unpaired t-test. **(B,C)** Total protein levels of GLUR1 are unaltered in *Atg5*flox:*CamKIIα*-Cre hippocampal lysates of KO mice (WT set to 100%, KO: 108.9±20.47%). p=0.343, n=3. Performed one-sample t-test. **(D,E)** Protein levels of phosphorylated GLUR1, normalized to total GLUR1 protein level, are unaltered in

*Atg5*flox:*CamKIIα*-Cre hippocampal KO brain lysates compared to the WT (WT set to 100%, KO: 119.1±19.31%). p=0.189, n=3. Performed one-sample t-test. **(F)** Total protein levels of GLUR1 are unaltered in *Atg5*flox:*Slc32a1*-Cre hippocampal KO lysates compared to the WT (WT set to 100%, KO: 92.74±17.48). p=0.3462, n=4. Performed one-sample t-test. **(G,H)** Protein levels of phosphorylated GLUR1, normalized to total GLUR1 protein level, are slightly reduced in hippocampal brain lysates of *Atg5*flox:*Slc32a1*-Cre KO mice compared to WT (WT set to 100%, KO: 78.86±10.88). p=0.050, n=4. Performed one-sample t-test. **(I)** SILAC-based proteome analysis identifies alterations in AKAP5 (AKAP79) protein levels in *Atg5*flox:*Slc32a1*-Cre KO brain lysates compared to the (SILAC ratios, WT: 1.325±0.023, KO: 1.008±0.096). p=0.013, n=5. Performed two-tailed unpaired t-test. **(J, K)** Raw data from gene ontology cellular component analysis using g:Profiler. Modified version is shown in Fig. 6A. **(L)** Confocal images of immunostained brain sections from 13-week-old *Atg5*flox:*CamKIIα*-Cre WT and KO mice, which were stereotactically injected with *CamKIIα*-eGFP to visualize the excitatory neurons. Images reveal increased colocalization of GLUR1 and PSD95 in dendrites of KO neurons. **(M)** Representative example of GLUR1 pHluorin fluorescence at the dendrite upon exposure to buffer containing 25 mM MES at pH 5.5 and 50 mM $NH_4Cl$ at pH 7.4. Exposure to pH 5.5 quenches GLUR1 pHluorin fluorescence on the dendrite surface (plasma membrane), and application of $NH_4Cl$ raises the intravesicular pH to 7.4, thereby dequenching vesicular GLUR1 pHluorin fluorescence. **(N)** Analysis of $4^{th}$ to $1^{st}$ peak difference of WT cells expressing $GCAMP7^{CamKII\alpha}$ treated with DMSO or CNQX show a decreased calcium response to electrical field stimulation (WT: 0.282± 0.117, KO: -0.070± 0.027). p=0.004, $n_{WT}$=25, $n_{WT\ CNQX}$=27. Performed two-tailed unpaired t-test. **(O)** Analysis of $4^{th}$ to $1^{st}$ peak difference of $GCAMP7^{CamKII\alpha}$ expressing neurons in a network with *Atg5*flox:*Slc32a1*-Cre KO neurons does not reveal changes in neuronal excitation (WT: 0.561± 0.125, KO: 0.721± 0.147). p= 0.406, $n_{WT}$=69, $n_{KO}$=64. Performed two-tailed unpaired t-test.

**Table S1:** Significantly ($p<0.05$) dysregulated proteome hits from of *Atg5*flox:*CamKIIα*-Cre, *Atg5*flox:*Slc32a1*-Cre, *Atg5*flox:*CamKIIα*-Cre/tdTomato and *Atg5*flox:*Slc32a1*-Cre/tdTomato WT and KO mice/ neurons shown in Fig. 1K-N. This table also contains detailed information about the GO cellular component analysis shown in Figure 1O and fig. S1S.

**Table S2:** The RNA sequencing data from the *Atg5*flox:*Slc32a1*-Cre/tdTomato FACS-sorted neurons. The table contains the Kallisto transcriptome, the heatmap shown in Fig. 4G, the sleuth-based DGE analysis and the ChEA TF enrichment analysis shown in Fig. 4H,I, the data represented by the heatmap in Fig. 4J, as well as the differentially expressed genes shown in Fig. S4D.

**Table S3:** The table contains the significantly ($p<0.05$) downregulated ($FC<-1.2$) hits of the global phosphoproteome of *Atg5*flox:*CamKIIα*-Cre/tdTomato and *Atg5*flox:*Slc32a1*-Cre/tdTomato WT and KO mice/ neurons. Part of the data is shown in Fig. 5C and the list of hits is used for analyses in Fig. 5 and Fig. 6.

**Table S4:** The table contains the data shown in Fig. 5 and Fig. 6. It contains the raw data of the phosphoproteome analysis of acute *Atg5*flox:*CamKIIα*-Cre/tdTomato brain slices treated with Forskolin and DMSO as control, and the data presented in Fig. 5E-J and Fig. 6A,B.

**Table S5:** Primers used for genotyping of genetically modified mice used in the current study.

| **Gene** | **Sequence (5’- 3’)** |
|---|---|
| ***Atg5*** | |
| *Atg5* forward 1 | GAA TAT GAA GCC ACA CCC CTG AAA TG |
| *Atg5* forward 2 | ACA ACG TCG AGC ACG CTG GCG AAG G |
| *Atg5* reverse | GTA CTG CAT AAT GGT TTA ACT CTT GC |
| ***Atg16L1*** | |
| *Atg16L1_1* | CAG AAT AAT TTC CGG CAG AGA CCG G |
| *Atg16L1_2* | AGC CAA AGA AGG AAG GTA AGC AAC GAA |
| ***Cre*** | |
| *Cre_1* | GAA CCT GAT GGA CAT GTT CAG G |
| *Cre_2* | AGT GCG TTC GAA CGC TAG AGC CTG T |
| *Cre_3* | TTA CGT CCA TCG TGG ACA |
| *Cre_4* | TGG GCT GGG TGT TAG CC |
| ***tdTomato*** | |

| | |
|---|---|
| *tdTomato_1* | AAG GGA GCT GCA GTG GAG TA |
| *tdTomato_2* | CCG AAA ATC TGT GGG AAG TC |
| *tdTomato_3* | GGC ATT AAA GCA GCG TAT CC |
| *tdTomato_4* | CTG TTC CTG TAC GGC ATG G |

**Table S6:** Plasmid DNA used for transfection of cells *in-vitro* in the current study.

| **Plasmid (source gene)** | **Manufacturer** | **Identifier** |
|---|---|---|
| GFP | Clontech | pT3027.5 |
| pEGFP-C1-hAPG5 (ATG5-GFP) | addgene | #22952 |
| Prkar1a shRNA | OriGene | TL501735 |
| Prkar1a shRNA | OriGene | TL501736 |
| pcDNA3-mouse PKA-RIalpha-mEGFP | addgene | #45525 |
| pcDNA3-mouse PKA-RIbeta-mEGFP | addgene | #45526 |
| pCMV-SEP-GluA1 | addgene | #64942 |

**Table S7:** AAVs used for transfection of primary neurons in the current study.

| **Name** | **Full description** | **Manufacturer** | **Identifier** |
|---|---|---|---|
| eGFP $^{CamKII\alpha}$ | AAV9.CamKIIα0.4.eGFP.WPRE.rBG | Penn Vector Core | 105541 |
| eGFP-Cre $^{CamKII\alpha}$ | AAV9.CamKIIα.HI.eGFP-Cre.WPRESV40 | Penn Vector Core | 105551 |
| Cre $^{GAD67}$ | ssAAV-5/2-hGAD67-chl-iCre-SV40p(A) | Viral Vector Facility VVF | V197-5 |
| Cre $^{CamKII\alpha}$ | ssAAV-9/2-mCaMKIIα-iCre-WPRE-hGHp(A) | Viral Vector Facility VVF | V206-9 |
| GCAMP7f $^{CamKII\alpha}$ | ssAVV-9/2-mCaMKIIα-jGCaMP7f-WPREbGHp(A) | Viral Vector Facility VVF | V493-9 |

**Table S8:** Antibodies & DNA labelling dyes and their used concentration in Immunoblotting (WB), Immunocytochemistry (ICC), Immunohistochemistry (IHC) and Immunoprecipitation (IP) analysis.

| **Antibodies** | **Used concentration** | | | | **Manufacturer** | **Catalog number** |
|---|---|---|---|---|---|---|
| **Target protein** | WB | ICC | IHC | IP | | |
| ATG5 | 1:1000 | - | - | - | Abcam | ab108327 |

| | | | | | | |
|---|---|---|---|---|---|---|
| β-Actin | 1:1000 | - | - | - | Sigma | A-5441 |
| Bassoon | - | - | 1:500 | - | Synaptic Systems | 141002 & 141004 |
| CREB | 1:1000 | - | - | - | Cell Signaling | 9197 |
| cAMP | - | 1:500 | - | - | Abcam | ab24856 |
| GAPDH | 1:1000 | - | - | - | Sigma-Aldrich | G8795 |
| GFP | - | 1:2000 | 1:1000 | - | Abcam | ab13970 |
| GLUA1 | 1:1000 | 1:500 | 1:500 | - | Synaptic Systems | 182011 |
| Glutamate receptor 1 (AMPA subtype) phospho S845 | 1:1000 | - | - | - | Abcam | ab76321 |
| LC3B | 1:2000 | - | - | - | Novus Biologicals | NB600-1384SS |
| MAP2 | - | 1:500 | - | - | Sigma-Aldrich | M9942 |
| NBR1 | - | - | 1:300 | - | Santa Cruz Biotechnology | sc-130380 |
| Normal Rabbit IgG | - | - | - | 2 µg | Cell Signaling | 2729 |
| Normal Mouse IgG | - | - | - | 2 µg | Millipore | 12-371 |
| Normal Sheep IgG | - | - | - | 2 µg | Jackson Immuno Research | 713-005-147 |
| p62 | 1:1000 | 1:1000 | 1:1000 | - | Progen | GP62-C |
| pCREB (Ser133) | 1:1000 | 1:500 | - | - | Cell Signaling | 9198 |
| pCREB (S133) | - | 1:500 | - | - | Thermo Fisher (Pierce) | MA5-11192 |
| pGluR1 (S845) | 1:1000 | - | - | - | Abcam | ab76321 |
| pPKA-R2-α | - | 1:500 | - | - | Abcam | ab32390 |
| pPKA-Substrate | 1:1000 | - | - | 2 µg | Cell Signaling | 9624 |
| PKA C-α | 1:1000 | - | - | - | Cell Signaling | 5842 |

| | | | | | | |
|---|---|---|---|---|---|---|
| PKA R1-α | 1:1000 | 1:300 | 1:300 | 2 µg | Cell Signaling | 5675 |
| PKA R1-α/β | 1:1000 | - | - | - | Cell Signaling | 3927 |
| PKA R1-β | 1:1000 | 1:300 | 1:300 | 2 µg | R&D Systems | AF4177 |
| PKA R2-α | 1:1000 | - | - | - | BD Transduction Laboratories | 612242 |
| PKA R2-β | 1:1000 | - | - | - | Abcam | Ab32390 |
| PSD95 | - | 1:300 | 1:300 | - | Synaptic Systems | 124 008 |
| S6 Ribosomal Protein (phosho Ser235/236) | 1:1000 | - | - | - | Cell Signaling | 2211 |
| α-Tubulin | 1:1000 | - | - | - | Synaptic Systems | 302 211 |
| Vinculin [EPR8185] | 1:1000 | - | - | - | Abcam | ab129002 |
| **DNA labelling dyes** | | | | | | |
| DAPI | - | 1:10000 | - | - | Roth | 6335.1 |
| DRAQ5 | - | 1:1000 | - | - | Thermo Scientific | 62254 |

**Table S9:** Secondary antibodies with HRP (WB) or fluorescence (ICC / IHC) tag and their used concentration.

| **Antibody target** | **Used concentration** | | | **Manufacturer** | **Catalog number** |
|---|---|---|---|---|---|
| | WB | ICC | IHC | | |
| Goat anti-Mouse IgG (H+L) peroxidase-conjugated | 1:5000 | - | - | Jackson Immuno Research | 115-035-003 |
| Goat anti-Rabbit IgG (H+L) peroxidase-conjugated | 1:5000 | - | - | Jackson Immuno Research | 111-035-003 |
| Goat anti-Guinea Pig IgG (H+L) peroxidase-conjugated | 1:5000 | - | - | Jackson Immuno Research | 106-035-003 |
| Goat anti-Sheep IgG (H+L) peroxidase-conjugated | 1:5000 | - | - | Invitrogen | A16041 |
| Alexa Fluor 488 Goat anti-Mouse IgG | - | 1:500 | - | Life Technologies GmbH | A11029 |
| Alexa Fluor 488 Goat anti-Rabbit IgG | - | 1:500 | 1:500 | Life Technologies GmbH | A11034 |
| Alexa Fluor 488 Goat anti-Chicken IgG | - | 1:500 | 1:500 | Life Technologies GmbH | A11039 |
| Alexa Fluor 488 Goat anti-Guinea Pig IgG | - | 1:500 | 1:500 | Life Technologies GmbH | A11073 |
| Alexa Fluor 488 Donkey anti-Sheep IgG | | 1:500 | 1:500 | Life Technologies GmbH | A11015 |
| Alexa Fluor 568 Goat anti-Mouse IgG | - | 1:500 | 1:500 | Life Technologies GmbH | A-11031 |
| Alexa Fluor 568 Goat anti-Rabbit IgG | - | 1:500 | 1:500 | Life Technologies GmbH | A11036 |
| Alexa Fluor 568 Goat anti-Guinea Pig IgG | - | 1:500 | 1:500 | Life Technologies GmbH | A11075 |

| | | | | | |
|---|---|---|---|---|---|
| Alexa Fluor 568 donkey anti-mouse IgG | - | 1:500 | 1:500 | Life Technologies GmbH | A10037 |
| Alexa Fluor 555 Donkey anti-Sheep IgG | - | 1:500 | 1:500 | Life Technologies GmbH | A21436 |
| Alexa Fluor 647 Goat anti-Mouse IgG | - | 1:500 | 1:500 | Life Technologies GmbH | A21236 |
| Alexa Fluor 647 Goat anti-Mouse IgG | - | 1:500 | - | Life Technologies GmbH | A327728 |
| Alexa Fluor 647 Goat anti-Rabbit IgG | - | 1:500 | 1:500 | Life Technologies GmbH | A21245 |
| Alexa Fluor 647 Goat anti-Guinea Pig IgG | - | 1:500 | 1:500 | Life Technologies GmbH | A21450 |
| Alexa Fluor 647 Donkey anti-Rabbit IgG | - | 1:500 | 1:500 | Life Technologies GmbH | A31573 |

**Movie S1:** *Atg5*flox:*CamKIIα*-Cre$^{tg}$ KO mice example video showing an ATG5 KO mice having a spontaneous recurrent seizure.

**Data S1**: Examples of physiological ECoG sleep patterns during transitions from slow-wave sleep (SWS) to REM sleep in wildtype and *Atg5*flox:*CamKII*α-Cre mice.
A) Representative ECoG trace and corresponding multitaper spectrogram (top two panels) during the transition from SWS to REM sleep in a wild-type mouse obtained during chronic radiotelemetry recordings. A zoomed-in trace and corresponding wavelet spectrogram of the REM period indicated by red dotted lines in the multitaper spectrogram are shown in the bottom two panels. B) Representative ECoG trace during an SWS-REM transition recorded from an *Atg5*flox:*CamKII*α-Cre mutant mouse and corresponding spectrograms as in (A) indicating the presence of physiological sleep ECoG patterns in *Atg5*flox:*CamKII*α-Cre mutants that are comparable to wildtype controls.

**Data S2:** ImageJ-macro for colocalization analyses shown in Fig. 6E-F
