## Supplementary figures and images for "Autophagy regulates neuronal excitability by controlling cAMP/Protein Kinase A signaling"

### Data S1

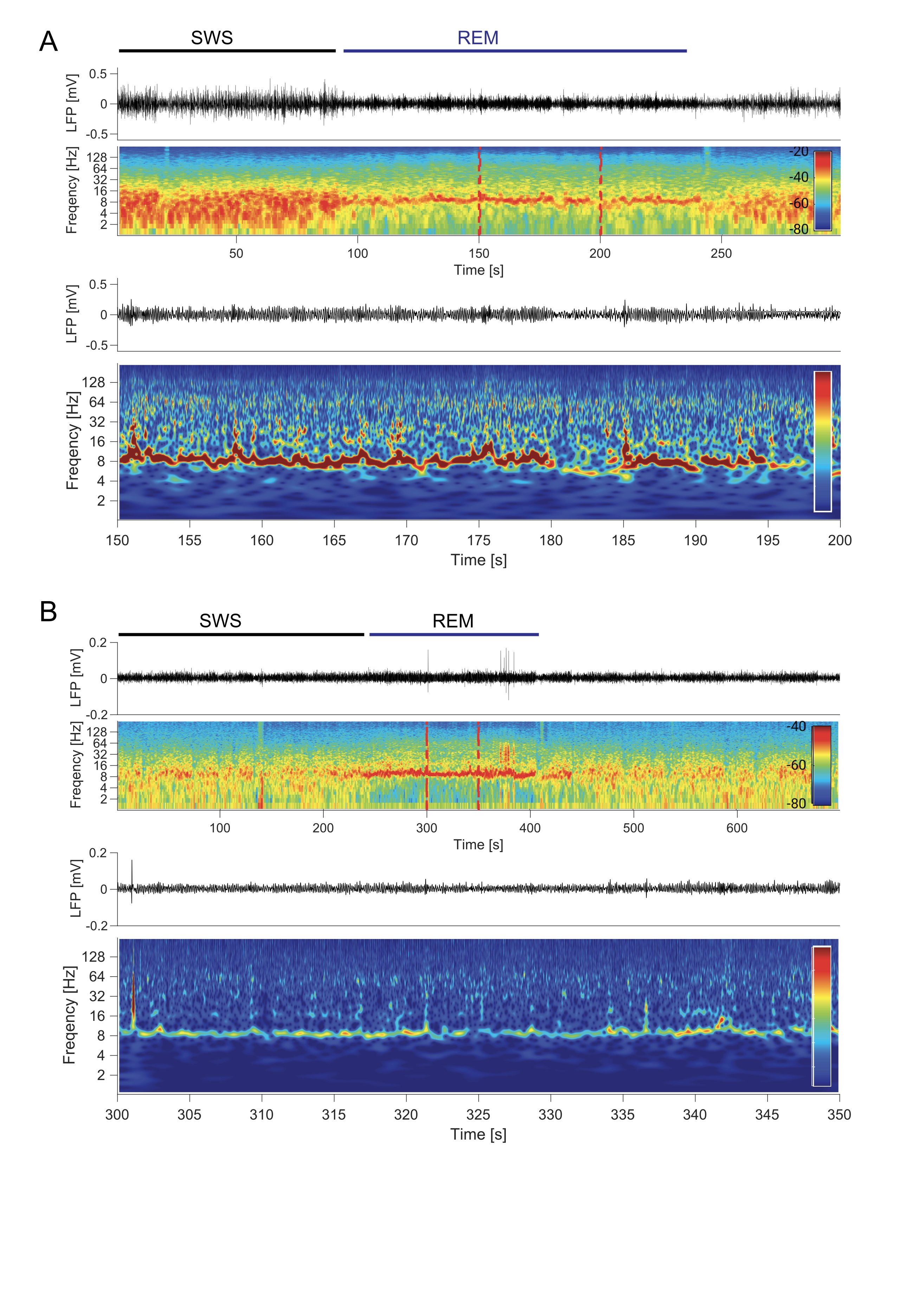
